# Spatial mapping of dextran sodium sulphate-induced intestinal inflammation and its systemic effects

**DOI:** 10.1101/2024.04.26.591292

**Authors:** Lauren Adams, Orhan Rasid, Heather Hulme, Tezz Quon, Richard Burchmore, Simon Milling, Richard J.A. Goodwin, Daniel M. Wall

**Affiliations:** School of Infection and Immunology, College of Medical, Veterinary and Life Sciences, Sir Graeme Davies Building, University of Glasgow, Glasgow G12 8TA, United Kingdom; Imaging and Data Analytics, Clinical Pharmacology and Safety Sciences, Biopharmaceuticals R&D, AstraZeneca, Cambridge CB4 0WG, United Kingdom

**Keywords:** Spatial biology, inflammation, colitis, microbiome, imaging

## Abstract

Inflammatory bowel disease (IBD) is a multifactorial disease and patients frequently experience extraintestinal manifestations affecting multiple sites. Causes of systemic inflammation remain poorly understood but molecules originating from the intestine likely play a role, with microbial and host small molecules polarizing host immune cells towards a pro- or anti-inflammatory phenotype. Using the dextran sodium sulphate (DSS) mouse model, which mimics the disrupted barrier function, microbial dysbiosis and immune cell dysregulation of IBD, we investigated metabolomic and phenotypic changes at intestinal and systemic sites. Using spatial biology approaches we mapped distribution and relative abundance of molecules and cell types across a range of tissues revealing significant changes in DSS-treated mice. Molecules identified as contributing to the statistical separation of treated from control mice were spatially localized within organs to determine their effects on cellular phenotypes through imaging mass cytometry. This spatial approach identified both intestinal and systemic molecular drivers of inflammation, including several not previously implicated in inflammation linked to IBD or the systemic effects of intestinal inflammation. Metabolic and inflammatory pathway interplay underpins systemic disease and determining drivers at the molecular level may aid the development of new targeted therapies.

## Introduction

Inflammatory bowel disease (IBD) is a complex multifactorial disease characterised by chronic relapsing-remitting inflammation. Many IBD patients experience extraintestinal manifestations (EIM) of disease that can affect nearly any organ system including dermatological, hepatopancreatobiliary, ocular, renal, neural and pulmonary systems (1). Complications such as non-alcoholic fatty liver disease (NAFLD) and gallstones have also been associated with disease and linked to metabolic dysfunction (2). The understanding of the inflammatory processes that underpin IBD onset and progression have advanced due to murine models of intestinal colitis. These models have enabled characterisation of the complexity of IBD pathogenesis by reflecting underlying molecular mechanisms of IBD such as genetic mutations, impaired barrier function, and dysfunctional innate and adaptive immune responses. Experimental models, such as the most common chemically induced colitis model using dextran sodium sulphate (DSS), are quick, reproducible, and controllable (3). DSS is a sulphate polysaccharide that acts as a colitogen *in vivo* with anticoagulant properties inducing similar clinical symptoms and histological features to those of human IBD. This model similarly exhibits EIMs including impaired brain function, inflammation of the liver and lungs, splenomegaly and immune cell dysfunction in the spleen and metabolic changes in the kidney (4, 5).

DSS-induced inflammation mostly affects, but is not limited to, the colonic mucosa as the negatively charged sulphate groups are thought to damage the intestinal mucosa and epithelial monolayer lining (6). The damage to the epithelium involves loss of crypts and an altered immune response characterised by infiltration of granulocytes leading to increasing gut permeability enabling commensal gut microorganisms to penetrate through the dysfunctional epithelial tight junctions into the lamina propria (6, 7). This influx of microbiota in IBD can result in the activation of intestinal macrophages which secrete high levels of inflammatory cytokines such as tumour necrosis factor alpha (TNF-α), interleukin-1 beta (IL-1β), and IL-6 (8). These cytokines act as chemoattractants to recruit immune cells such as dendritic cells, T cells, B cells and neutrophils into the colon which perpetuates inflammation (9).

DSS-induced colitis in mice has highlighted that intestinal bacteria are essential for the development of inflammation that is representative of human IBD (7, 10). Studies using germ-free (GF) mice showed that DSS induced only minimal inflammation, with no colonic thickening or shortening and low levels of proinflammatory cytokines, in spite of mice having impaired barrier function (10). Conventionally colonized mice, treated with antibiotics to perturb the intestinal microbiota, had an increased inflammatory response but were less prone to epithelial injury. This was attributed to an increase in IL-10 and increases in other barrier preservation related markers (10). Furthermore, longitudinal microbiome analysis studies have indicated that DSS colitis results in dynamic fluctuations in microbial diversity, transitioning from a normal state to dysbiosis (11). Whether gut microbial dysbiosis is a cause or a consequence of DSS colitis is inconclusive but the evidence suggests colitis involves simultaneous interactions between the epithelial barrier, intestinal microbiota, and host immune system (11).

Intestinal microbial dysbiosis during DSS colitis can reduce specific immune regulatory metabolites promoting inflammation (12–14). Ursodeoxycholic acid, tryptophan and derivatives promote barrier integrity and reduce pro-inflammatory cytokine release (13, 15). Multi-omic approaches have provided a better understanding of the molecular basis of IBD pathogenesis and similar application of these technologies to DSS colitis has highlighted significant alteration of the metabolome and specific pathways such as amino acid and lipid metabolism which are upregulated as severity of inflammation increases (16, 17).

Here we used the DSS colitis model to further study the effects of microbial and host metabolites in influencing intestinal and systemic inflammation. Across a range of organs we investigated the molecular and cellular effects of intestinal inflammation, identifying discriminative molecules for inflammation and molecules common to both intestinal and systemic sites.

## Materials and methods

### Animals and ethical approval

Male C57BL/6 mice (7-8 weeks old) were purchased from Envigo and housed in sterile cages. Mice received a standard laboratory chow diet. Approval for these procedures was given prior to their initiation by an internal University of Glasgow ethics committee and all procedures were carried out in accordance with the relevant guidelines and regulations as outlined by the U.K. Home Office (PPL P64BCA712).

### Induction of colitis and evaluation of colitis clinical scores

Dextran sodium sulphate (DSS) with a molecular weight of 40 kDa was purchased from Alfa Aesar by Thermo Fisher Scientific. Following a 7-day adaptation period, mice were randomly assigned to groups (5 mice per group), consisting of a normal drinking water control or drinking water supplemented with 3% weight over volume DSS for 7 days. Mice were observed daily for clinical symptoms of colitis and body weight was monitored. Immediately before and at the end of the experiment, fresh faeces were sampled from each mouse for gut microbiota compositional analysis.

### Tissue preparation

After euthanasia the entire colon was removed and length was recorded without stretching. Colon and ileum contents were gently removed and the colon cut longitudinally along the mesenteric line and “swissed rolled” before embedding into hydropropyl-methylcellulose/ polyvinylpyrrolidone hydrogel. Tissue was then snap frozen in a slurry of dry ice and isopropanol before rinsing in a slurry of dry ice and isopentane for 30 seconds. Frozen blocks were left on dry ice to allow alcohol evaporation before storing at -80°C. Tissue types including liver, kidney, spleen, and lung were also similarly removed, frozen and stored. Ten micrometre (μM) thick sections were cut from frozen tissue blocks using a cryostat microtome (Thermo Scientific). Consecutive cutting of sections was repeated until enough sections were obtained and these were thaw mounted onto SuperFrost non-conductive microscope slides for DESI-orbitrap-MSI. All slides were dried, vacuum packed and stored at −80°C following sectioning.

### Desorption electrospray ionization mass spectrometry imaging (DESI-MSI)

Vacuum sealed slides were brought to room temperature (RT) before opening and all DESI-MSI experiments were set up as follows. DESI-MSI was performed on a Thermo Scientific Q-Exactive mass spectrometer equipped with an automated Prosolia 2D DESI source. A home-built DESI sprayer assembly was used with the spray tip positioned at 1.5 mm above the sample surface and at an angle of 75°. The distance between the sprayer to mass spectrometer inlet was 7 mm with a collection angle of 10°. The spray solvent was methanol/water (95:5 v/v), delivered at 1.5 μL/ min using a Dionex Ultimate 3000 pump (Thermo Scientific) at a spray voltage of 4.5 kV. Nitrogen was used as the nebulisation gas at a pressure of 7 bars. The Q-Exactive mass spectrometer was operated in positive ion and negative ion mode for all analysis using an S-Lens setting of 75 V. To acquire full mass spectra in the mass range of *m/z* 100–1000 a mass resolution of 70000, AGC target of 5000000 and injection time of 150 ms was used. The .raw files were converted into mzML files using ProteoWizard msConvert and compiled into. imzML format using imzML converter version 1. Data sets were then converted into .slx files and analysed using SCiLS Lab MVS (version 2020b Premium 3D). All DESI data analysis is performed on root mean square (RMS) normalised data. Relative operating characteristic (ROC) curve analysis was performed to visualize the discrimination capability of an *m/z* value between two conditions. Area under the curve (AUC) values were generated by ROC analysis and cut off ranges were set at AUC>0.75 and AUC<0.25 to produce a list of *m/z* that might discriminate between conditions.

### Multivariate and univariate analysis

Following ROC analysis, unsupervised and supervised multiclass classification and correlation were performed to visualise data and highlight metabolites of interest. This multivariate analysis (MVA) was performed using SIMCA 17 software package (Sartorius Stedim Biotech). Unsupervised principal component analysis (PCA) was applied to obtain on overview of data and detect any potential outliers. Subsequently, supervised partial least squares-discriminate analysis (PLS-DA) was used to identify specific changes amongst the groups. In PLS-DA, the *Y* variable was assigned to a defined class and corresponded to *X* variable. The model was generated after the data was autofitted according to R², Q², and classification performance. The quality of the model was validated using two parameters: R2Ycum (goodness of fit) and Q2cum (goodness of prediction). A threshold of >0.5 is widely accepted as good model classification to assure reliable predictive capabilities. Group separation is presented as score plots. Variable importance in projections (VIP) was used as a readout from PLS-DA that reflects the metabolites contribution to the model. A VIP score >1 indicates a higher percent variation in the model when a metabolite is included, whereas VIP<0.5 indicates that a metabolite plays a less important role. Following multivariate analysis, univariate analysis was performed in Graphpad prism (8.4.3). The measurement data were expressed as mean relative abundance ± SD. One-way analysis of variance (ANOVA) was used for comparing more than two groups and nonparametric t-test with 5% false discovery rate (FDR) correction was used for comparing two groups. Statistical significance was established as *p*<0.05.

### Histological staining

After DESI-MSI, tissue sections were fixed by submerging in 4% paraformaldehyde (PFA) for 10 minutes. Sections were then stained with Mayer’s haematoxylin for 1 minute, before rinsing with tap water and submerging in acid alcohol. Tissue was then stained with eosin for 20 seconds, followed by rinsing with tap water and washed 3 times with absolute ethanol.

Tissue sections were then submerged in xylene for 1 minute and cover slips applied using dibutylphthalate polystyrene xylene (DPX) mountant. Haematoxylin and eosin-stained tissue was imaged with an Aperio CS2 digital pathology scanner (Aperio Tech) at 40x and observed in ImageScope software (Aperio Tech).

### Identification of metabolites

Metabolites that were significantly altered were putatively identified using the Human Metabolome Database (HMDB) based on the mass accuracy provided by DESI-MSI analysis. Searches were focused to mass (M) +/- most common adducts, negative mode included M-Hydrogen (H) and M+Chlorine (Cl), positive mode included M+H, M+Potassium (K) and M+Sodium (Na). To further identify metabolites, DESI-Tandem MS (MSMS) was performed on tissue sections using Thermo Scientific Q-Exactive Orbitrap mass spectrometer. The data was acquired in either positive or negative mode with a spray voltage of 4.5kV, 70,000 resolution, maximum injection time of 1000 ms, and S-lens settings of 75 V. Precursor ion mass was set with a mass tolerance of 0.5 *m/z* and the mass range acquired was optimised for each analyte. Higher energy collision dissociation (HCD) with normalised collision energy (NCE) was used for fragmentation of ions during MSMS measurements ranging from 10 to 50 NCE. The identities of the ions were established based on the product ion spectra after comparing to spectra from previously published data using online platforms, HMDB and mzCloud (https://www.mzcloud.org/). Spectra of standards were also acquired using DESI-MSMS and was compared to spectra from tissue, before using *in vitro* experimentation.

### Metabolite enrichment analysis

Significantly changed metabolites are shown as clustered heatmaps generated by the online MetaboAnalyst 5.0 platform (https://www.metaboanalyst.ca/). All available metabolite identities were used in MetaboAnalyst metabolite set enrichment analysis (MSEA). This tool detects major pathways that are associated with the metabolites present in the study. The applied library was the KEGG human metabolic pathways, comprising of 84 metabolite sets. The enrichment ratio was defined as the ratio of observed hits (detected metabolite) per pathway to the count expected by chance (18). One-tailed *p* values were provided after adjusting for multiple testing, enrichment was considered significant when *p*<0.05.

### Imaging mass cytometry

Tissue sections were fixed using 4% paraformaldehyde for 10 minutes at room temperature (RT). This was followed by tissue permeabilization with 1 x casein solution containing 0.1% Triton X-100 for 5 minutes at RT. Tissue was then incubated with blocking buffer (1x casein solution) for 30 minutes at RT inside a humidified chamber. An antibody cocktail was prepared with the appropriate dilution of antibodies (Supplementary Table 1). Tissues were fully covered with antibody solution and incubated overnight in the humidifier chamber at 4°C. DNA Ir-Intercalator (Fluidigm) was diluted 1:400 using phosphate buffered saline (PBS) and pipetted onto tissue, followed by a 30-minute incubation at RT. Tissue was washed in PBS for 5 minutes and repeated three times after each step of the protocol. After the final PBS wash, tissue was rinsed for 30 seconds in deionised water and left to dry at room temperature. Slides were then stored at RT before imaging. Tissue was imaged using the Hyperion Imaging System (Fluidigm), rasterizing at 200 hertz (Hz) with an ablation energy of 5 dB. Laser was frequently tuned between imaging runs to ensure tissue was fully ablated without leaving scratches on the glass. Once acquisition was complete images were exported from MCDviewer as a 32-bit TIFF an imported into HALO image analysis platform (Indica Laboratories) for analysis. Using random forest machine learning tissue classifier module, colon and ileum were segmented into muscularis and mucosa, whilst liver and lung were analysed as whole tissue and vessels based on morphology and blood vessel cell marker, CD31. The Hiplex module was firstly used to segment the DNA intercalator from each cell with a proxy for the cytoplasm being 1 μM radius from the nucleus. Thresholds were then set to define positive cell staining for each marker. Phenotypes for cells of interest were defined using markers cited in current literature (Supplementary Table 2). Percentage positive cells and percentage positive phenotype cells were the defined within tissue regions. Statistical analysis was performed in Graphpad prism, comparisons made for the DSS-treated mice used T-tests to compare treatment groups to control (naïve). Differences between DSS-treated and control samples were considered significant when *p*<0.05.

### Microbiome sequencing and bioinformatic analysis

Fresh faecal samples were collected from the colon of individual mice after culling and immediately stored on dry ice in sterile microfuge tubes before storing at -80°C. DNA was extracted from samples using QIAamp PowerFecal Pro DNA Kit following manufacturer’s protocol (Qiagen). Where possible, stool was collected at the beginning of the experiment and prepared as described to determine microbiome changes over time. The quantity of extracted DNA was measured using Qubit (Thermo Fisher Scientific) before storing at -20°C. 16S rDNA Amplicon sequencing was carried out at Novogene. PCR amplification of the bacterial 16S rRNA genes V3-V4 region was performed using a forward and reverse primer that were connected to barcodes for multiplexed sequencing. PCR products with the proper size were selected by 2% agarose gel electrophoresis. Equal amounts of PCR products were pooled, end-repaired, A-tailed, and further ligated with Illumina adapters. Libraries were then sequenced on a paired-end Illumina platform to generate 250 base pair (bp) paired-end raw reads. Paired-end reads were assigned to samples based on their unique barcodes and truncated by cutting of the barcode and primer sequences. Paired-end reads were merged using FLASH (V1.2.7)(19). Splicing sequences called raw tags were quality filtered under specific conditions according to Qiime (V1.7.0)(20). Tags were compared with reference database (SILVA) using UCHIME algorithm to detect chimera sequences which were subsequently removed. Novogene performed analysis and plotted results using packages in R software (Version 2.15.3).

### Cell-based assays for metabolite characterization

Human intestinal epithelial HCT-8 cells were purchased from the American Type Culture Collection (ATCC) and cultured in Roswell Park Memorial Institute 1640 (RPMI) media supplemented with 10% horse serum (HS), 1% penicillin/streptomycin (P/S), 1% L-glutamine and 5 mM sodium pyruvate. Cells were incubated at 37°C, 5% CO_2_ and passaged every 2-3 days up to a maximum of 30 times. HS was reduced to 3% during experimentation. HCT-8 cells were seeded at 2 x 10⁵ in 24-well plates and treated with the appropriate metabolites (cis-4,7,10,13,16,19-docosahexaenoic acid, creatine, 1-methylnicotinamide – all Merck). All metabolites used in this study were prepared by dissolving in water, filter sterilisation and dilution to the appropriate concentration in RPMI media. After treatment, cells were incubated at 37°C, 5% CO_2_ for 72 hours. Supernatants and lysates were collected and stored followed by the quantification of lactate dehydrogenase (LDH) release, caspase-3/7 activity and cytokine secretion. LDH levels were quantified in cell supernatants using an LDH-Cytotoxicity Assay Kit following manufacturers’ protocol (Abcam, United Kingdom). To quantify caspase-3/7 activity, cells were lysed using 0.1% TritonX-100 and stored at -20°C. Caspase-3/7 activity was quantified using the Apo-ONE® Homogeneous Caspase-3/7 Assay (Promega) according to the manufacturer’s protocol. Assays were read on a FluoStar Optima plate reader (BMG Biotech). The protein concentration of cell lysates was also determined using a bicinchoninic acid (BCA) protein assay kit (Thermo Fisher Scientific) and data presented depicts caspase-3/7 activity as relative fluorescence unit (RFU) per gram of protein. ANOVA was performed for each molecule versus the control condition.

The secretion of TNF-α, IL-8, IL-6 and IL-15 was quantified in cell supernatants using a sandwich ELISA Max Deluxe Set Human (Biolegend), according to the manufacturer’s protocols. The secretion of IFN-γ was also quantified in cell supernatants using a sandwich mouse IFN-γ Duo Set ELISA (DY485-05, R&D systems). Standards and samples were diluted appropriately to obtain results within the stated detection levels. Absorbance was read at 450 nanometres (nm) and 570 nm using FLUOstar Optima plate reader and results are shown as picogram (pg) per ml. The protein concentration of the cell lysate was also determined using a BCA assay (Thermo Fisher Scientific). For each assay ANOVA was performed for each molecule versus the control condition.

### Splenic analysis of immune cell activation and function

Spleens were collected from C5BL/6J mice and kept in in Hanks’ Balanced Salt Solution (HBSS) on ice until further processing. Spleens were mechanically dissociated, homogenised, and passed through a 70 μM cell strainer before undergoing 2 washes in MB (PBS with 0.5% foetal calf serum (FCS) and 2 mM ethylenediaminetetraacetic acid (EDTA)). Red blood cells (RBC) were lysed with 1-2 ml of RBC lysis buffer for 3-5 minutes (Thermo Fisher Scientific). Following another washing step cells were counted and cultured in 96 well round-bottom plates at a final concentration of 2.5×10⁶/ml in complete RPMI (RPMI with 10% FCS, 1% L-glutamine and 1% P/S). Cultured cell suspensions were stimulated with either recombinant mouse IL-12 and IL-18 to activate innate and adaptive immune effector cells (both at 5 ng/ml), IL-4 to activate humoral and adaptive immune cells (10 ng/ml) or appropriately diluted metabolites. Cells were incubated overnight at 37°C, 5% CO_2_. In the final 4 hours of culture brefeldin A and monensis (eBioscience) were added to block secretion of intracellular cytokines for detection by cytometry.

Cell culture plates were centrifuged at 1400 rpm, 4°C for 7 minutes and supernatants were stored at -20°C until further use. Live/dead stain (eBioscience Fixable Viability Dye eF506) was diluted 1:1000 in clean PBS and incubated with cells for 20 minutes at 4°C. Cells were vortexed to resuspend pellet before adding 150 μL MB to each well. Isotypes were generated by taking equal volumes of each sample from 1 condition and pooling together in spare wells.

Cells were stained with antibodies to CD69 (clone H1.2F3, Biolegend), CD19 (clone 6D5, Biolegend), NK-1.1 (clone PK136, Biolegend), granzyme B (clone GB11, Biolegend), IFN-γ (clone XMG1.2, Biolegend), CD3 (clone 17A2, Biolegend), CD40 (clone 5C3, Biolegend), MHCII (clone M5/114.15.2, Biolegend) and CD45 (clone HI30, Biolegend). Surface stain was performed for 45-60 minutes at 4°C. In the final 5 minutes of staining, intracellular fixation buffer was added and washed with MB. For intracellular cytokine staining, cells were first fixed with 1x Fix/Perm buffer (eBioscience Foxp3 kit) for 1 hour on ice. Intracellular staining was performed overnight at 4°C. Following washing the cell preparations were filtered on 70 μM nylon mesh and samples were analysed on flow cytometers (BD Celesta). Data was analysed using FlowJo software (Tristar).

## Results

### Spatial metabolomic mapping of the murine ileum and colon during DSS-induced intestinal colitis

Sequencing of faecal samples identified changes in the relative abundance of bacterial phyla and genera between day 0 and day 7 of the experiment when mice were treated with 3% DSS (Fig. S1). Firmicutes and Bacteroidota remained the most abundant phyla in the 3% DSS treated group, however microbial diversity was reduced with an increase in the abundance of Proteobacteria. Genera including *Clostridium, Escherichia, Lactobacillus, Parabacteroides, Bacteroides* and *Sutterella* were increased in DSS treated mice, with *Ruminococcus, Prevotella, Lachnospiraceae* and *Faecalibacterium* reduced.

MSI was next employed to elucidate molecular changes across a range of intestinal and systemic sites during DSS colitis. Mass spectra were acquired (*m/z* 50-1000 Da) for each tissue type to identify metabolites and lipids and ROC analysis was performed to identify metabolites which could discriminate between control and DSS-treated mice. Unsupervised and supervised multivariate analysis was performed on these metabolites see which contributed most to separation of the treatment groups and univariate analysis was performed to further validate the metabolite differences between the groups. Metabolites with a VIP score >1 and/or a *p* value of <0.05 are reported here.

Our analysis identified 88 metabolites significantly changed by DSS treatment (Fig. S2 and Supplementary Tables 3 and 4). Of these 49 could be putatively identified using the Human Metabolome Database (HMDB) (Fig. 1A). In the colon, (Fig. S3 and Supplementary Tables 5 and 6) univariate analysis indicated that 3 metabolites were significantly decreased, and 27 metabolites were significantly increased in the 3% DSS treated mice compared to the control. Twenty of these metabolites could be assigned a putative identity (Fig. 1B). A very large number of metabolites were putatively identified but, given their number, not all could have their identities confirmed by MSMS. However, MSMS was used to confirm the molecular identities of a number of metabolites by matching their spectra to those available online, these included *m/z* 132.07-creatine, *m/z* 162.11-carnitine, *m/z* 259.24-arachidonic acid, *m/z* 327.23-docosahexaenoic acid (DHA), *m/z* 317.21-leukotriene A4 and *m/z* 351.21, lipoxin A4 and gamma-linolenic acid (GLA).

**Figure 1A.**
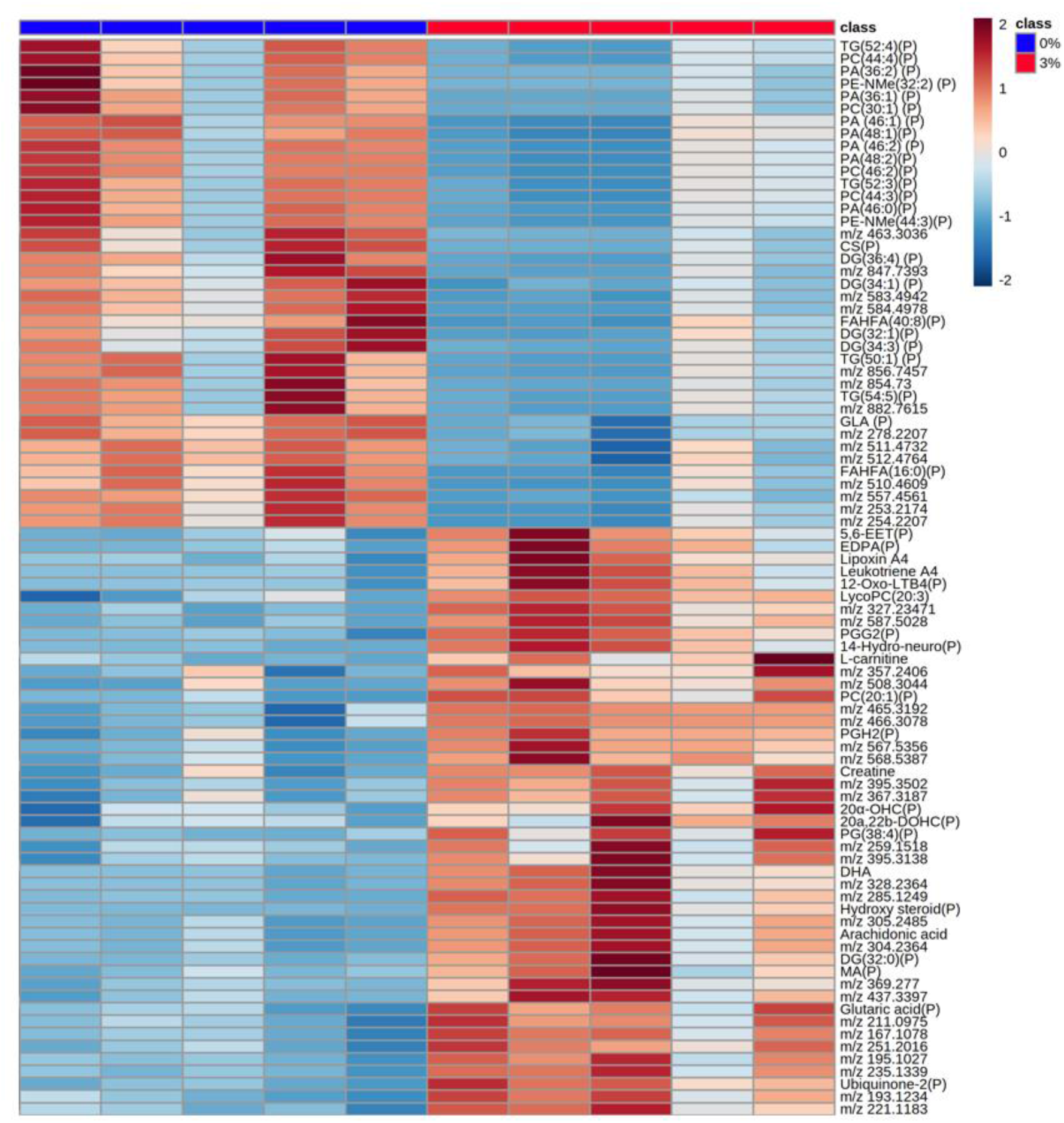
Heatmap of metabolites increased and decreased in abundance in the ileum of 3% DSS treated mice compared to controls. Heatmap shows *m/z* of molecules unable to be identified, putatively identified molecules (marked with a P) and molecules with an MSMS confirmed identification. Heatmap was created using Metaboanalyst 5.0. The colour of each sample is proportional to the significance of change in metabolite abundance indicated by the colour bar (red, up-regulated in DSS colitis group; blue, down-regulated in colitis group). Rows correspond to metabolites and columns correspond to samples.

**Figure 1B.**
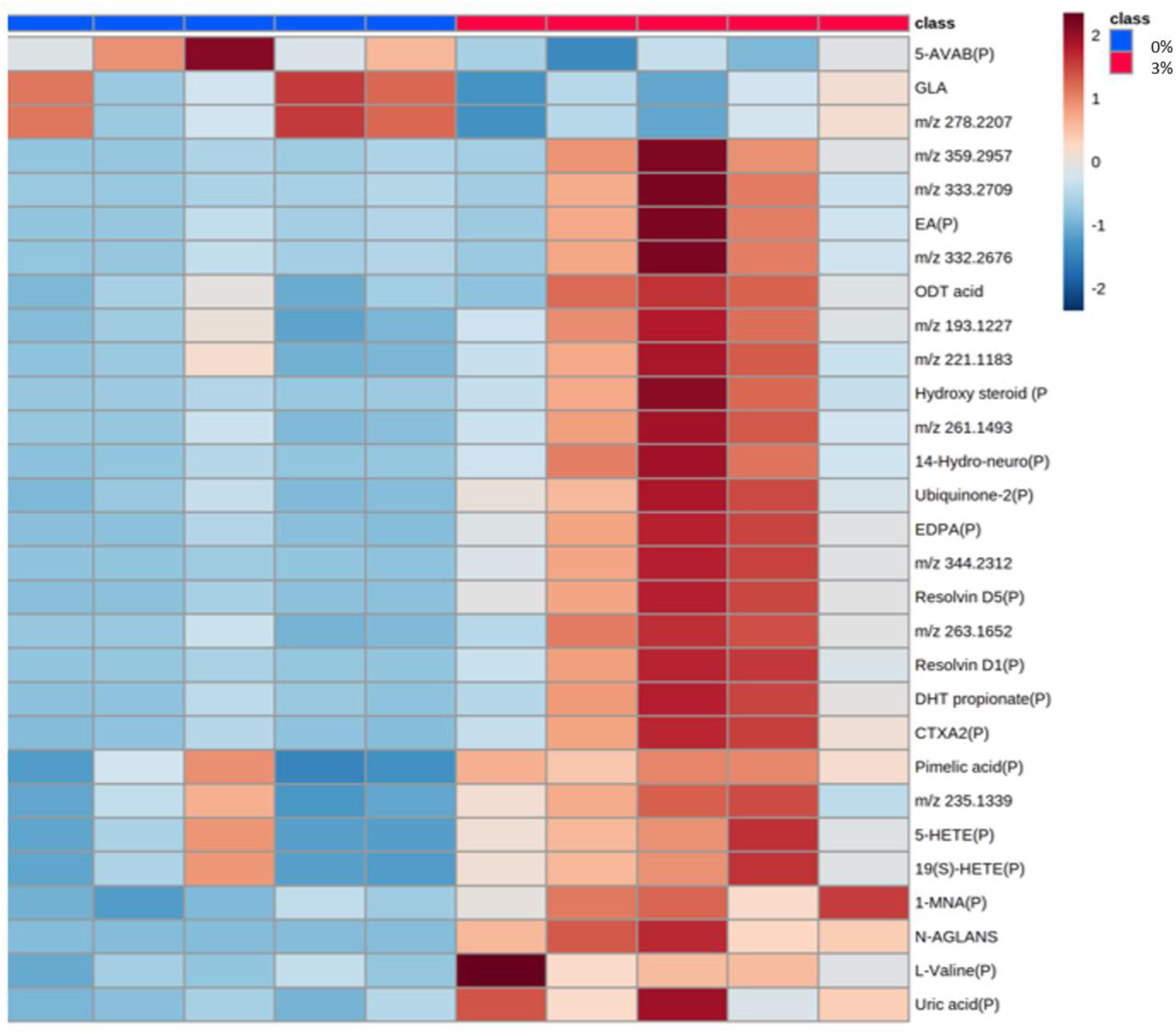
Heatmap of metabolites increased and decreased in abundance in the colon of 3% DSS treated mice compared to controls. Heatmap shows *m/z* of molecules unable to be identified, putatively identified molecules (marked with a P) and molecules with an MSMS confirmed identification. Heatmap was created using Metaboanalyst 5.0. The colour of each sample is proportional to the significance of change in metabolite abundance indicated by the colour bar (red, up-regulated in DSS colitis group; blue, down-regulated in colitis group). Rows correspond to metabolites and columns correspond to samples.

### KEGG Pathway enrichment analysis of significantly changed metabolites

Given the number of metabolites changed across the intestine, KEGG pathway enrichment analysis was used to identify pathways significantly altered by treatment. Twelve pathways were enriched in the ileum of DSS-treated mice, but only the arachidonic acid (AA), steroid hormone biosynthesis and primary bile acid biosynthesis pathways were significantly enriched (Fig. S4). Fourteen KEGG pathways were identified through enrichment analysis in the colon, but in the colon only the AA metabolism pathway was significantly enriched (Fig. S5).

### Mapping the effects of identified metabolites on the intestinal immune response

To better understand cellular and functional changes in the intestinal immune response post-DSS treatment, imaging mass cytometry (IMC) was employed. To guide this analysis the locations of specific metabolites, identified as significantly changed in response to treatment, were used as a guide. As 128 metabolites were changed in the ileum and colon during 3% DSS treatment, metabolites were selected that were significantly changed by DSS treatment and which had 3 characteristics: an interesting spatial distribution, a VIP>1, and an identity confirmed by MSMS. Metabolites fitting these criteria were docosahexaenoic acid (DHA) (*m/z* 327.233, VIP=1.057), which was increased 2.66-fold (*p*=0.00197), creatine (*m/z* 132.0771, VIP=1.050) which was increased 2.29-fold (*p*=0.0054) in the ileum, and lastly 1-methylnicotinamide (1-MNA) (*m/z* 137.071, VIP=1.605) which increased 2.59-fold (*p*=0.001162) in the colon (Fig. 2). IMC was performed at a 1 μM spatial resolution (on tissue regions indicated by blue rectangular boxes in Fig. 2), revealing changes in structural and immune protein markers. Individual cells were segmented and a random forest machine learning tissue classifier segmented the colon and ileum into mucosa and muscularis regions. In the ileal mucosa, the percentage of cells positive for a number of markers were decreased in the DSS-treated mice compared to controls, each approximately 2-fold, including F480, CD69, Ki67, CD4 and MHCII (Fig. S6). The markers identified via IMC were grouped to define single-cell phenotype and function such as proliferation, apoptosis, and activation state. In the ileal mucosa post-DSS treatment, there was a decrease in helper T cells (CD3^+^, CD4^+^), dendritic cells (CD11c^+^, MHCII^+^), M2b macrophages and M1 macrophages (F480^+^), and proliferative cells (Ki67^+^)(Fig. 3). While no individual markers were significantly changed in the ileal muscularis of 3% DSS treated mice compared to control mice, the M1 macrophage population decreased 2.81-fold (*p*=0.0319) post-treatment. In the colonic mucosa, 20 markers of cellular function, proliferation and structure were altered between the 3% DSS treated group and the control (Fig. S7). Those markers that decreased included ATPase, Pan-CK, CD103, CD11c, E Cadherin, Glut1, Ki67 and EpCAM, with each decreasing approximately 5-fold or less. Significant, and larger, fold increases were seen in number of markers post-DSS treatment including vimentin, CD68, Ly6G, CD11b, F480, CD163, collagen, Lyve1, FoxP3, NKp46, CD4, and granzyme B. However, in the case of Ly6G and CD68, these increases were approximately 20-fold and 40-fold, respectively, far larger than those of any other markers. The abundance of all these markers was then used to predict the dysregulation of 14 cellular phenotypes and functions in DSS-treated mice. The percentage of positive cells that were proliferative (Ki67^+^), hypoxic (Glut1^+^), or antigen-presenting dendritic cells (CD11c^+^, MHCII^+^, CD103^+^) was significantly reduced in the colonic mucosa of the DSS-treated mice (Fig. 4). In contrast the percentage of a range of cellular phenotypes including helper T cells (CD3^+^, CD4^+^), regulatory T cells (CD3^+^, CD4^+^, FOXP3^+^), natural killer (NK) cells (NKp46^+^), neutrophils (Ly6G^+^, CD11b^+^), blood vessels (CD31^+^) and mesenchymal cells (vimentin^+^) was significantly increased after DSS treatment, with neutrophil levels increasing greater than 300-fold. The percentage of macrophages (F480^+^, CD68^+^, CD11b^+^) was increased greater than 50-fold with subsets including M2c (F480^+^, CD163^+^), M2a (F480^+^, CD206^+^), M2b (F480^+^) and M1 (F480^+^, MHCII^+^) all contributing through their own significant fold-change increase in the colonic mucosa post-DSS treatment. In the colonic muscularis of the 3% DSS-treated group, the percentage of CD11b^+^ (8.24-fold, *p<*0.0001), F480^+^ (7.20-fold, *p*=0.0008) and granzyme B^+^ (7.00-fold, *p*=0.0234) cells increased compared to the control, while the percentage of M2c (6.15-fold, *p*=0.0009), M1 and M2 macrophages (7.20-fold, *p*=0.0008), and M1 macrophage positive cells (3.85-fold, *p*=0.0419) again all significantly increased in a manner similar to that observed in the mucosa of the 3% DSS treated group.

**Figure 2.**
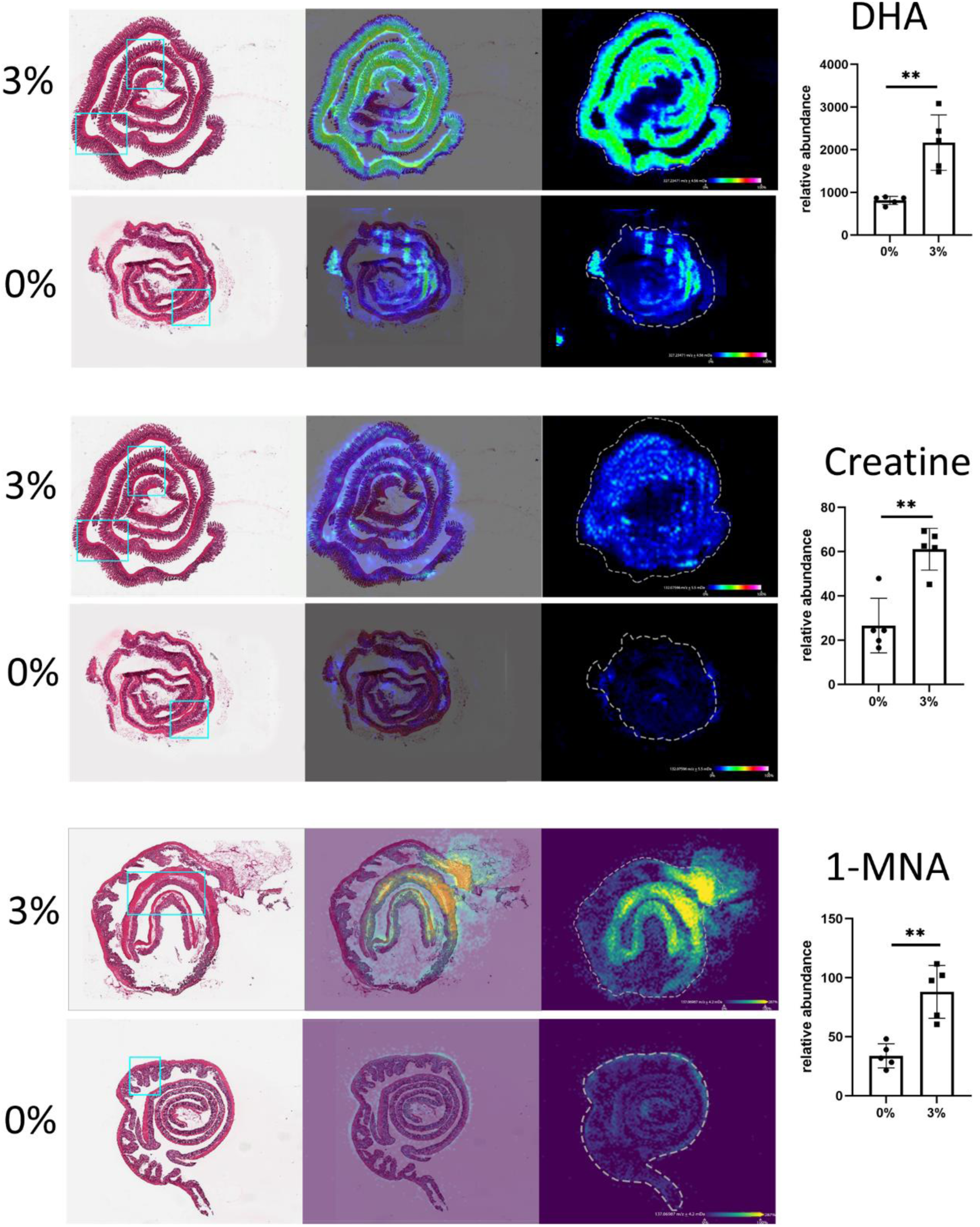
MSI of DHA, creatine and 1-MNA abundance in the intestine. From left to right; haematoxylin and eosin (H&E)-stained sections, overlayed image of H&E image and MSI of specific molecule, MSI heatmap of molecules (colour bar shows 0% to 100% relative abundance). Bar plot shows the relative abundance of DHA and creatine in the ileum and 1-MNA in the colon. Data are shown as the mean of five biological replicates ± standard deviation (SD) (error bars). A multiple T-test was performed with FDR 5% adjustment to compare metabolite abundance between the groups (0% DSS and 3% DSS) \**p*<0.05, \*\**p*<0.01, \*\*\**p*<0.001, \*\*\*\**p*<0.0001 was considered statistically significant. Blue rectangular box shows the tissue region selected for IMC.

**Figure 3.**
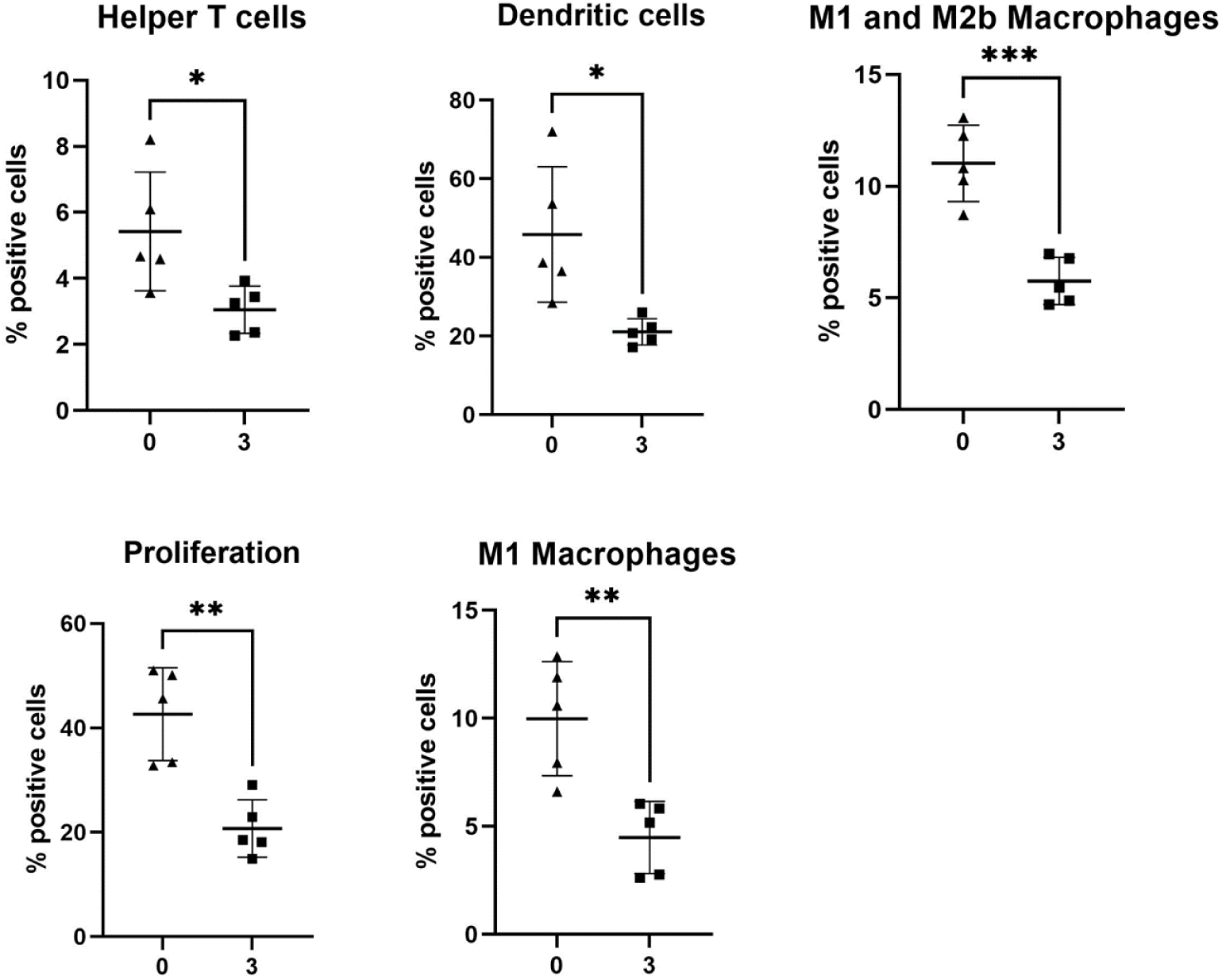
Cell phenotype and function are significantly changed in the ileal mucosa of DSS-treated mice. Percentage positive cells represented as bar graph showing five biological replicates. Helper T cells (CD3^+^, CD4^+^)(1.78-fold, *p*=0.0252), dendritic cells (CD11c^+^, MHCII^+^)(2.17-fold, *p*=0.0136), M2b (1.91-fold, *p*=0.0004) and M1 macrophages (F480^+^)(2.22-fold, *p*=0.0043), and proliferation positive cells (Ki67^+^)(2.05-fold, *p*=0.0016) were decreased in the 3% DSS treated colitis group compared to the control. A t-test was performed to compare the two groups and \**p*<0.05, \*\**p*<0.01, \*\*\**p*<0.001, \*\*\*\**p*<0.0001 were considered statistically significant.

**Figure 4.**
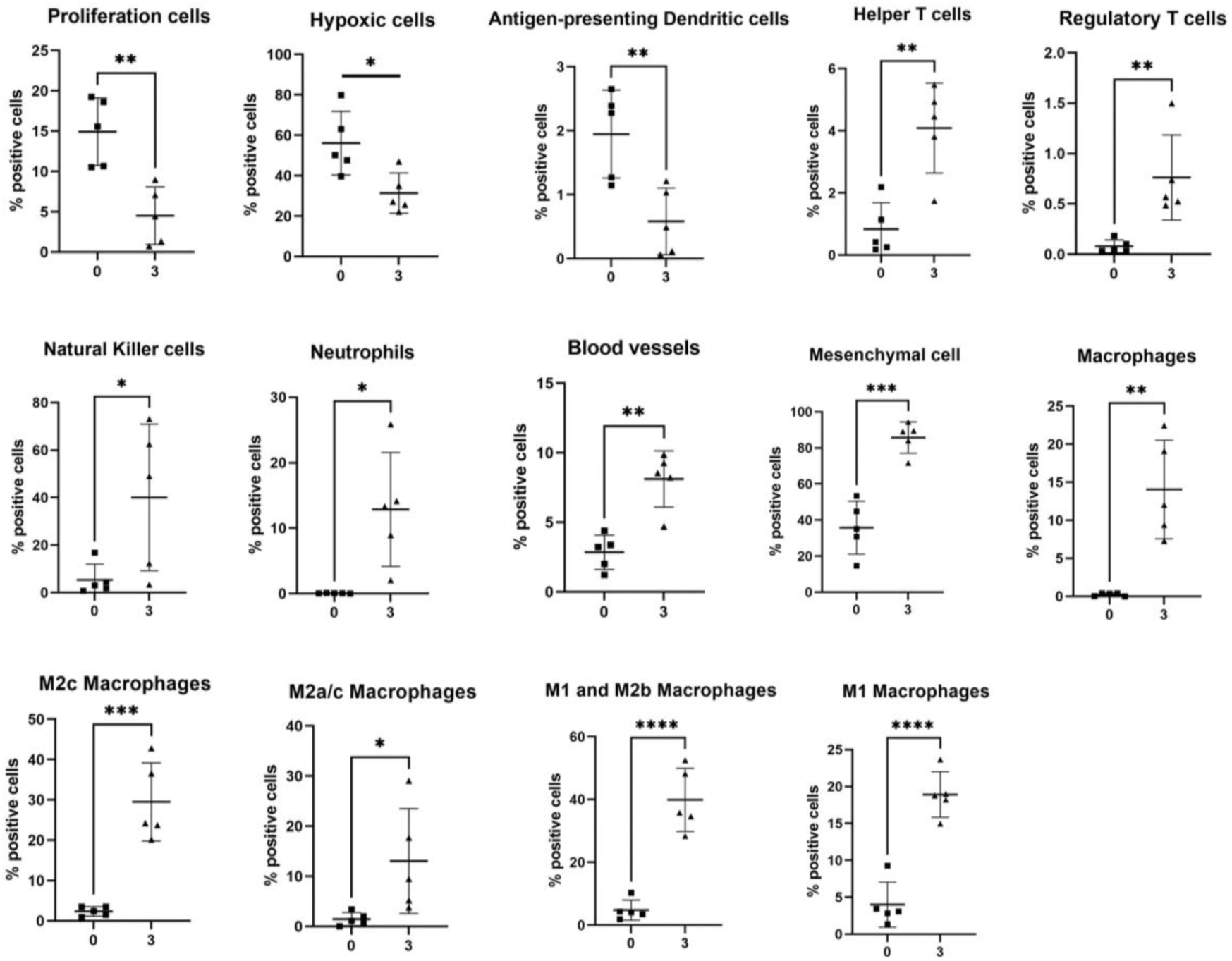
Bar graph of cell phenotype and function significantly changed in the colonic mucosa of DSS-treated mice. Percentage positive cells represented in bar graphs showing five biological replicates. Proliferative (Ki67^+^), hypoxic (Glut1^+^), and antigen-presenting dendritic cells (CD11c^+^, MHCII^+^, CD103^+^) were reduced post-treatment. Helper T cells (CD3^+^, CD4^+^), regulatory T cells (CD3^+^, CD4^+^, FOXP3^+^), natural killer (NK) cells (NKp46^+^), neutrophils (Ly6G^+^, CD11b^+^), blood vessels (CD31^+^), mesenchymal cells (vimentin^+^), macrophages (F480^+^, CD68^+^, CD11b^+^) and macrophage subsets including M2c (F480^+^, CD163^+^), M2a(F480^+^, CD206^+^), M2b (F480^+^) and M1 (F480^+^, MHCII^+^) were increased in the 3% DSS group compared to the control. A t-test performed to compare the two groups and \**p*<0.05, \*\**p*<0.01, \*\*\**p*<0.001, \*\*\*\**p*<0.0001 were considered statistically significant.

### Testing of immunomodulatory effects of specific metabolites *in vitro* and *ex vivo*

To further elucidate how the metabolites utilized to select regions for IMC may be contributing to the altered immune profile these metabolites; creatine, DHA and 1-MNA, were tested *in vitro*. Firstly, their identity was confirmed by MSMS (Fig. S8A-C). Lactate dehydrogenase and caspase-3 activity assays indicated that creatine, DHA and 1-MNA have cytotoxic effects on intestinal epithelial cells but do not induce apoptosis (Fig. S9). Cytokine release in response to the metabolites varied, but none of the metabolites resulted in IL-6 release (Fig. S10). 50 μM creatine or 100 μM DHA significantly reduced IL-8 release while 100 μM DHA was observed to significantly increase the release of TNF-α. Increasing the concentration of creatine, DHA and 1-MNA to 100 μM, from 50 μM, resulted in increased release of IL-15 in response to all the metabolites (Fig. S10).

After determining that creatine and DHA have an immunomodulatory effect on IECs we next established their effect on immune cells. Immune cells were isolated from mouse spleens and stimulated with creatine, DHA, or a cytokine control (IL-4 or IL-12+IL-18) (Fig. S11). While DHA did not significantly change the percentage of any immune cell type, stimulating cells with creatine increased the percentage of NK cells expressing CD69 (2.67-fold, *p*=0.0007)(Fig. 5). A proteome profiler mouse cytokine array indicated that IFN-γ was increased in the cell supernatant following molecular exposure and IFN-γ levels were quantified by ELISA. Creatine treatment did not significantly alter IFN-γ release compared to the control but treating cells with DHA increased the secretion of IFN-γ 13-fold (*p*<0.0001)(Fig. 5).

**Figure 5.**
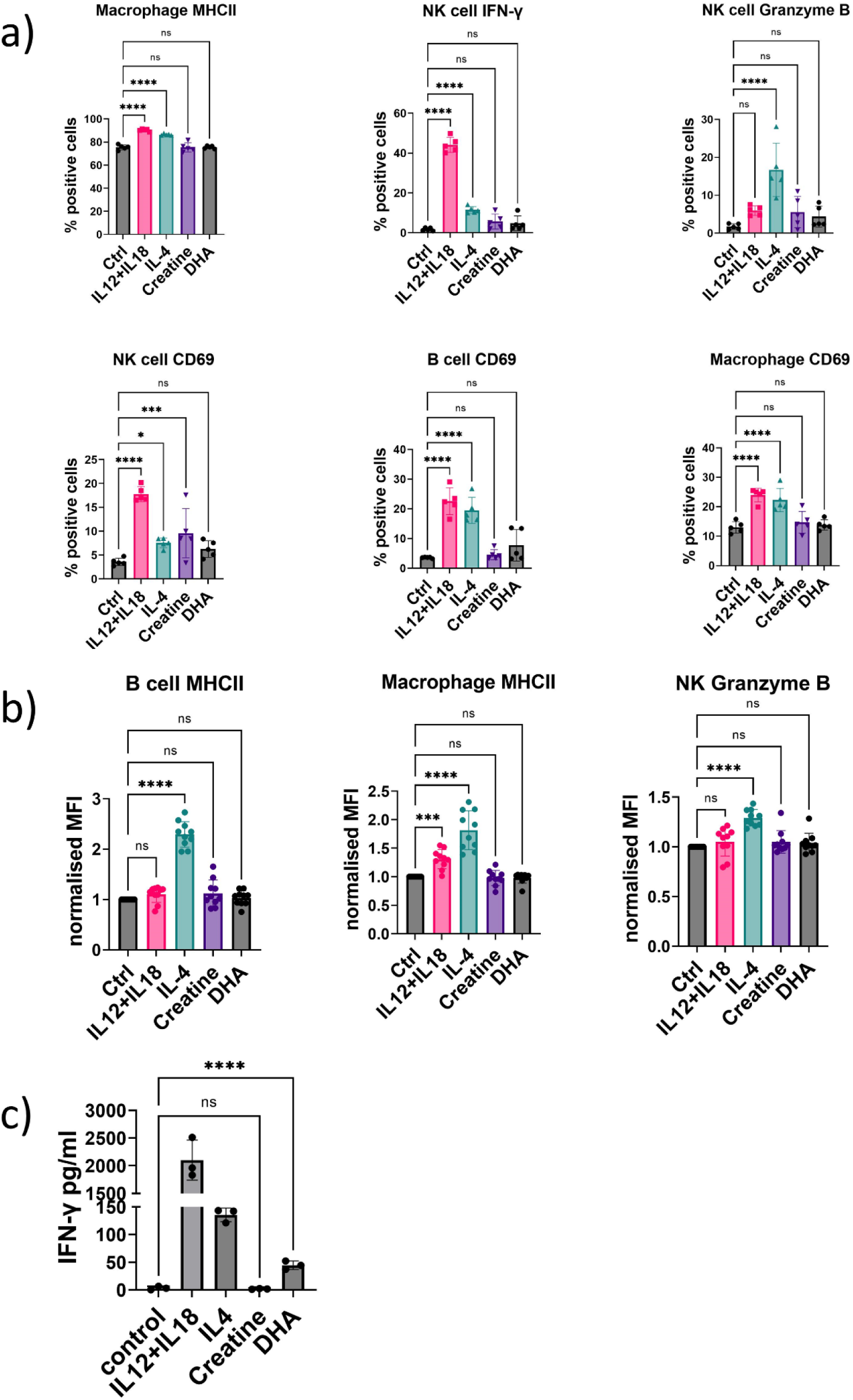
Flow cytometry and cytokine release analysis of splenic immune cells following stimulation with creatine and DHA. Immune cells were isolated from spleen tissue and stimulated with 50 μM creatine or DHA as well as IL12+IL-18 and IL4 as positive immune activating controls. A) Percentage of immune cells (NK cells, B cells and macrophages) positively expressing specific markers (MHCII, IFN-γ, granzyme B and CD69). B) Mean fluorescence intensity (MFI) was normalised to control (no stimulation) and shows the relative amount of expressed marker within a positive population. A-B) One-way ANOVA was performed to compare stimulation versus the control condition (cells without treatment). C) An ELISA was used to quantify the release of IFN-γ into immune cell supernatant and is shown as pg/ml. An unpaired t-test was performed to compared creatine or DHA stimulated groups to control and did not include positive controls (cytokine stimulations). Data is shown as the mean of 3 biological replicates ± standard deviation (SD). \**p*<0.05, \*\**p*<0.01, \*\*\**p*<0.001, \*\*\*\**p*<0.0001 were considered statistically significant.

### Spatial metabolomic mapping of the murine liver, spleen and kidney during DSS-induced intestinal colitis

Given the changes in metabolite abundance seen in the intestine post-treatment of mice with 3% DSS, systemic metabolomic changes were next determined focusing on the liver, spleen, kidney and lung. Univariate analysis post-MSI of the liver indicated 239 metabolites were significantly changed with putative identities assigned to 125 metabolites (Fig. S12 and Supplementary Tables 7 and 8). Given the large number of metabolites identified in the liver these could not be depicted in a heatmap. In the spleen univariate statistical analysis identified 65 peaks that were significantly changed with 34 molecules assigned a putative identity (Fig. S13 and Supplementary Tables 9 and 10). Metabolites significantly changed in the spleen are shown as a heatmap (Fig. 6). In the kidney of DSS-treated mice 16 metabolites were significantly changed between the groups following univariate analysis (Fig. S14 and Supplementary Tables 11 and 12) and are shown in a heatmap (Fig. 7). This study could assign 11 putative identities to the metabolites using the HMDB. MSI was also performed on lung tissue but no significantly changed metabolites were found between control and DSS-treated animals.

**Figure 6.**
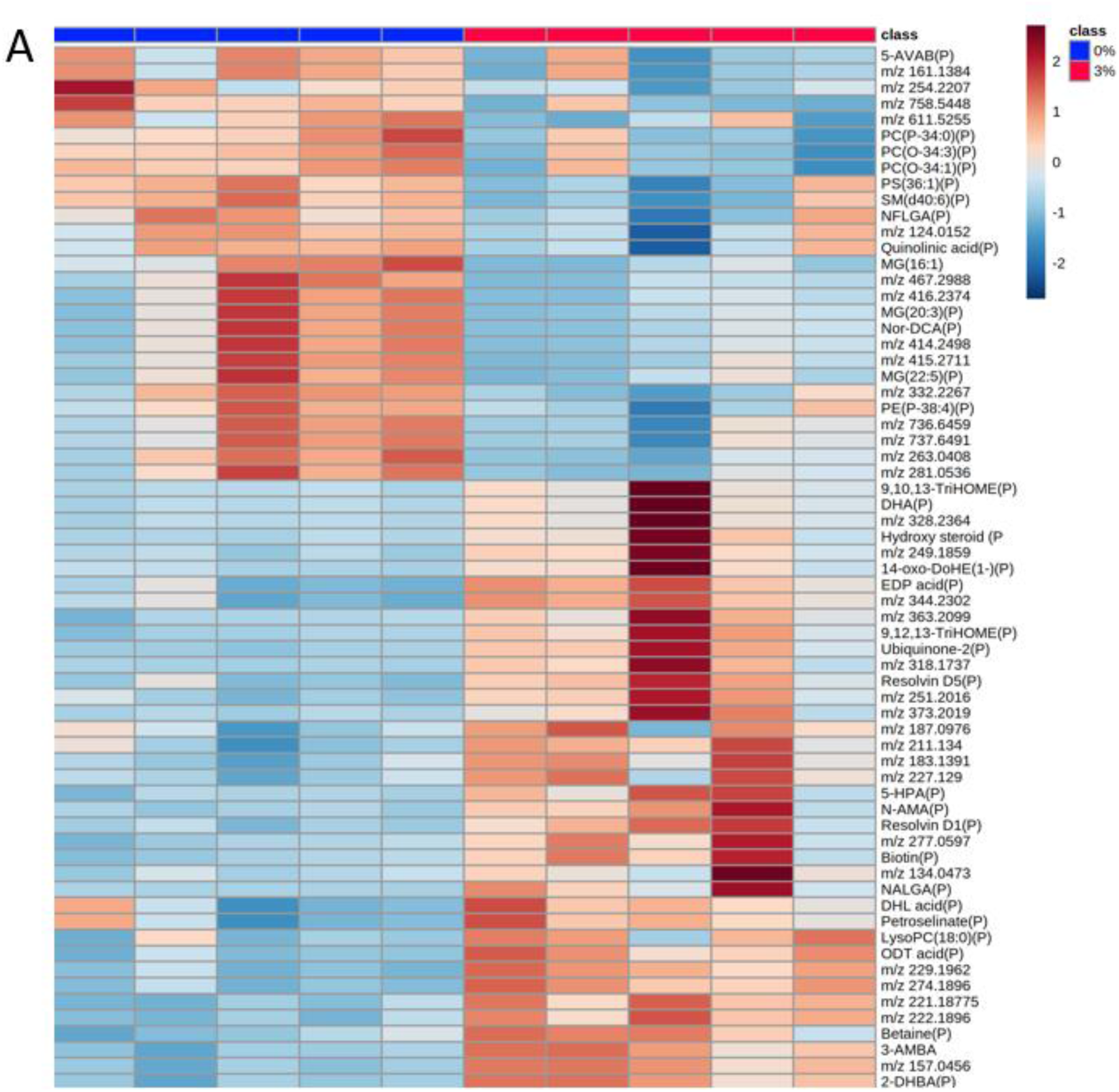
Heatmap of metabolites increased and decreased in abundance in the spleen of 3% DSS treated mice compared to controls. Heatmap shows *m/z* of unidentified molecules and putatively identified molecules (marked with a P). Heatmap was created using Metaboanalyst 5.0. The colour of each sample is proportional to the significance of change in metabolite abundance indicated by the colour bar (red, up-regulated in DSS colitis group; blue, down-regulated in colitis group). Rows correspond to metabolites and columns correspond to samples.

**Figure 7.**
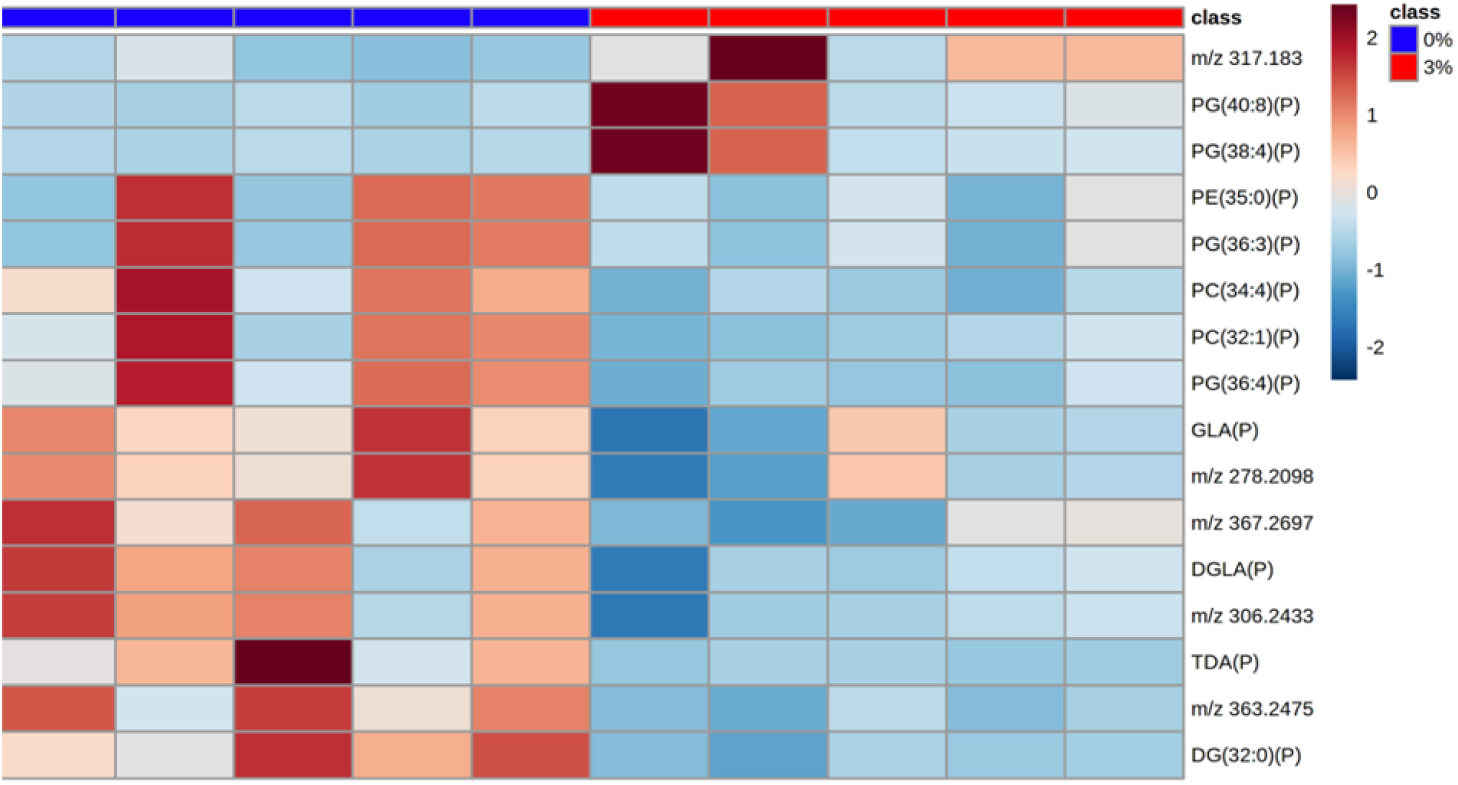
Heatmap of metabolites increased and decreased in abundance in the kidney of 3% DSS treated mice compared to controls. A) Heatmap shows *m/z* of molecules unable to be identified and putatively identified molecules (marked with a P). Heatmap was created using Metaboanalyst 5.0. The colour of each sample is proportional to the significance of change in metabolite abundance indicated by the colour bar (red, up-regulated in DSS colitis group; blue, down-regulated in colitis group). Rows correspond to metabolites and columns correspond to samples.

KEGG enrichment analysis indicated that the metabolites found in the liver dataset are involved in at least 25 pathways (Fig. S15) but only the galactose metabolism and biosynthesis of unsaturated fatty acids pathways were significantly enriched. No significantly enriched pathways were identified in the spleen (Fig. S16), and in the kidney the small number of identified metabolites prevented KEGG pathway analysis being informative.

### Identification of putative biomarkers of colitis at systemic sites

Next, we identified molecular biomarkers of intestinal inflammation that were also altered at systemic sites which may contribute to systemic inflammation. To guide this approach we identified metabolites that were significantly changed in abundance at both intestinal and systemic sites. A number of metabolites were significantly altered both in the ileum and colon in response to DSS. These included metabolites increased in both tissues such as 11b-hydroxyandrost-4-ene-3,17-dione (*m/z* 301.1658; ileum 15.8-fold, *p*=0.03; colon 4.84-fold, *p*=0.001), ubiquinone-2 (*m/z* 317.1759; ileum 3.06-fold, *p*=0.0012; colon 7.9-fold, *p*=0.02), epoxydocosapentaenoic acid (*m/z* 343.228; ileum 3.40-fold, *p*=0.0038; colon 7.1-fold, *p*=0.02) and 14-hydroperoxy-H4-neuroprostane (*m/z* 391.2126; ileum 4.45-fold, *p*=0.0019; colon 12.64-fold, *p*=0.023). In contrast *m/z* 277.2172, putatively identified as gamma-linolenic acid was significantly decreased in both the colon (2.09-fold, *p*=0.045) and ileum (2.11-fold, *p*=0.0012) post-DSS treatment. The metabolites with *m/z* 193.1234 and *m/z* 221.1183 were significantly increased in both the ileum (1.78-fold, *p*=0.0019) and colon (2.32-fold, *p*=0.0239) of DSS-treated mice but no putative identity could be assigned to them.

Metabolites significantly increased across the whole of the intestine were then compared against those from systemic sites to determine whether the same metabolites were significantly increased at all sites. Only two metabolites were altered significantly in each of the ileum, colon and liver with ubiquinone-2 increased in the liver of DSS-treated mice in a manner similar to that seen in the intestine, while gamma-linolenic acid was similarly decreased in the liver (Fig. S17). Ubiquinone-2 was again significantly increased in the spleen, alongside 11b-hydroxyandrost-4-ene-3-17-dione and epoxydocosapentaenoic acid (Fig. S18), while gamma-linoleic acid was the only metabolite significantly altered in the kidney where it was again decreased in a manner similar to the intestine (Fig. S19). So while ubiquinone-2 and gamma-linoleic acid were found at two systemic sites where their abundance was similarly altered when compared to the intestine, no metabolite was found to be altered, in both the intestine and all systemic sites tested, in a similar manner.

### Intestinal colitis results in a dysregulated immune cell profile in the liver

As MSI revealed hundreds of metabolite changes in the liver during colitis, IMC was employed for single-cell phenotyping to identify changes in function. As MSI heatmaps indicated that, unlike in the intestine most metabolites were homogenous throughout liver tissue, the sections were selected for IMC at random. The liver was segmented into regions, tissue and blood vessels based on tissue morphology. In the tissue of the 3% DSS treated group, the percentage of CD68^+^, CD45^+^, F480^+^, CD163^+^, E cadherin^+^, collagen^+^, Glut1^+^ and CD206^+^cells increased (Fig. S20). The markers identified were used to predict changes in specific cell types and the results showed that in the 3% DSS treated group, the percentage of regulatory T cells (CD3^+^, CD4^+^, FOXP3^+^) and activated cytotoxic T cells (CD3^+^, CD8^+^, granzyme B^+^) was significantly reduced 2.44-fold (*p*=0.0493) and 14.42-fold (*p*=0.0244), respectively, compared to controls. However, the percentage of macrophage cells in the subclass M2c (F480^+^, CD163^+^) and M2a (F480^+^, CD206^+^) were significantly increased 2.42-fold (*p*=0.0050) and 2.32-fold (*p*=0.0018), respectively, in the 3% DSS-treated group. Lastly, the percentage of cells that were hypoxia positive (Glut1^+^) increased 2.03-fold (*p*=0.0271) during treatment compared to the control (Fig. 8).

**Figure 8.**
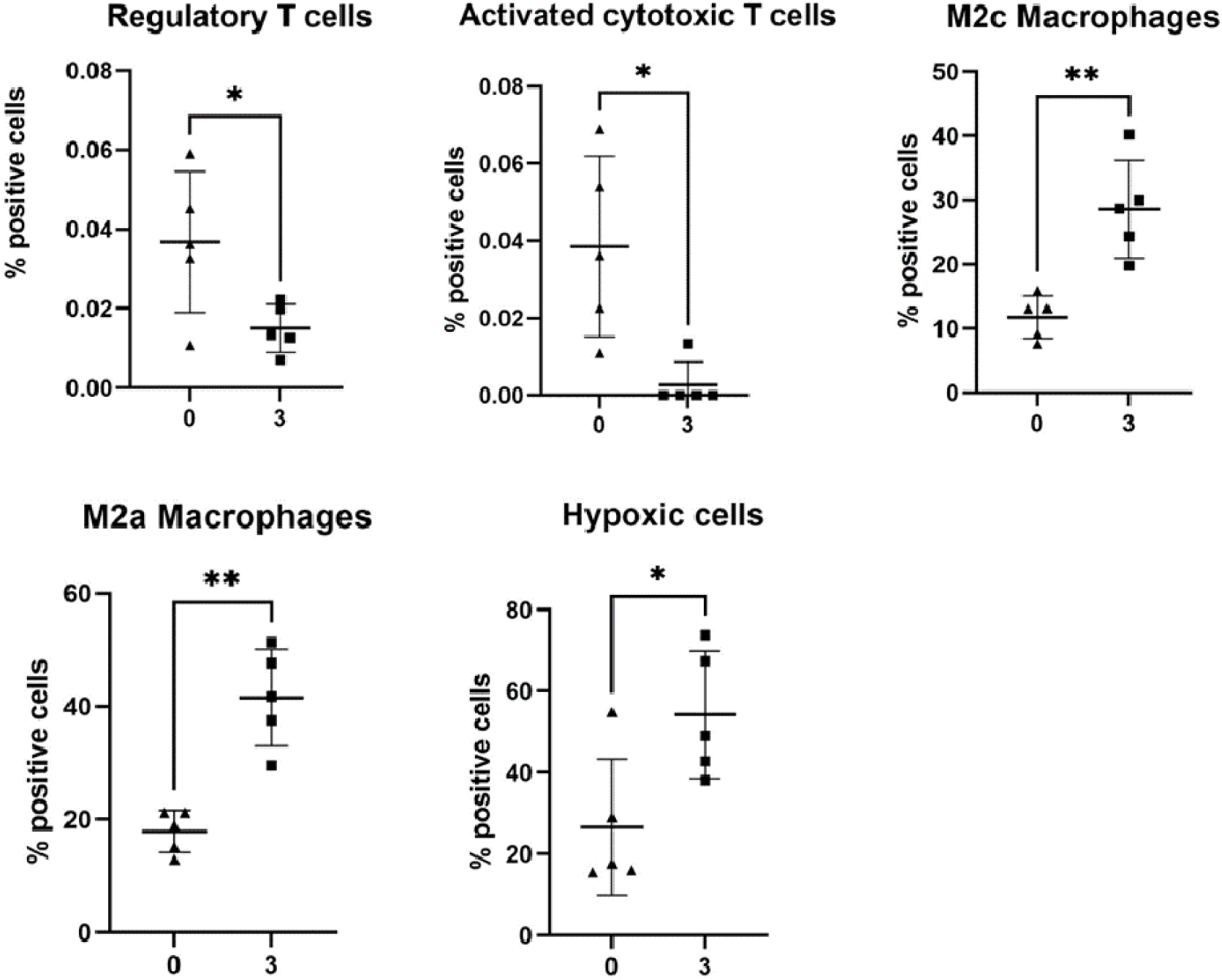
Bar graphs denoting significant changes in cell phenotype and function in the liver of 3% DSS-treated mice. Percentage positive cells represented by bar graph showing five biological replicates. Regulatory T cells (CD3^+^, CD4^+^, FOXP3^+^) and activated cytotoxic T cells (CD3^+^, CD8^+^, granzyme B^+^) were significantly reduced, whereas the percentage of macrophages in the subclass M2c (F480^+^, CD163^+^) and M2a (F480^+^, CD206^+^) and hypoxia (Glut1^+^) were significantly upregulated in 3% DSS treated group compared to controls. A t-test performed to compare the two groups and \**p*<0.05, \*\**p*<0.01, \*\*\**p*<0.001, \*\*\*\**p*<0.0001 were considered statistically significant.

In liver vessels, the percentage of cells expressing the following markers increased in the 3% DSS-treated group compared to controls; CD68 (2.50-fold, *p*=0.0346), F480 (2.25-fold, *p*=0.0005), CD11b (2.25-fold, *p*=0.0396), CD163 (2.98-fold, *p*=0.0169), E cadherin (1.83-fold, *p*=0.0162), Lyve1 (2.29-fold, *p*=0.0179), NKp46 (1.07-fold, *p*=0.0221), CD206 (1.51-fold, *p*=0.0092), Glut1 (1.38-fold, *p*=0.0044) and ATPase (1.04-fold, *p*=0.0006), respectively. Furthermore, the percentage of M2c (3.26-fold, *p*=0.0081) and M2a (2.73-fold, *p*=0.0005) macrophage positive cells significantly increased in the 3% group compared to controls while vessels in DSS-treated mice had a 1.38-fold (*p*=0.0044) increase in the percentage of cells positive for hypoxia. This indicates that during intestinal colitis the liver undergoes injury via hypoxia.

## Discussion

The gut microbiome exerts its effects through direct interactions with the intestinal epithelium and through the release of countless metabolites that exert their effects both locally and systemically. Here, using a spatial biology approach we gained an understanding of the influence of gut microbiota disruption and intestinal inflammation in the DSS mouse model of colitis.

In DSS-treated mice the arachidonic acid (AA) pathway was significantly enriched in both the ileum and colon. AA is hydrolysed by phospholipase A2 (PLA_2_) into a free form before its breakdown to produce bioactive eicosanoids such leukotriene B4 (LTB_4_), a potent chemoattractant which is one of the main mediators of neutrophil recruitment in the intestine (21). While the AA pathway was enriched in the ileum and colon, there were differences in the AA-derived metabolites detected at each site. In the ileum 20(S)-HETE, prostaglandin G2, thromboxane B2 and LTB_4_ were increased, whereas 11,12-EET, 15S-HETE and 5(S)-HETE were increased in the colon. It is likely that this diversity of metabolites influences the distinctive cellular profiles detected at each site by IMC (22). 5(S)-HETE is synthesized by neutrophils in response to infection, eliciting a strong pro-inflammatory response by increasing transepithelial neutrophil chemotaxis and neutrophil degranulation with the consequent release of anti-bacterial products and reactive oxygen species contributing to further dysbiosis, tissue injury and inflammation (23, 24). In contrast in the ileum, helper T cells, dendritic cells, M2b and M1 macrophages and proliferation positive cells in general, were decreased in the mucosa of DSS-treated mice suggesting an anti-inflammatory or pro-resolving state (25, 26). A corresponding increase in the ileum of the AA derivative lipoxin A4 (LXA_4_), a pro-resolving anti-inflammatory eicosanoid, was noted alongside the decreased inflammation (27, 28). LXA_4_ is produced instead of LTB_4_ after neutrophils are exposed to prostaglandins via the induction of 15-LOX, acting as a signal to stop further neutrophil recruitment (29). The presence of LXA_4_ indicates that, in contrast to the colon, the ileum is likely resolving inflammation with the alternate AA derivatives produced being crucial to initiation of this process.

The steroid hormone biosynthesis and primary bile acid biosynthesis pathways were additionally enriched in the ileum post-DSS treatment and these pathways both utilize 25-hydroxycholesterol and dihydroxy-cholesterols. 25-hydroxycholesterol has dual functionality, exerting both pro- and anti-inflammatory effects (30, 31). It enhances B cell secretion of IL-8 and IL-6 and its production correlates with increased cytokine C-C Motif chemokine Ligand 5 expression (CCL5), the production of which promotes immune cell infiltration. However, as our data does not show the ileal mucosa was overcome with immune cell infiltrate, it appears likely that here 25-hydroxycholesterol is exerting a complementary effect to LXA_4_ whereby it decreases production of the proinflammatory cytokine IL-1β, thus reducing the recruitment and retention of macrophages (31). It is likely that the significant decrease in macrophage subsets observed in the ileal mucosa here may be due to these 25-hydroxycholesterol-driven attempts at resolution of inflammation.

In the inflamed colonic mucosa cells that were proliferative or hypoxic were significantly reduced alongside antigen-presenting dendritic cells, whereas helper T cells, regulatory T cells, NK cells, neutrophils and macrophages were all increased. NK cells, macrophages and neutrophils play roles in CD progression by producing large quantities of proinflammatory cytokines including TNF-α, IL-6 and IL-8 (32, 33). Moreover, antigen-presenting dendritic cells are pivotal in balancing tolerance and active immunity towards commensals, thus their decreased presence in the inflamed mucosa may be contributing to exacerbation of inflammation here.

The differences in cellular function between the colon and ileum during DSS-induced colitis that were observed here were likely influenced by specific identified metabolites. Creatine, DHA and 1-MNA were studied in detail here as these were significantly increased in DSS-treated mice and were significant contributors to the separation of the clusters of DSS-treated mice from controls. DHA is a precursor of anti-inflammatory docasanoids such as resolvins (Rv), protectins (PD) and maresins (Ma), the anti-inflammatory properties of which reduce production of pro-inflammatory AA-derived prostaglandins and leukotrienes promoting resolution of inflammation (34, 35). While DHA also exerts pro-inflammatory effects, the outcome of DHA presence in the intestine is dependent on whether it is degraded to its anti-inflammatory end products. If unmetabolized DHA accumulates in the intestine it can contribute to inflammation, something which needs to be fully considered given it has been suggested as an IBD therapeutic. Here increased DHA levels in the ileum correlated with a decrease in helper T cells, dendritic cells, and M2b and M1 macrophages, while DHA increased splenic immune cell secretion of IFN-γ *in vitro*. Therefore, with aberrant production of IFN-γ a major driver of IBD, DHA is likely contributing to an excessive immune response resulting in mucosal infiltration and damage. Thus, unmetabolized DHA may activate activation of IFN-γ production and be a major contributor to intestinal pathology (36). The molecule putatively identified as epoxydocosapentaenoic acid (EDP) was increased in the ileum, colon, and spleen of colitis mice. This is also a naturally occurring bioactive derivative of DHA, derived via cytochrome P450 epoxygenases (37). To our knowledge, research has largely overlooked this molecule as focus has been on DHA itself, but EDP also has significant anti-inflammatory potential (37, 38). EDP is increased in the circulation of breast cancer patients following DHA supplementation and has a positive effect on patient prognosis (39, 40). This has been attributed to EDP ability to inhibit angiogenesis, suppress endothelial cell migration, and primary tumour growth (41). Furthermore, EDP can reduce the expression of TNF-α and TNF-α-induced leukocyte adhesion in Müller cells and retinal endothelial cells, thus it may be useful in treating inflammatory eye conditions (38). Therefore, the increased EDP levels in the intestine and spleen may exert protective anti-inflammatory effects and EDP warrants further investigation as an intestinal and systemic inflammatory therapeutic target.

The creatine phosphate shuttle is increased in epithelial cells during DSS-induced metabolic stress, so the observed increase in creatine levels in the DSS-treated ileum observed here may be a metabolic response to restore cellular energy (42). Creatine supplementation in DSS colitis reduces disease severity and improves barrier function, thus the use of creatine in IBD patients has been suggested (43). Creatine supplementation also reduces the secretion of TNF-α from macrophages, dampening the M1 inflammatory response (44). Our data loosely supports this idea as IMC revealed a reduction in cells with a macrophage-like phenotype in the ileum where creatine was increased. The third metabolite tested, 1-MNA, also has been suggested to be anti-inflammatory exerting a hepatoprotective effect in liver inflammation by suppressing IL-4 and TNF-α immune cell signalling (45, 46). 1-MNA was seen here to reduce production of IFN-γ, something previously described in CD4^+^ T cells (47). CD4^+^ T cell production of IFN-γ however is essential for bactericidal activity against many microbes (48). So, 1-MNA suppression of IFN-γ may be a double-edged sword, reducing inflammation but resulting in impaired bacterial clearance which could further contribute to dysbiosis.

Primary bile acid biosynthesis was enriched in the ileum of DSS-treated mice but bile acid pathways in the liver were surprisingly unaffected with galactose metabolism the most significantly enriched pathway. Galactose is hydrolysed from ingested lactose in the intestine and travels to the liver where it is metabolised exclusively (49). Glucose transporter 1 (Glut1), was significantly increased in the liver of colitic mice, indicative of increased glucose needs due to stress and a likely reliance on D-galactose metabolism for liver energy (50–52). The pathway for biosynthesis of unsaturated fatty acids pathway was also enriched in the liver reflecting a decrease in the abundance of gamma-linolenic acid (GLA) in the liver, ileum, colon, and kidney of colitis mice. GLA is rapidly enzymatically converted into dihomo-gamma-linolenic acid (DGLA) and due to continuous conversion, it is naturally found at low levels with a corresponding increase in DGLA (53). Liver cells, have the capacity to desaturate DGLA to form AA and AA was increased in the liver of colitis mice suggesting that GLA and DGLA were used as precursors for AA production (54, 55). Furthermore, this study putatively identified prostaglandin G2, an AA derivative, as being increased in the liver during colitis and has been implicated as a pathophysiological factor in liver cirrhosis by promoting platelet aggregation and thrombosis (56). Therefore, the bidirectional liver-gut axis may exacerbate inflammation and tissue damage in both organs via production of AA and its derivatives. However, the disappearance of precursor GLA under these conditions could prove to be a useful biomarker of systemic disease progression (55, 57).

Other metabolites were putatively identified that could potentially be biomarkers of disease and warrant further investigation. Ubiquinone-2, also known as coenzyme Q2 (COQ2), is involved in the biosynthesis of endogenous coenzyme Q10 (CoQ10) and was increased in the ileum, colon, liver, and spleen of DSS-treated mice (58). Within mitochondria, CoQ10 acts as an electron carrier that increases the generation of ATP via oxidative phosphorylation (59). In addition, CoQ10 can be reduced to ubiquinol, which strong antioxidant and anti-inflammatory properties (60). During LPS-induced stress, ubiquinol treatment blocks protein kinase C (PKC) activity and improves mitochondrial function, preventing ROS generation and cellular damage (61). CoQ10 supplementation in ulcerative colitis patients, also decreased the severity of inflammation and reduces blood pressure (62). As ubiquinone-2 was increased in the intestine, liver, and spleen, it can be suggested that increasing biosynthesis or supplementation would not only reduce inflammation in the intestine but could have a systemic protective effect given the localisation observed here, increasing its usefulness as a therapeutic target (61–63).

In conclusion, spatial biology approaches were employed to discover metabolites derived by the host and microbiome that could play a role in the onset and progression of DSS-induced colitis. These spatial metabolomic approaches were complemented by determining the identity and phenotypes of cells in the same organs, both in the intestine and systemically. Our study revealed numerous molecular pathways of interest and specific molecules, such as creatine, DHA and 1-MNA, that have immunomodulatory effects and have the potential to perpetuate inflammation within the intestinal mucosa. Furthermore, the study shows that specific metabolites are altered in the liver, spleen and kidney during inflammation and have the potential to drive pro and anti-inflammatory mechanisms. Some of these metabolites such as GLA may result in systemic disease manifestations and may be new targets for therapeutic intervention. Lastly, some metabolites such as ubiquinone-2 and EDP are present in multiple tissue including the intestine during colitis; thus, if used as therapeutic target could dampen both intestinal and systemic inflammation.

## Supporting information

Adams et al. Supplementary Materials

