## Supplementary material for "Spatial mapping of dextran sodium sulphate-induced intestinal inflammation and its systemic effects": Adams et al. Supplementary Materials

**Supplementary Figure 1**

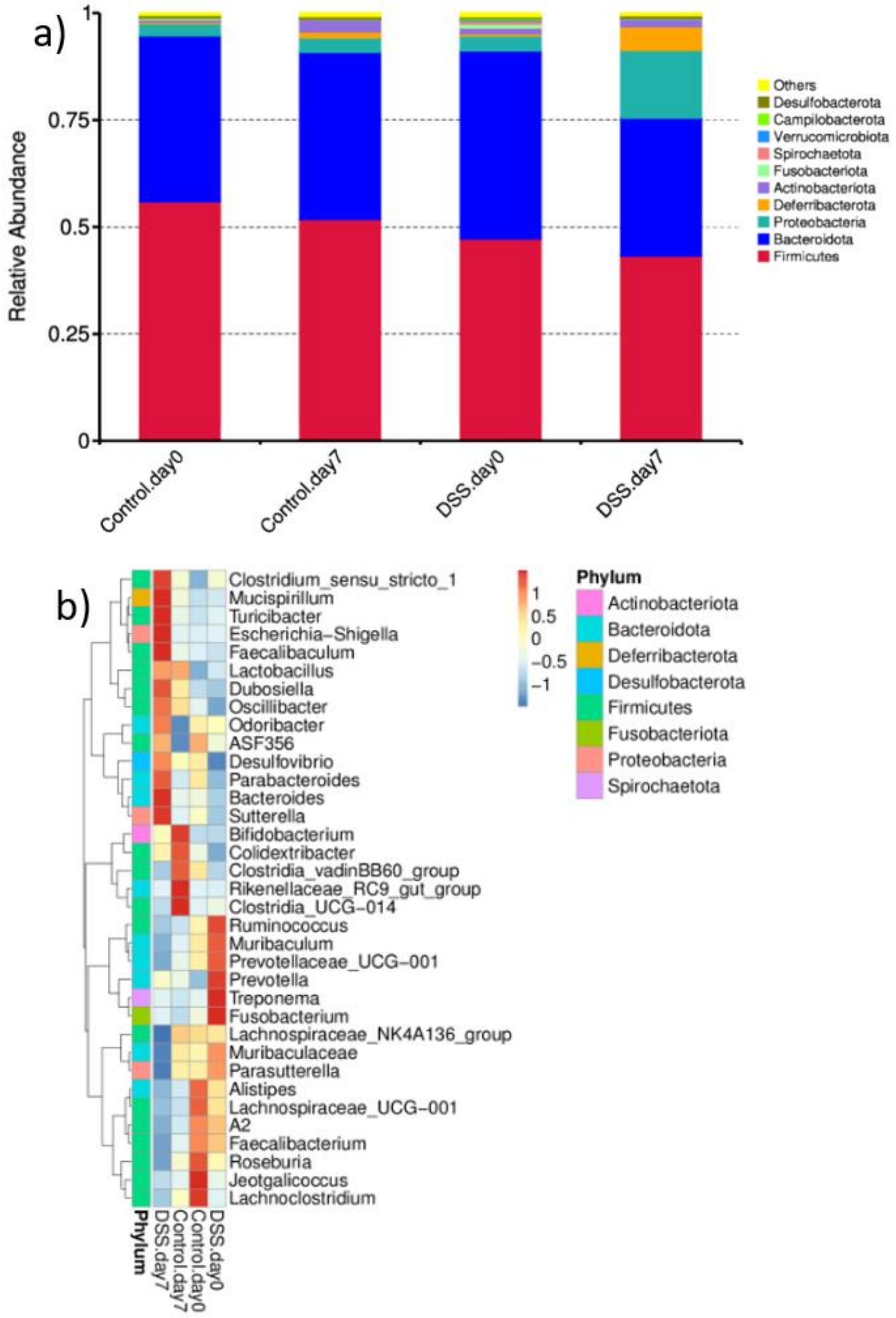


**Figure S1. Relative abundance of bacterial phyla and heatmap of genera in 3% DSS treated mice and controls.** a) Histogram of relative abundance of the top 10 bacterial phyla in the faeces of control mice (0% DSS) and 3% DSS treated mice at the start (day 0) and end (day 7) of experiment. b) Heatmap of top 35 genera in control groups and 3% DSS treated mice on day 0 and day 7.

**Supplementary Figure 2**

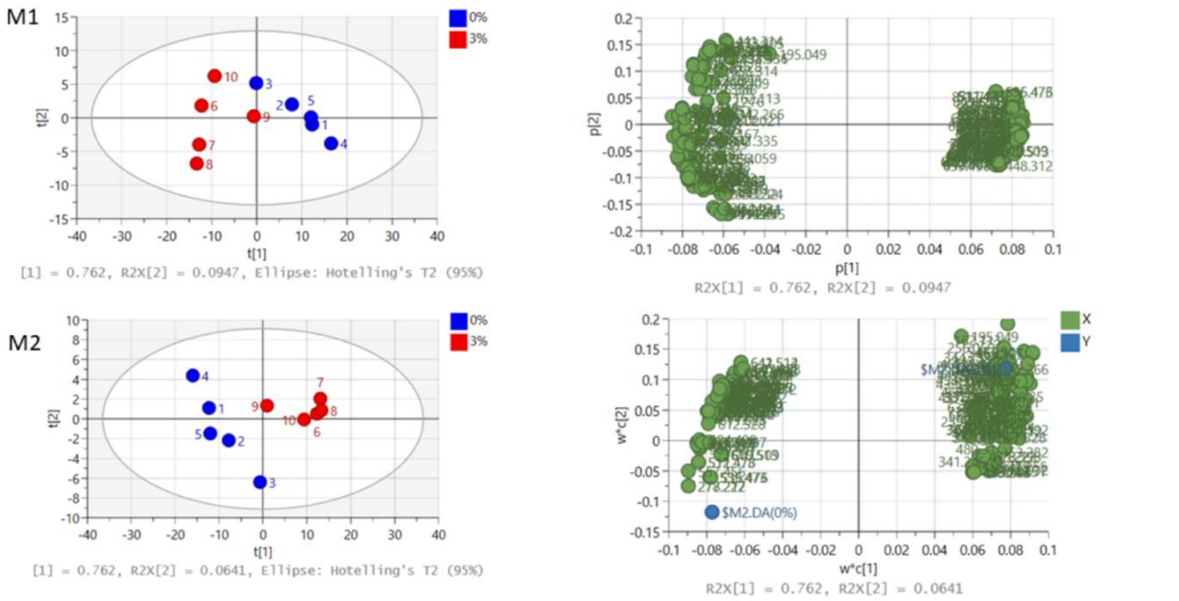


| Number | Model | Type | A | N | R2X(cum) | R2Y(cum) | Q2(cum) |
|---|---|---|---|---|---|---|---|
| 1 | M1 | PCA-X | 2 | 10 | 0.857 | | 0.754 |
| 2 | M2 | PLS-DA | 2 | 10 | 0.826 | 0.907 | 0.788 |

**Figure S2. Unsupervised and supervised discriminant analysis of the ileum of 3% DSS treated mice and controls.** M1) Unsupervised PCA analysis and M2) Supervised PLS-DA analysis show the molecules in the ileum of mice can discriminate between the control (0% DSS-blue circles) and treated group (3% DSS-red circles). Analysis was performed using SIMCA 17 software. PLS-DA and PCA score plots consist of components 1 (t [1]) and 2 (t [2]). The ellipse represents the 95% confidence region for Hotelling's T2 statistic for the model.

**Supplementary Figure S3**

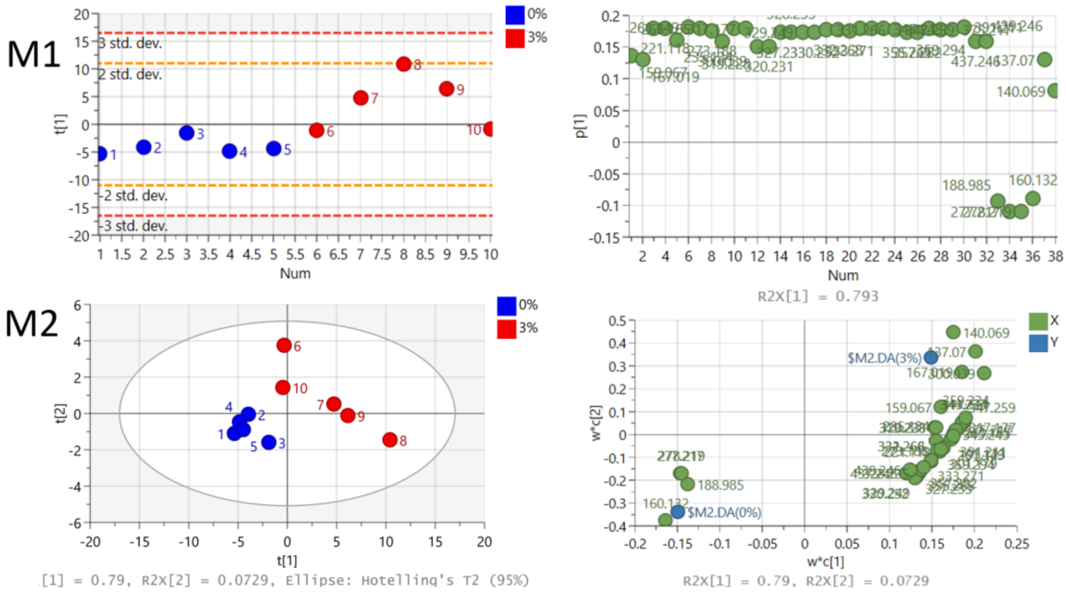


| Model | Type | A | N | R2X(cum) | R2Y(cum) | Q2(cum) |
|---|---|---|---|---|---|---|
| M1 | PCA-X | 1 | 10 | 0.793 | | 0.727 |
| M2 | PLS-DA | 4 | 10 | 0.946 | 0.993 | 0.93 |

**Figure S3. Unsupervised and supervised discriminant analysis of the colon of 3% DSS treated mice and controls.** M1) Unsupervised PCA analysis was able to discriminate between the groups using metabolite features. M2) Supervised PLS-DA analysis show the molecules in the colon of mice can discriminate between the control (0% DSS, blue circles) and treated groups (3% DSS, red circles). Analysis was performed using SIMCA 17 software. Analysis was performed using SIMCA 17 software. PLS-DA and PCA score plots consist of components 1 (t [1]) and 2 (t [2]). The ellipse represents the 95% confidence region for Hotelling's T2 statistic for the model.

**Supplementary Figure 4**

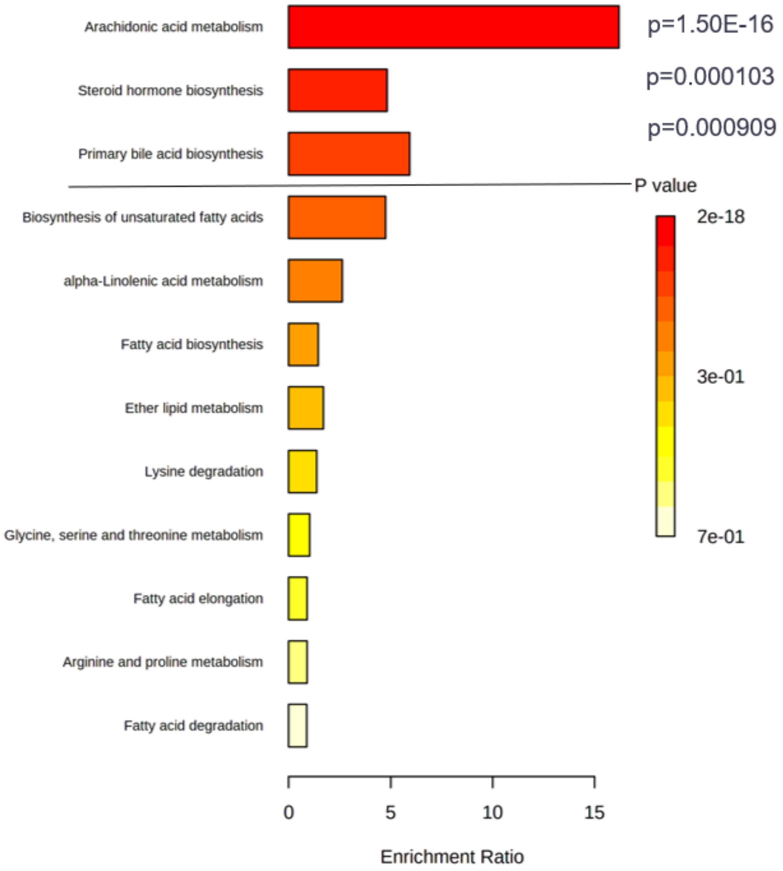


**Figure S4. Enrichment of pathways in the ileum of DSS colitis model.** Enrichment pathway analysis using KEGG as reference found molecules involved in 12 different pathways: arachidonic acid metabolism, steroid hormone biosynthesis, primary bile acid biosynthesis, biosynthesis of unsaturated fatty acids, alpha-linolenic acid metabolism, fatty acid biosynthesis, ether lipid metabolism, lysine degradation, glycine, serine and threonine metabolism, fatty acid elongation, arginine and proline metabolism and fatty acid degradation. The top 3 pathways were significantly enriched.

**Supplementary Figure 5**

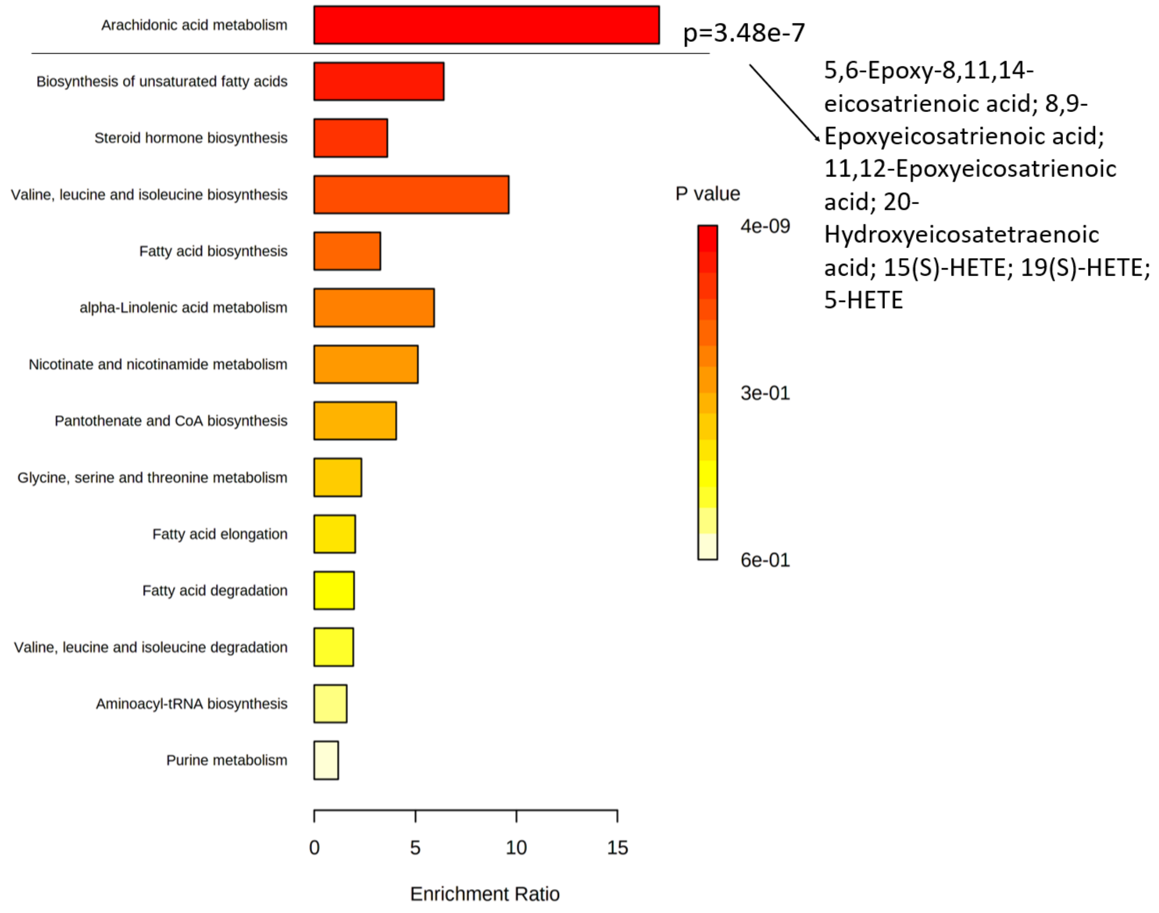


**Figure S5. Enrichment of pathways in the colon of DSS colitis model.** Enrichment pathway analysis using KEGG as reference found molecules involved in 14 different pathways; arachidonic acid metabolism, biosynthesis of unsaturated fatty acids, steroid hormone biosynthesis, valine, leucine and isoleucine biosynthesis, fatty acid biosynthesis, alpha-Linolenic acid metabolism, nicotinate and nicotinamide metabolism, pantothenate and CoA biosynthesis, glycine, serine and threonine metabolism, fatty acid elongation and degradation, valine, leucine and isoleucine degradation, aminoacyl-tRNA biosynthesis and purine metabolism. The top pathway was significantly enriched, and the molecules involved in the pathway that are present in the dataset are listed.

**Supplementary Figure 6**

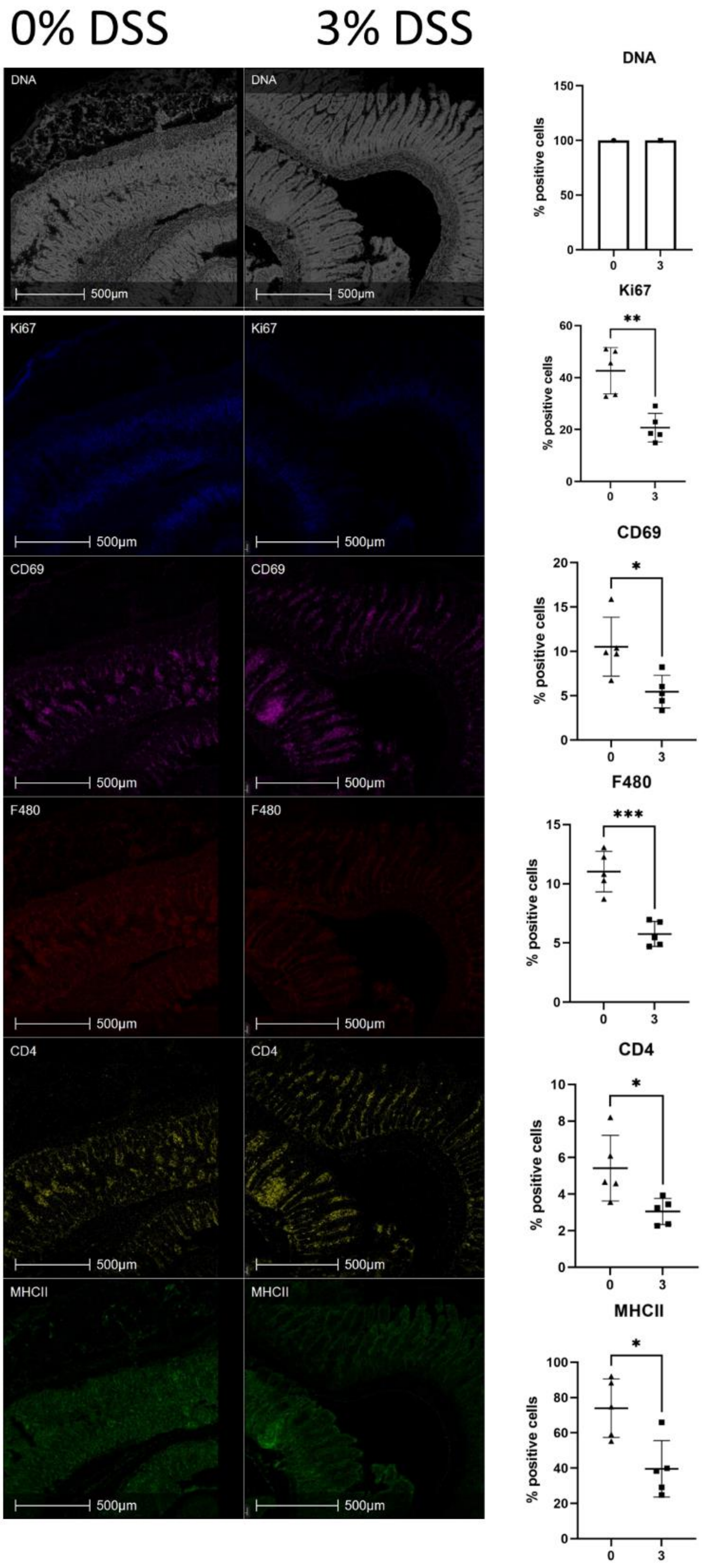

**Figure S6. Representative IMC images and percentage of cells positive for biological markers of cell function in the ileum of DSS colitis model.** Image shown is the region indicated in Fig 2. From top to bottom, DNA intercalator identified single cells within the tissue section, markers for cell proliferation and immune cell function are shown (fold change and *p* value); Ki67 (2.05-fold, $p$=0.0016), CD69 (1.96-fold, $p$=0.0176), F480 (2.0-fold, $p$=0.0004), CD4 (1.77-fold, $p$=0.0252) and MHCII (1.86-fold, $p$=0.0105). Percentage positive cells represented as bar graph showing five biological replicates. A t-test was performed to compare the two groups and *$p<0.05$, **$p<0.01$, ***$p<0.001$, ****$p<0.0001$ were considered statistically significant.

**Supplementary Figure 7**

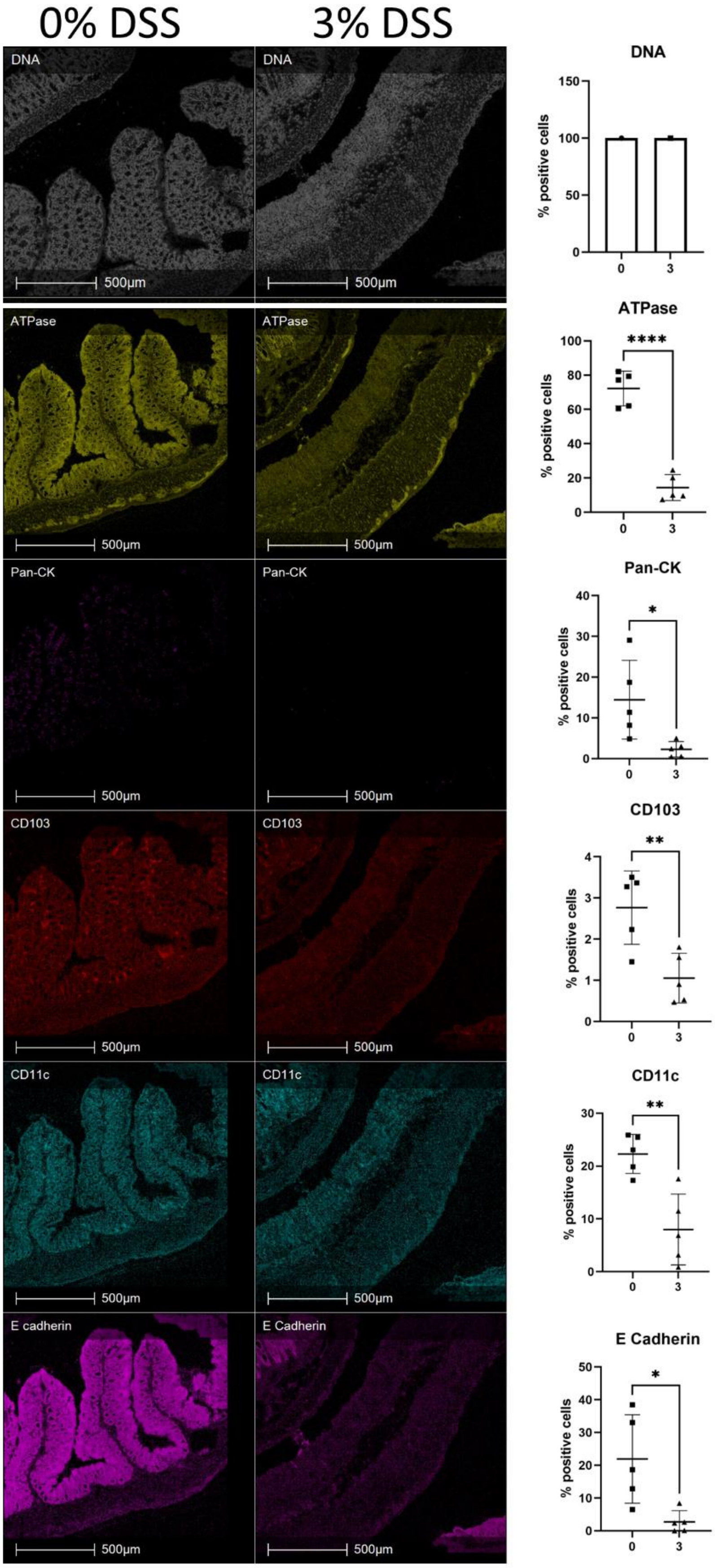

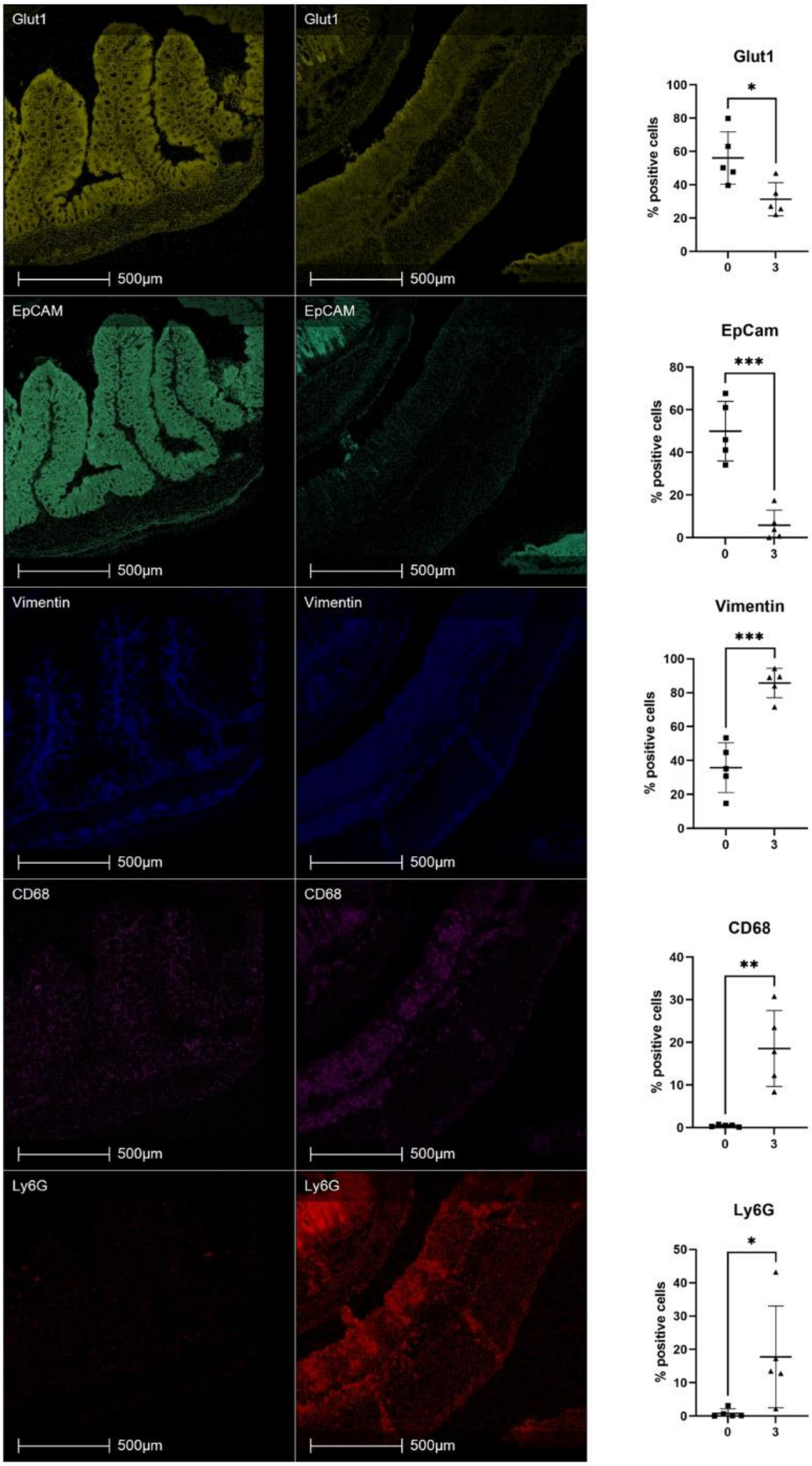
Glut1
Glut1
500µm
500µm
EpCAM
EpCAM
500µm
500µm
Vimentin
Vimentin
500µm
500µm
CD68
CD68
500µm
500µm
Ly6G
Ly6G
500µm
500µm
Glut1
% positive cells
*
EpCam
% positive cells
***
Vimentin
% positive cells
***
CD68
% positive cells
**
Ly6G
% positive cells
*
0
3

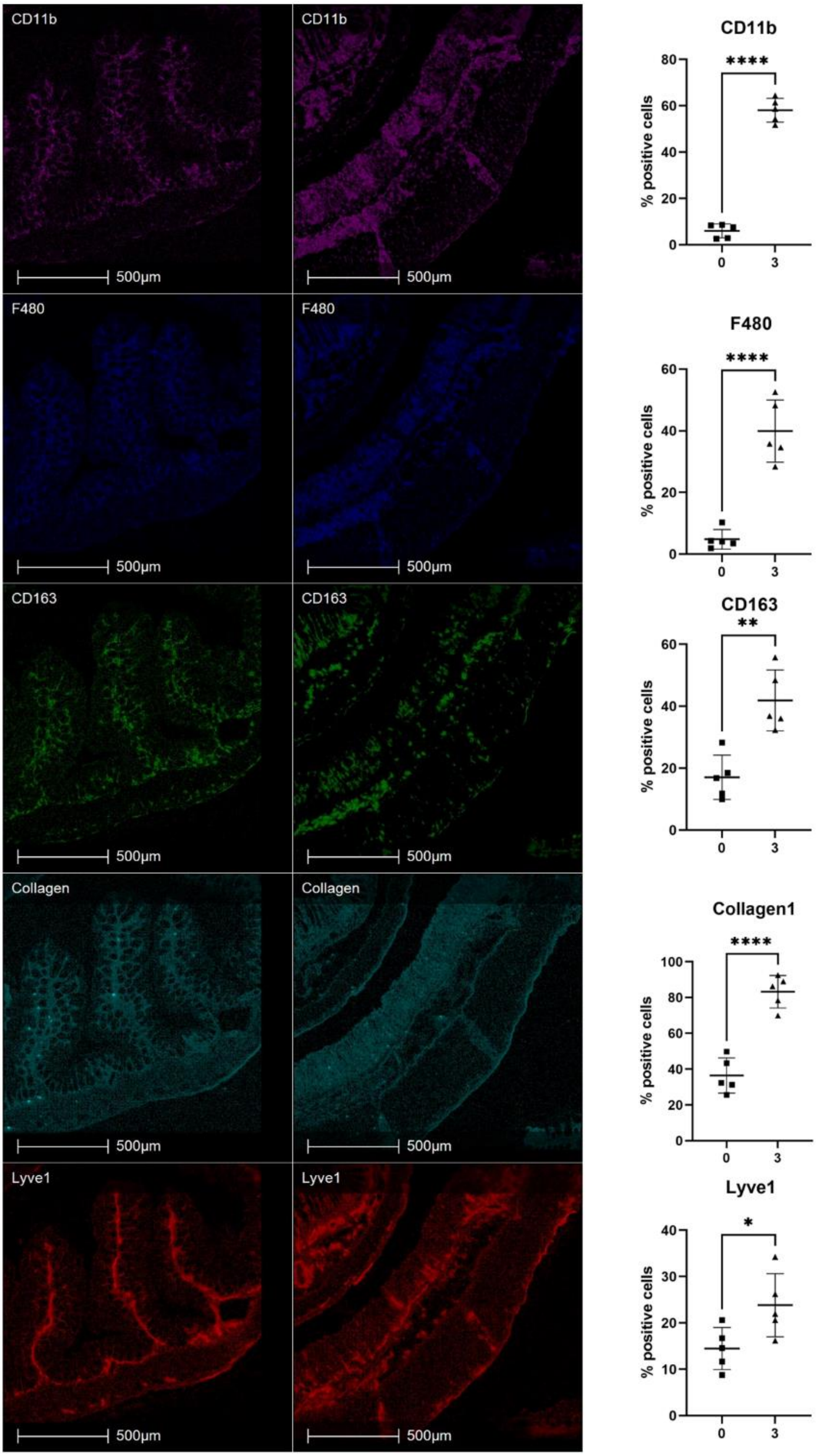
CD11b
500µm
CD11b
500µm
F480
500µm
F480
500µm
CD163
500µm
CD163
500µm
Collagen
500µm
Collagen
500µm
Lyve1
500µm
Lyve1
500µm
CD11b
****
% positive cells
0
3
F480
****
% positive cells
0
3
CD163
**
% positive cells
0
3
Collagen1
****
% positive cells
0
3
Lyve1
*
% positive cells
0
3

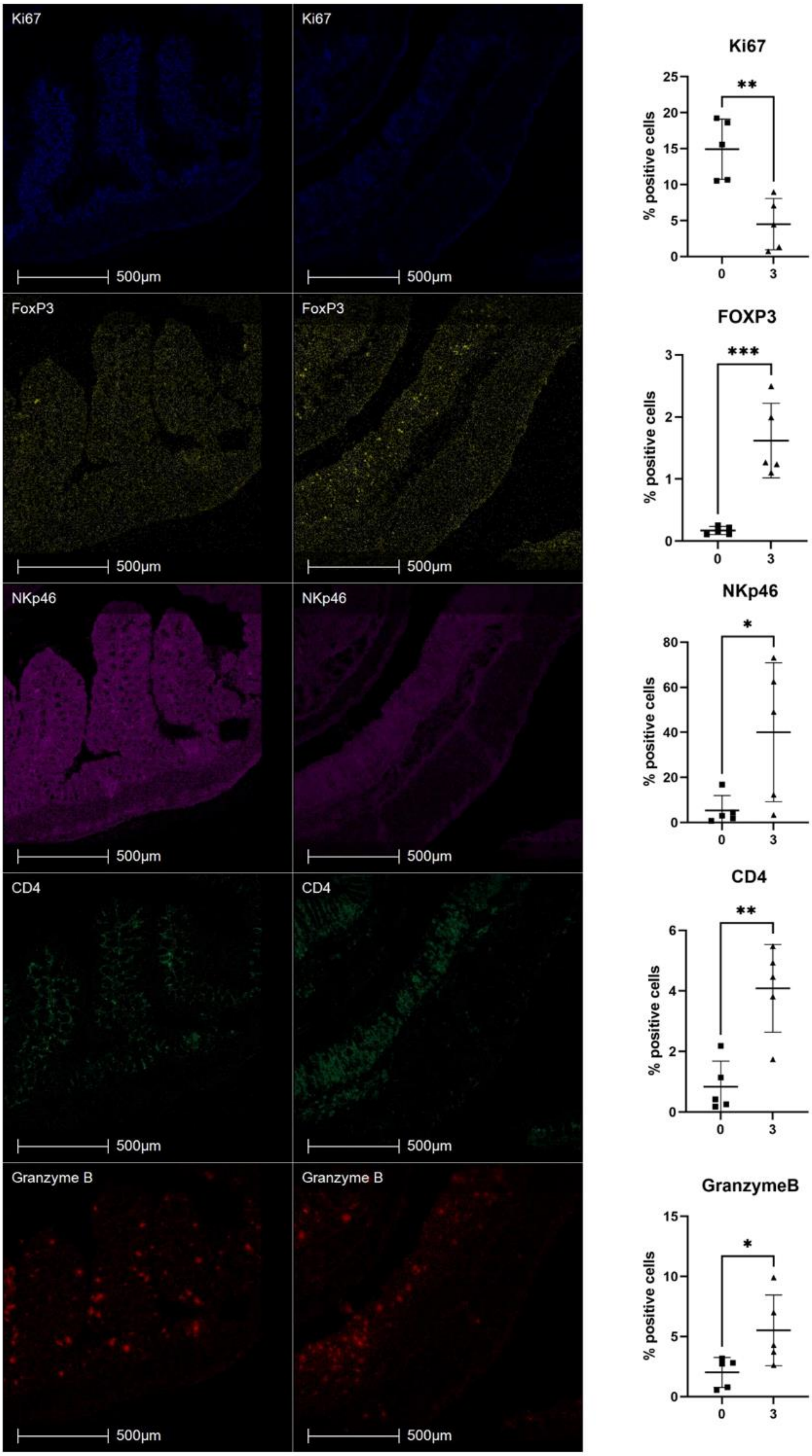
Ki67
Ki67
500µm
500µm
FoxP3
FoxP3
500µm
500µm
NKp46
NKp46
500µm
500µm
CD4
CD4
500µm
500µm
Granzyme B
Granzyme B
500µm
500µm
Ki67
**
% positive cells
0
3
FOXP3
***
% positive cells
0
3
NKp46
*
% positive cells
0
3
CD4
**
% positive cells
0
3
GranzymeB
*
% positive cells
0
3

**Figure S7. Representative IMC images and percentage of cells positive for biological markers of cell function in the colon of DSS colitis mice.** Image shown is the same region as in Fig. 2. From top to bottom, DNA intercalator identified single cells within the tissue section, markers for cell proliferation and immune cell function ATPase, Pan-CK, CD103, CD11c, E Cadherin, Glut1, EpCAM, Vimentin, CD68, Ly6G, CD11b, F480, CD163, Collagen1, Lyve1, Ki67, FOXP3, NKp46, CD4 and Granzyme B. Percentage positive cells represented as bar graph showing five biological replicates. A t-test was performed to compare the two groups and *$p<0.05$, **$p<0.01$, ***$p<0.001$, ****$p<0.0001$ were considered statistically significant. Markers that decreased included ATPase 5.01-fold ($p<0.0001$), Pan-CK 6.24-fold ($p=0.0246$), CD103 2.62-fold ($p=0.0074$), CD11c 2.62-fold ($p=0.0074$), E Cadherin 2.79-fold ($p=0.0030$), Glut1 1.78-fold ($p=0.0177$), Ki67 3.3-fold ($p=0.0028$) and EpCAM 3.30-fold ($p=0.0028$). In contrast the following markers increased significantly post-DSS treatment; vimentin 2.39-fold ($p=0.0002$), CD68 39.18-fold ($p=0.0019$), Ly6G 22.13-fold ($p=0.0382$), CD11b 9.64-fold ($p<0.0001$), F480 8.30-fold ($p<0.0001$), CD163 2.45-fold ($p=0.0018$), collagen 2.28-fold ($p<0.0001$), Lyve1 1.64-fold ($p=0.0342$), FoxP3 9.50-fold ($p=0.0007$), NKp46 7.42-fold ($p=0.0393$), CD4 4.88-fold ($p=0.0025$), and granzyme B 2.72-fold ($p=0.0402$).

**Supplementary Figure 8**

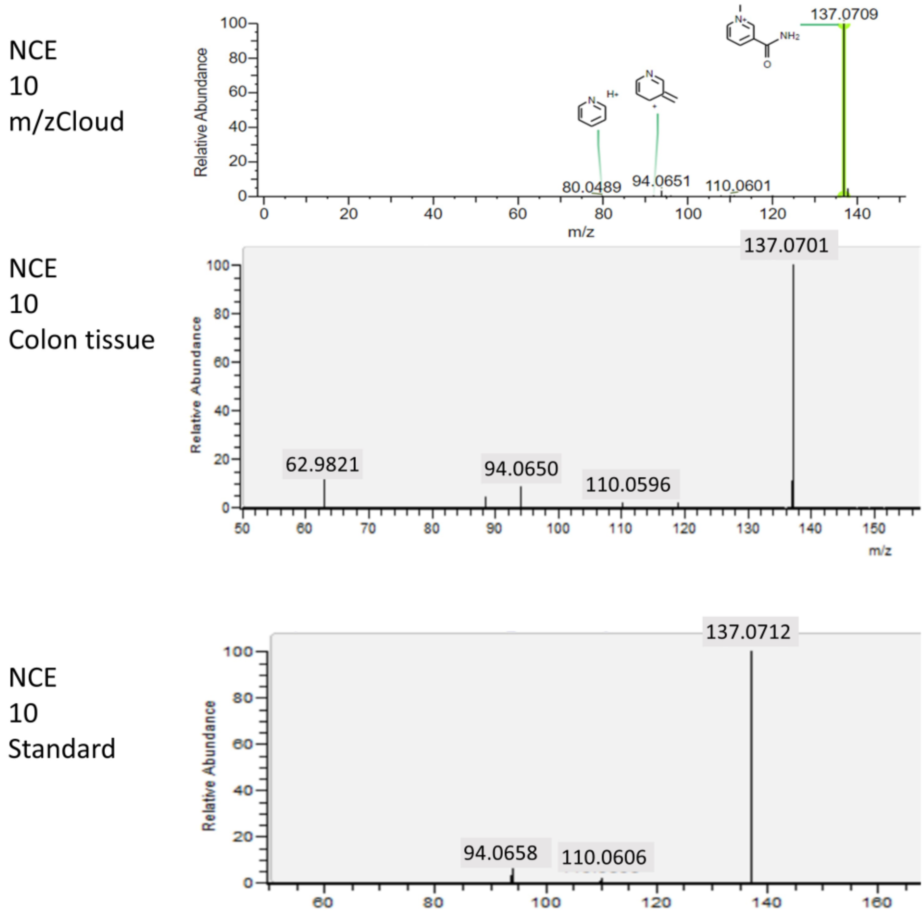


**Figure S8A. MSMS spectra for 1-MNA standard alongside tissue spectra.** MSMS fragmentation pattern (*mz* plotted against relative abundance) using NCE 10 shown in mzCloud, colon tissue and 1-MNA purchased standard. Peaks at 137.07, 110.06, and 94.06 are of similar abundance in all three fragmentation patterns; thus, 137.07 in colon tissue is most likely to be 1-MNA.

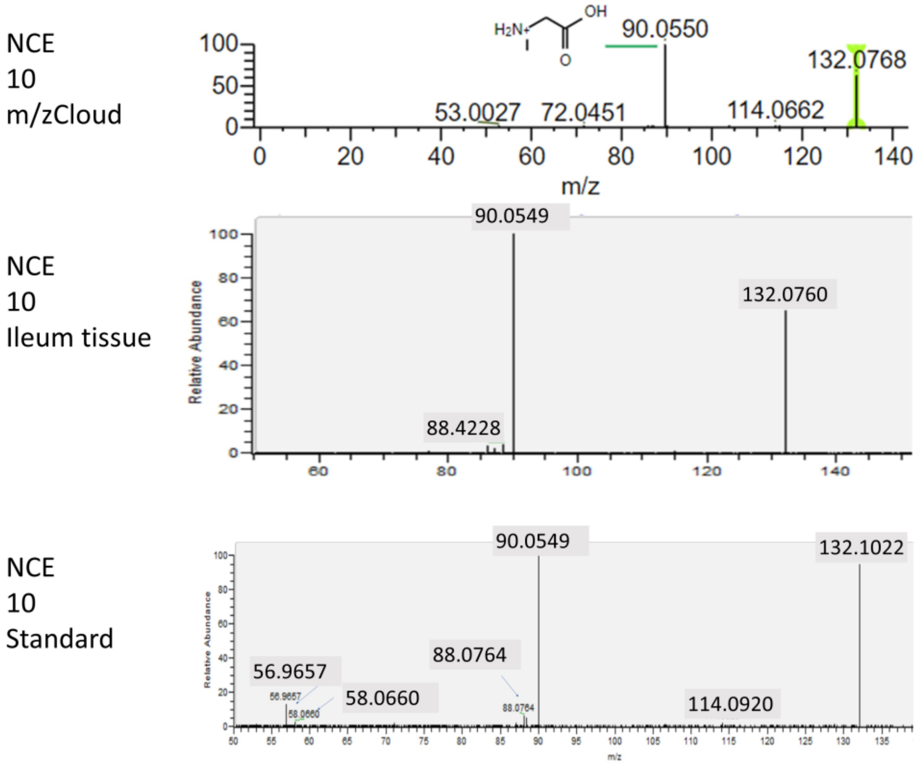


**Figure S8B. MSMS spectra for creatine standard alongside tissue spectra.** MSMS fragmentation pattern (*m/z* plotted against relative abundance) using NCE 10 shown in mzCloud, ileum tissue and a creatine purchased standard. Peaks at 132.07 and 90.05 are of similar abundance in all three fragmentation patterns; thus, 132.07 in ileum is most likely to be creatine.

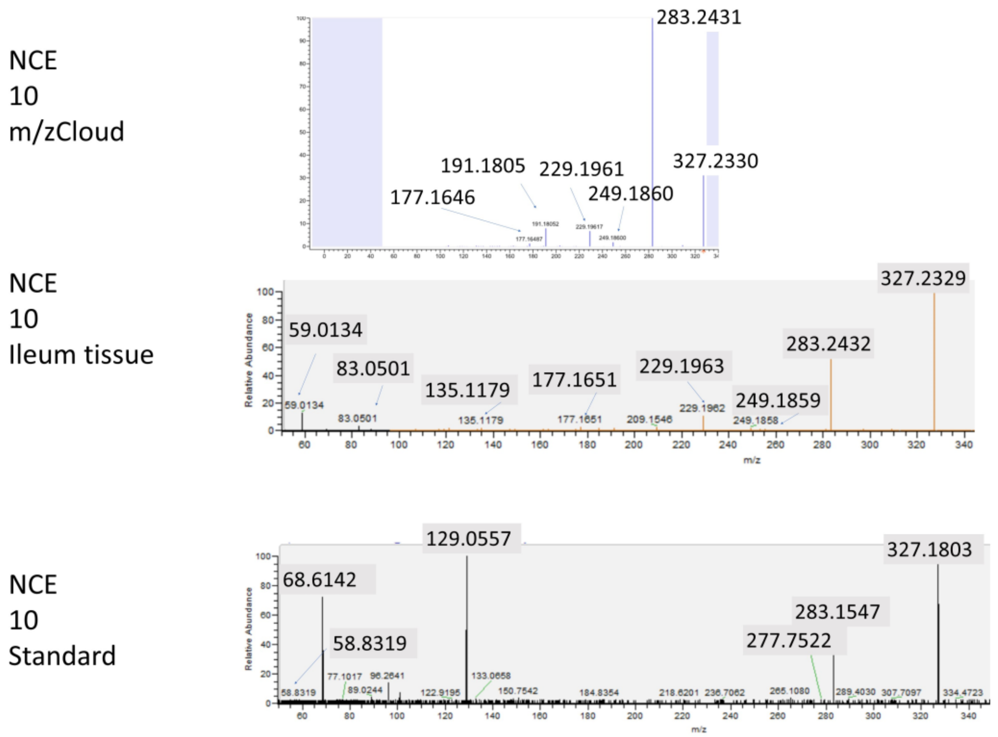


**Figure S8C. MSMS spectra for DHA standard alongside tissue spectra.** MSMS fragmentation pattern (m/z plotted against relative abundance) using NCE 10 shown in m/z cloud, ileum tissue and a DHA purchased standard. Peaks at 327.23, 283.24, 249.18, 229.19 and 177.16 are of similar abundance in mzCloud and ileum fragmentation patterns; thus, the molecule was putatively identified as DHA.

**Supplementary Figure 9**

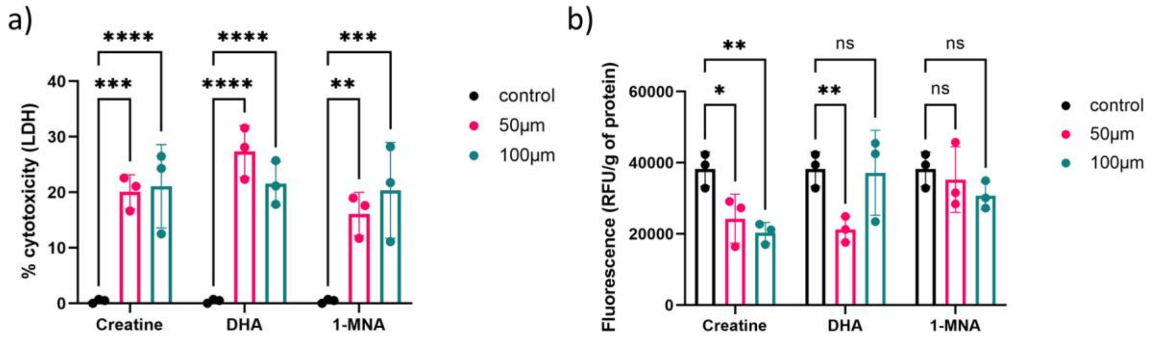


**Figure S9. LDH release and caspase-3/7 activity from HCT-8 cells treated with molecules of interest.** HCT-8 cells were exposed to Creatine, DHA and 1-MNA for 72 h. a) LDH assay used HCT-8 cells alone and cells treated with 2% triton-x as low and high LDH release controls, respectively. The percentage of cytotoxicity was calculated as %= (measured absorbance of sample-low control)/ (high control-low control) x100. b) The caspase-3/7 assay calculated enzyme activity by measuring relative fluorescence units (RFU) of activity normalised to protein concentration in cell lysates. Data are shown as the mean of three biological replicates ± standard deviation (SD) (error bars). Two-way ANOVA was performed across to compared metabolite exposure to cell only controls *$p$<0.05, **$p$<0.01, ***$p$<0.001, ****$p$<0.0001 versus the control condition (cells without molecules) was considered statistically significant.

**Supplementary Figure 10**

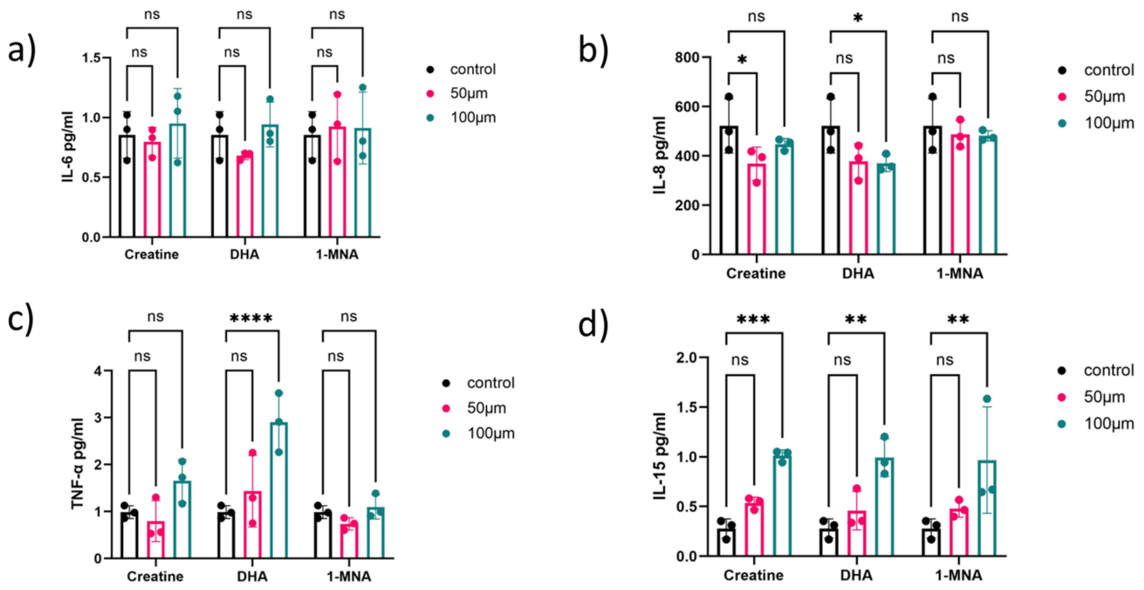


**Figure S10. Cytokine release into cell supernatant after 72h exposure to metabolites**. HCT-8 cells were stimulated with creatine, DHA and 1-MNA for 72 h before supernatants were collected. ELISA was used to quantify the release of a) IL-6 pg/ml, b) IL-8 pg/ml, c) TNF-α pg/ml and d) IL-15 pg/ml. Data is shown as the mean of 3 biological replicates ± standard deviation (SD). Two-way ANOVA was performed for each molecule versus the control condition (cells without treatment). *$p$<0.05, **$p$<0.01, ***$p$<0.001, ****$p$<0.0001 is considered statistically significant.

**Supplementary Figure 11**

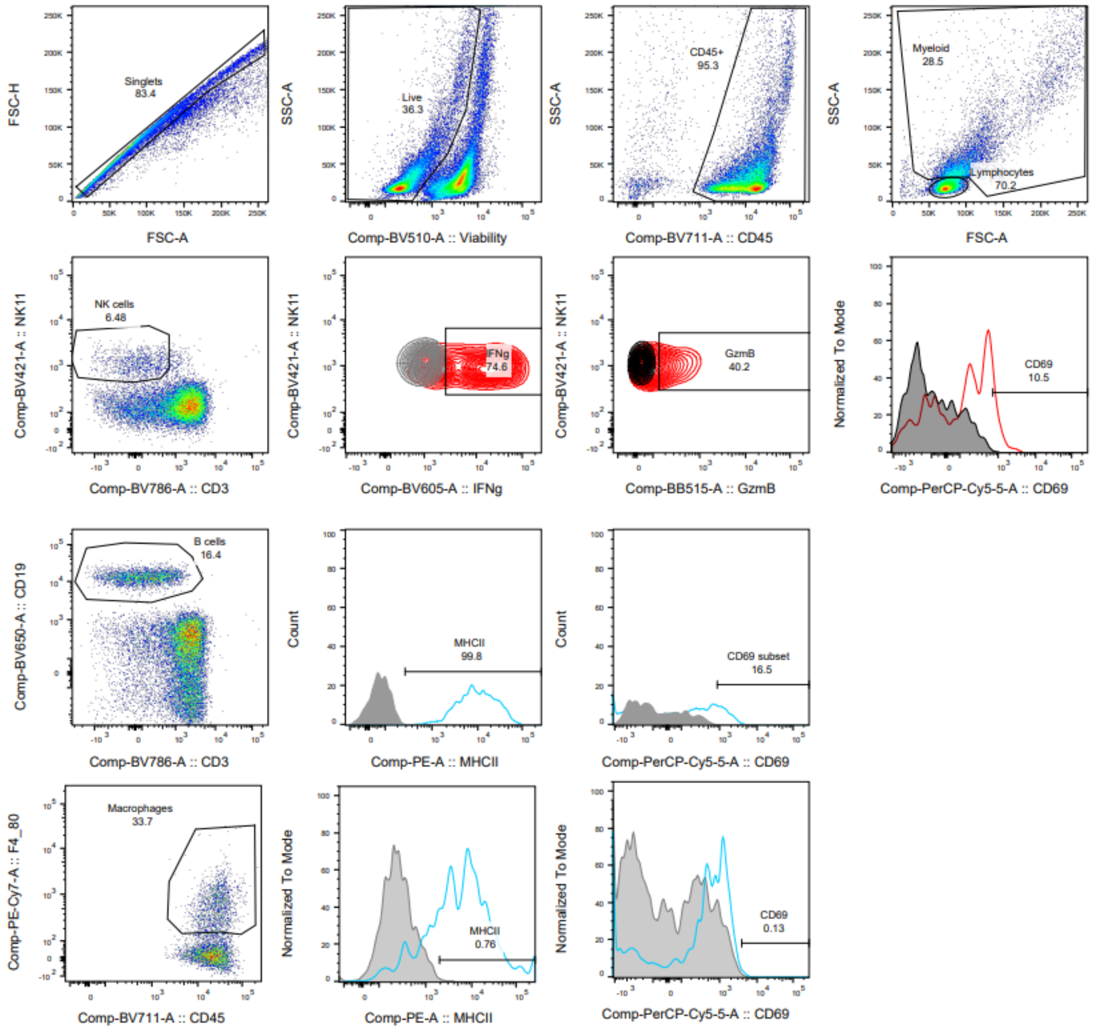


**Figure S11: Gating strategy for isolation of splenic cells for *ex vivo* analysis.** Cells were gated on FCS-A vs H to exclude duplets. Among singlets dead cells were excluded based on viability staining. Among live cells CD45+ hematopoietic cells were used for analysis. CD45+ cells were split into lymphocytes and myeloid cells based on size (FSC) and granularity (SSC). Lymphocytes were further divided into NK cells (NK1.1+CD3-) and B cells (CD19+CD3-). Macrophages were identified based on F4/80 expression out of myeloid cells. CD69 activation marker was used as a readout of putative activation on both myeloid and lymphoid cells. The figure shows the overlay of actual signal vs isotype control (background) stain (in grey). IFN-γ and Granzyme B levels were assessed in NK cells with positivity being considered based on isotype control staining (overlayed in grey/black). MHCII expression levels were quantified on macrophages and B cells based on isotype control stains for background levels (negative vs positive signal). Percentages of positive or MFI (mean fluorescence intensity) levels were extracted and used to graph results and perform statistical analysis.

**Supplementary Figure 12**

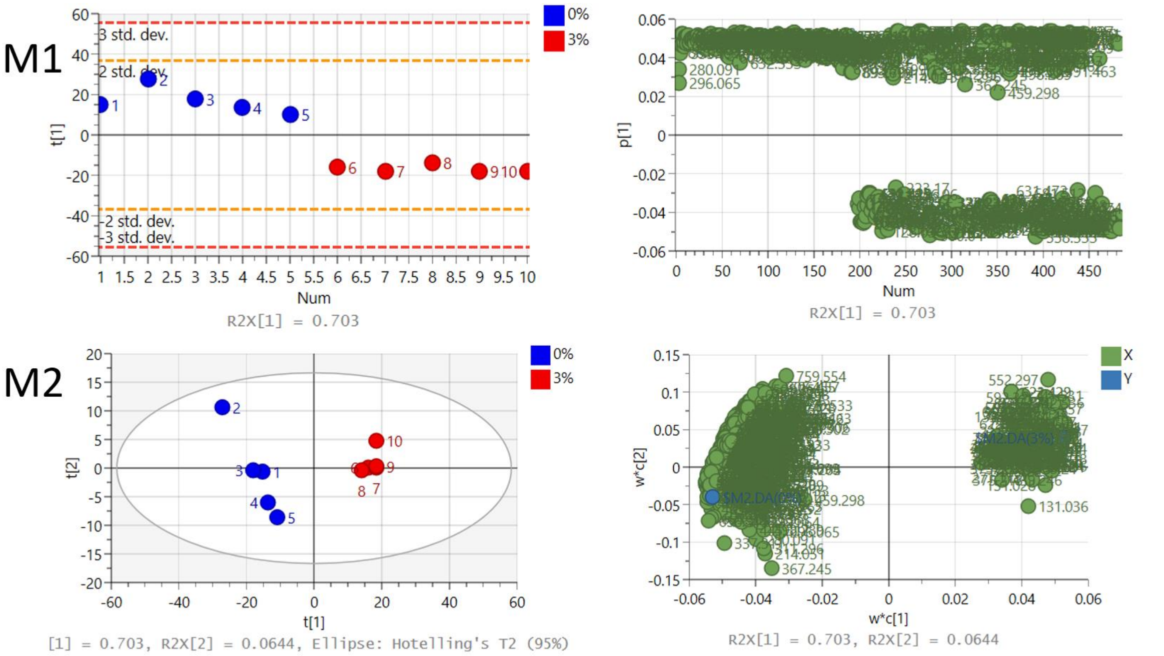


| Model | Type | A | N | R2X(cum) | R2Y(cum) | Q2(cum) |
|---|---|---|---|---|---|---|
| M1 | PCA-X | 1 | 10 | 0.703 | | 0.605 |
| M2 | PLS-DA | 2 | 10 | 0.767 | 0.986 | 0.939 |

**Figure S12. Unsupervised and supervised discriminant analysis of the liver of 3% DSS treated mice and controls.** M1) Unsupervised PCA analysis was able to discriminate between the groups using metabolite features. M2) Supervised PLS-DA analysis show that molecules in the mouse liver can discriminate between the control (0% DSS, blue circles) and treated groups (3% DSS, red circles). Analysis was performed using SIMCA 17 software. PLS-DA and PCA score plots consist of components 1 (t [1]) and 2 (t [2]). The ellipse represents the 95% confidence region for Hotelling’s T2 statistic for the model.

**Supplementary Figure 13**

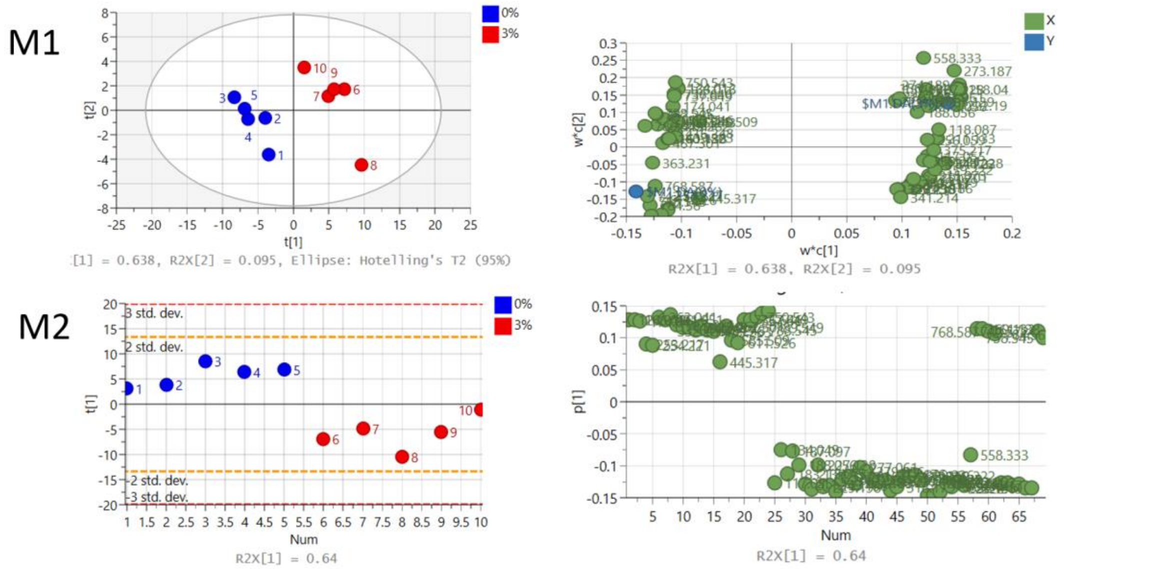


| Model | Type | A | N | R2X(cum) | R2Y(cum) | Q2(cum) |
|---|---|---|---|---|---|---|
| M1 | PLS-DA | 2 | 10 | 0.733 | 0.965 | 0.894 |
| M2 | PCA-X | 1 | 10 | 0.64 | | 0.5 |

**Figure S13. Unsupervised and supervised discriminant analysis of molecules altered in the spleen of control and 3% DSS treated mice.** M1) Supervised PLS-DA analysis show the molecules in the mouse spleen can discriminate between the control (0% DSS, blue circles) and treated group (3% DSS, red circles). M2) Unsupervised PCA analysis was not able to discriminate between the groups using metabolite features. Analysis was performed using SIMCA 17 software. PLS-DA and PCA score plots consist of components 1 (t [1]) and 2 (t [2]). The ellipse represents the 95% confidence region for Hotelling'g T2 statistic for the model.

**Supplementary Figure 14**

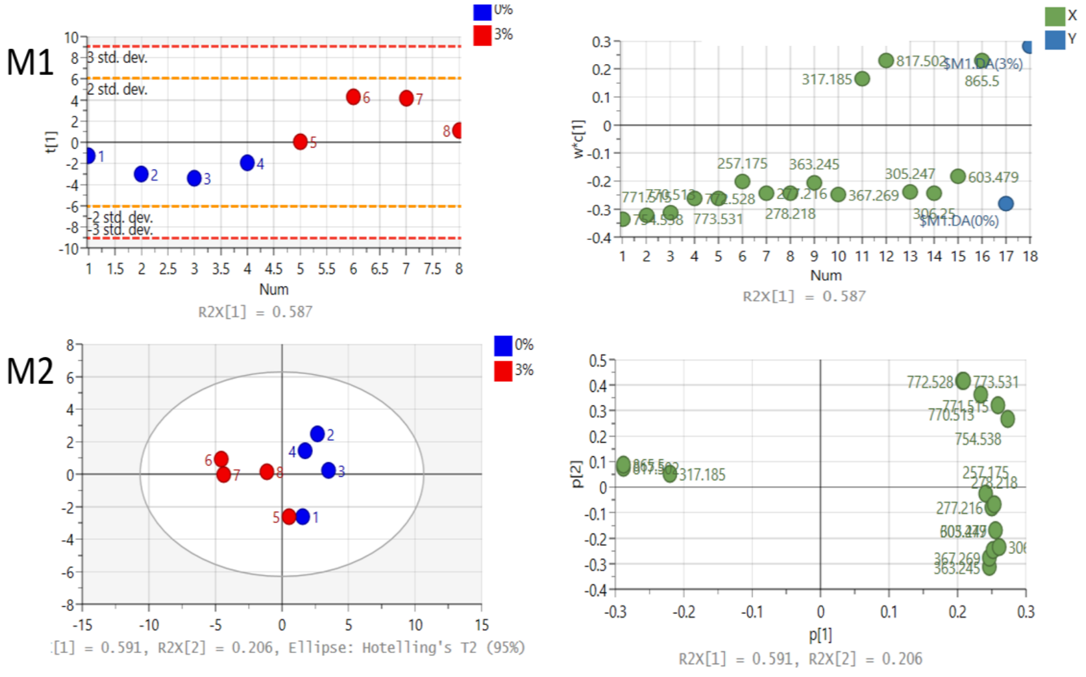


| Model | Type | A | N | R2X(cum) | R2Y(cum) | Q2(cum) |
|---|---|---|---|---|---|---|
| M1 | PLS-DA | 1 | 8 | 0.587 | 0.729 | 0.625 |
| M2 | PCA-X | 3 | 8 | 0.904 | | 0.685 |

**Figure S14. Unsupervised and supervised discriminant analysis of the kidney in 3% DSS treated mice compared to controls.** M1) Supervised PLS-DA analysis show the molecules in the kidneys of mice can discriminate between the control (0% DSS, blue circles) and treated group (3% DSS, red circles). M2) Unsupervised PCA analysis was able to discriminate between the groups using metabolite features. Analysis was performed using SIMCA 17 software. PLS-DA and PCA score plots consist of components 1 (t [1]) and 2 (t [2]). The ellipse represents the 95% confidence region for Hotelling's T2 statistic for the model.

**Supplementary Figure 15**

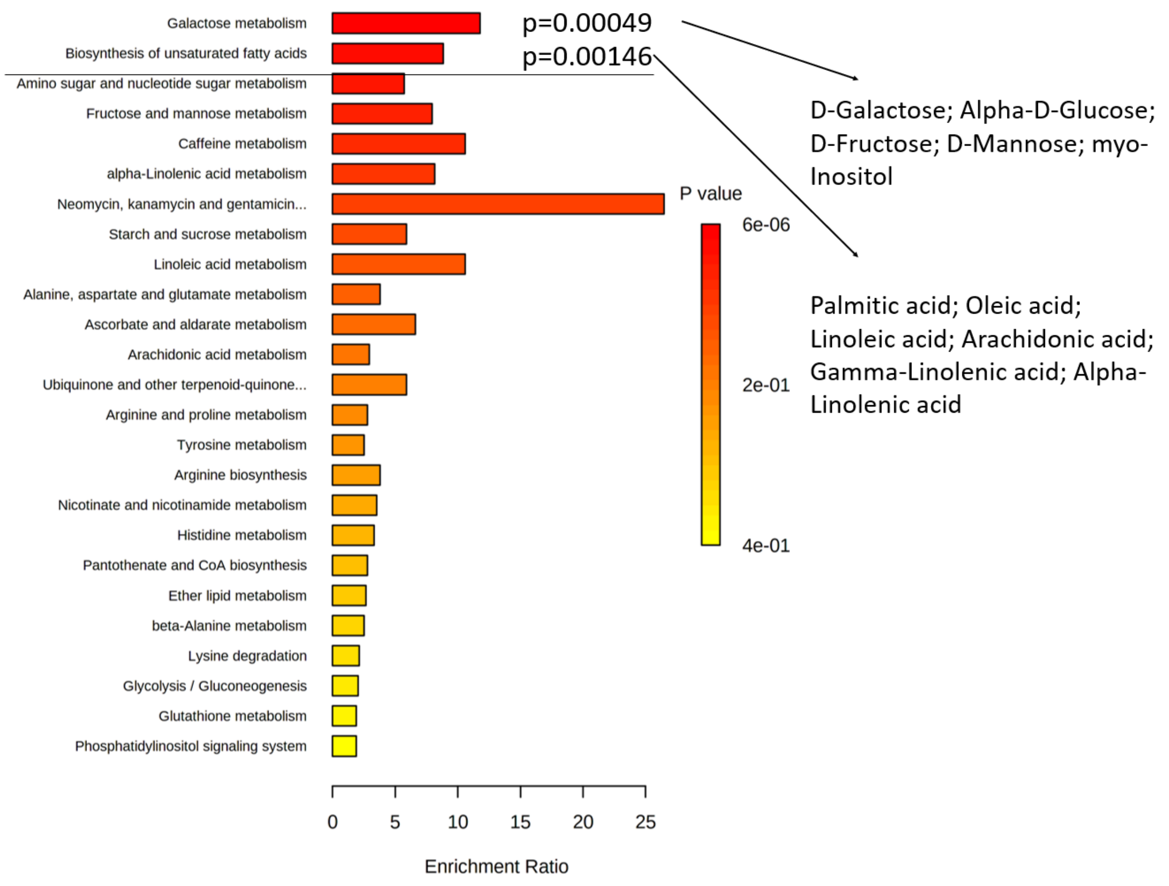


**Figure S15. Enrichment of pathways in the liver of 3% DSS treated mice compared to controls.** Enrichment pathway analysis using KEGG as reference found molecules involved in 25 different pathways. Galactose metabolism and biosynthesis of unsaturated fatty acids were significantly enriched, and the molecules involved in the pathway that are present in the dataset are listed.

**Supplementary Figure 16**

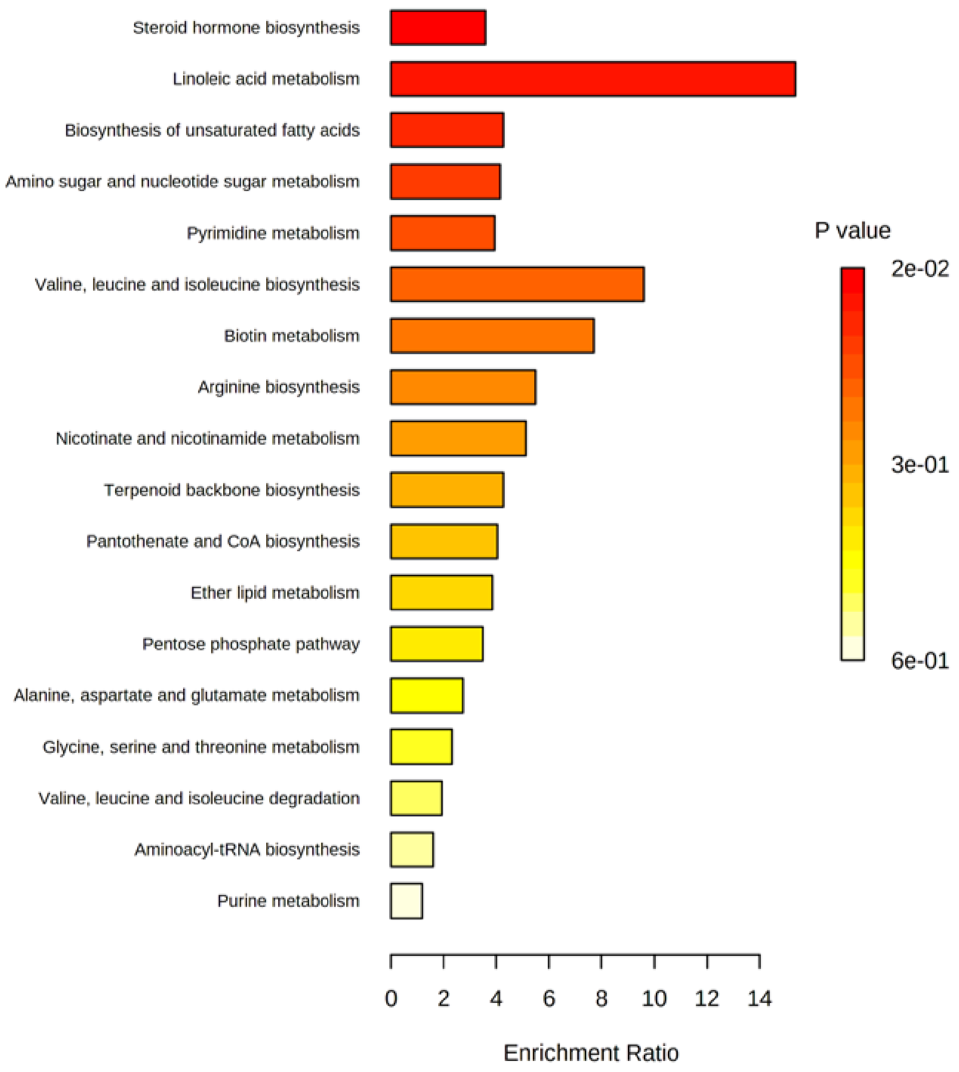


**Figure S16. Pathways altered in spleen of 3% DSS treated mice compared to controls.** Enrichment pathway analysis performed using KEGG as reference and found molecules involved in 18 different pathways such as steroid hormone biosynthesis, linoleic acid metabolism, biosynthesis of unsaturated fatty acids, amino sugar and nucleotide sugar metabolism, and pyrimidine metabolism. However, none of the pathways identified were significantly enriched.

**Supplementary Figure 17**

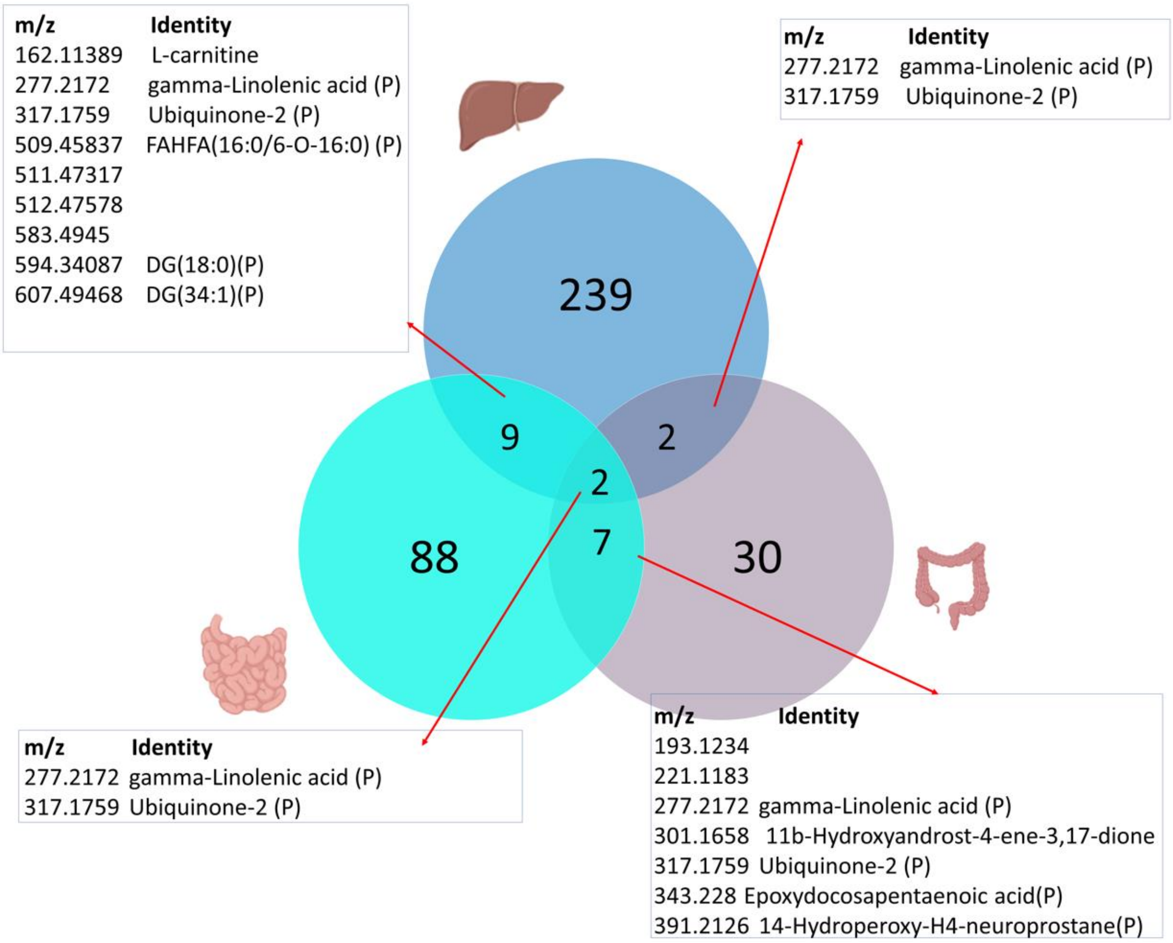


**Figure S17. Venn diagram showing molecules altered in the ileum, colon, and liver in 3% DSS treated mice compared to controls.** The top circle represents the 239 molecular changes in the liver, bottom-left circle represents 88 molecular changes in ileum and bottom-right represents 30 molecular changes in the colon. Overlapped segments indicate the number of molecules that are changed similarly between different tissue. Two metabolites were altered significantly in each of the ileum, colon and liver with ubiquinone-2 increased in the liver of DSS-treated mice in a manner similar to that seen in the intestine (2.73-fold, $p$=0.00108), while gamma-linolenic acid was similarly decreased in the liver (1.67-fold, $p$=0.0016) A further 7 metabolites were found to be common and similarly altered in both the liver and ileum. Carnitine (*m/z* 162.1138) was found to be significantly increased in both liver (3.73-fold, $p$=0.0034) and ileum (1.57-fold, $p$=0.0098) of DSS-treated mice. A further five metabolites were significantly altered but were all decreased in abundance in the ileum and liver of the DSS-treated mice. Fatty acyl esters of hydroxy fatty acid (FAHFA) (*m/z* 509.4583; ileum 3.91-fold, $p$=0.0019; liver 3.35-fold, $p$=0.00014) and a diglyceride (DG(34:1))(*m/z* 607.4946; ileum 3.18-fold, $p$=0.00233; liver 4.38-fold, $p$=0.00007) could be assigned putative identities but *m/z* 511.4731 (ileum 1.98-fold, $p$=0.001; liver 1.50-fold, $p$=0.000435), *m/z*

512.4757 (ileum 2.18-fold, *p*=0.0019; liver 1.53-fold, *p*=0.000405) and *m/z* 583.4945 (ileum 5.09-fold, *p*=0.001; liver 8.88-fold, *p*=0.0001) all remained unknowns. Lastly a diglyceride (DG(18:0)) (*m/z* 594.34087) was the only metabolite to show a significant decrease in the ileum (3.70-fold, *p*=0.00190) but a significant increase (2.17-fold, *p*= 0.00235) in the liver after DSS treatment. The *m/z* and putative identities of the molecules commonly shared between tissue are highlighted in boxes.

**Supplementary Figure 18**

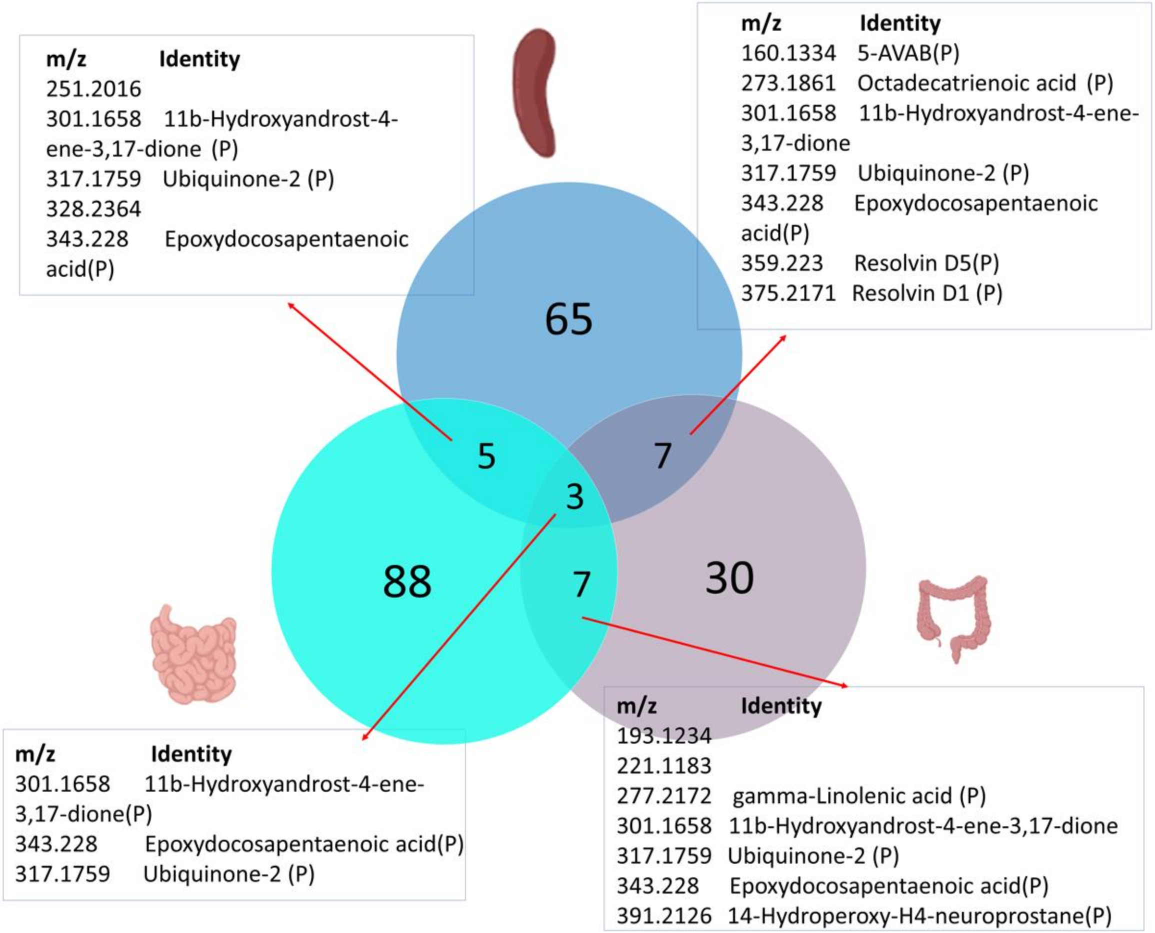


**Figure S18. Venn diagram of molecules altered in the ileum, colon, and spleen in 3% DSS treated mice compared to controls.** The top circle represents the 65 molecular changes in the liver, bottom-left circle represents 88 molecular changes in ileum and bottom-right represents 30 molecular changes in the colon. Overlapped segments indicate the number of molecules that are changed similarly between different tissue. In the spleen and colon 7 metabolites were found to be altered at both sites during DSS-induced colitis. *m/z* 160.1334 was putatively identified as 5-AVAB. This *m/z* was significantly decreased in the spleen (1.63-fold, *p*=0.028) and in the colon (1.81-fold, *p*=0.02) of DSS-treated mice. Six other metabolites were significantly increased in both the spleen and colon of 3% DSS treated mice. These metabolites were putatively identified as octadecatriencoic acid (2.04-fold colon, *p*=0.04; 1.54-fold spleen, *p*=0.0003), 11b-hydroxyandrost-4-ene-3,17-dione (15.8-fold colon, *p*=0.03; 4.65-fold spleen, *p*=0.04), ubiquinone-2 (7.9-fold colon, *p*=0.02; 3.38-fold spleen, *p*=0.005), epoxydocosapentaenoic acid (7.1-fold colon, *p*=0.02; 2.37-fold spleen, *p*=0.0016), resolvin D5 (19.0-fold colon, *p*=0.02; 2.28-fold spleen, *p*=0.0058) and resolvin D1 (19.14-fold colon, *p*=0.02; 2.26-fold spleen, *p*=0.0062)). Five metabolites were similarly altered in the ileum and spleen. Three of these metabolites, described above, are 11b-hydroxyandrost-4-ene-3-17-dione, ubiquinone-2 and epoxydocosapentaenoic acid and these were increased in the spleen 4.8-fold (*p*=0.0019), 3.06-fold (*p*=0.0012) and 3.4-fold (*p*=0.0038), respectively. The remaining two upregulated metabolites, which could not be assigned identities, have *m/z* 251.2016 (2.13-fold ileum, *p*=0.0013; 1.58-fold spleen, *p*=0.0091) and *m/z* 328.2364 (2.66-fold ileum, *p*=0.0012; 2.62-fold, *p*=0.0391).

The *m/z* and putative identities of the molecules commonly shared between tissue are highlighted in boxes.

**Supplementary Figure 19**

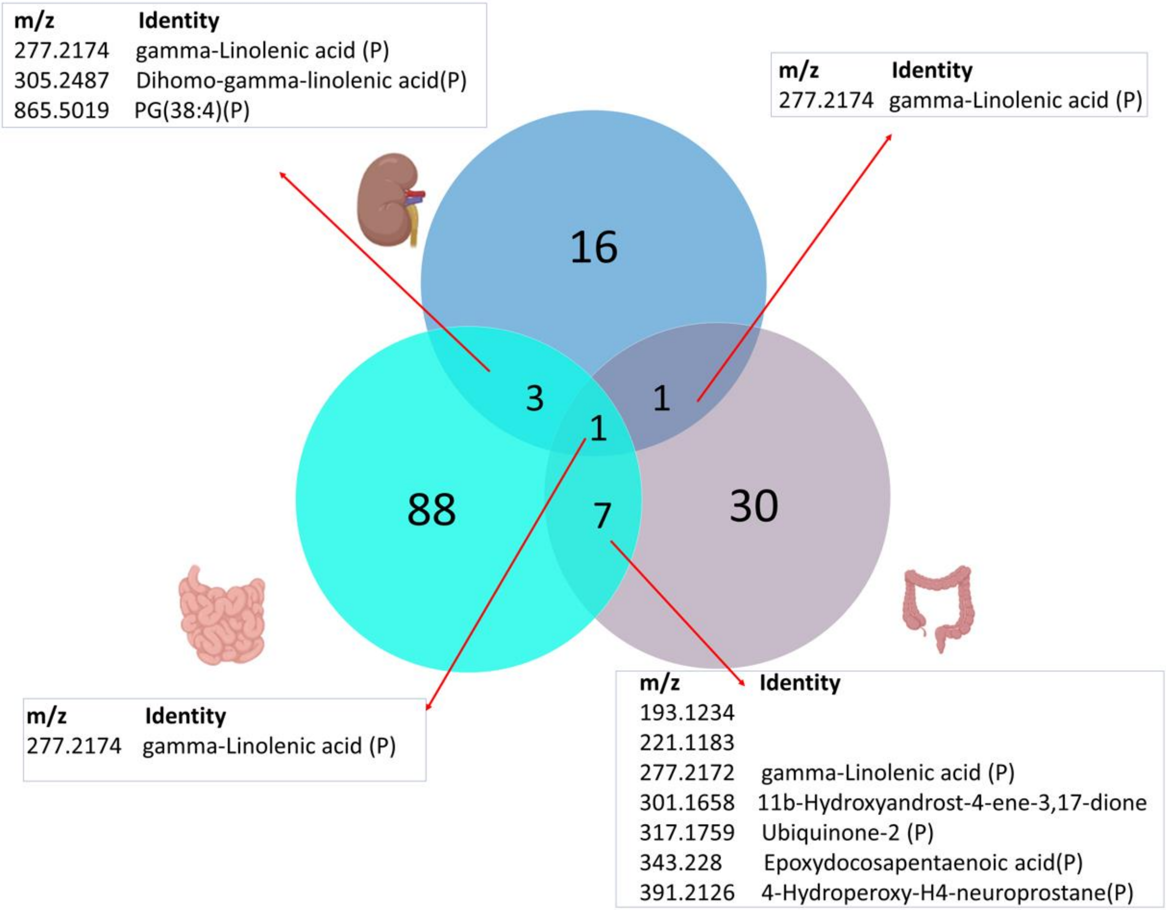


**Figure S19. Venn diagram of molecules altered in the ileum, colon, and kidney in 3% DSS treated mice compared to controls.** The top circle represents the 16 molecular changes in the kidney, bottom-left circle represents 88 molecular changes in ileum and bottom-right represents 30 molecular changes in the colon. Overlapped segments indicate the number of molecules that are changed similarly between different tissue. Gamma-linolenic acid was significantly changed in the ileum, colon, and liver in the colitis model. Again gamma-linolenic acid was also found to be significantly decreased 2.03-fold ($p$=0.016) in the kidney of the 3% DSS treated group compared to the controls. The molecule with *m/z* 865.5019 was putatively identified as a phosphatidylglyceride (PG(38:4)) and was found to be upregulated in the ileum (4.52-fold, $p$=0.0028) and kidney (14.5-fold, $p$=0.0259) during DSS colitis compared to the control. Lastly, the molecule with *m/z* 305.2481 was putatively identified as dihomo-gamma-linolenic acid. This molecule was significantly increased (1.96-fold, $p$=0.0019) in the ileum and decreased in the kidney (1.71-fold, $p$=0.0156) in the DSS-treated group compared to the controls. The *m/z* and putative identities of the molecules commonly shared between tissue are highlighted in boxes.

580 **Supplementary Figure 20**

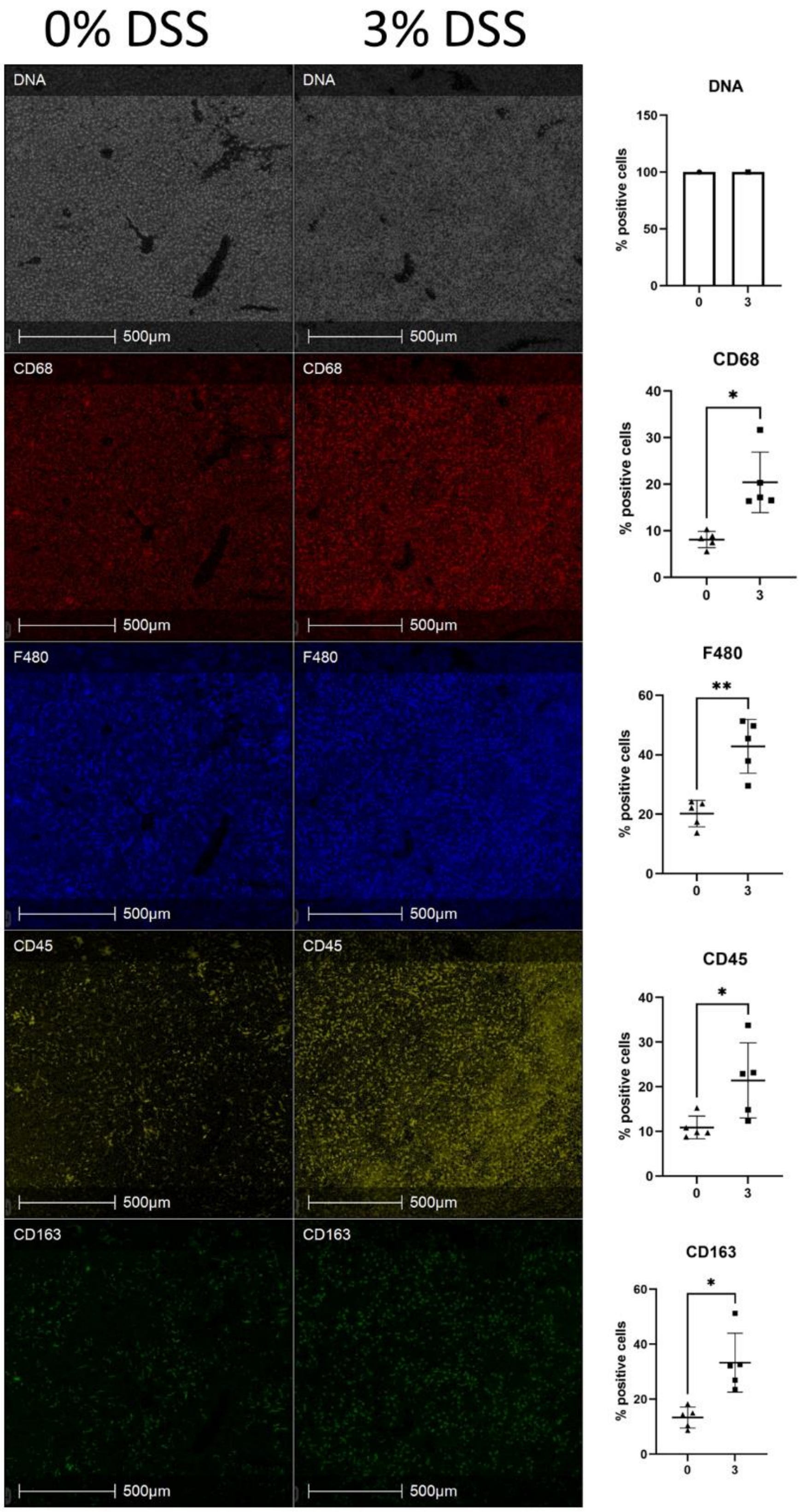


581

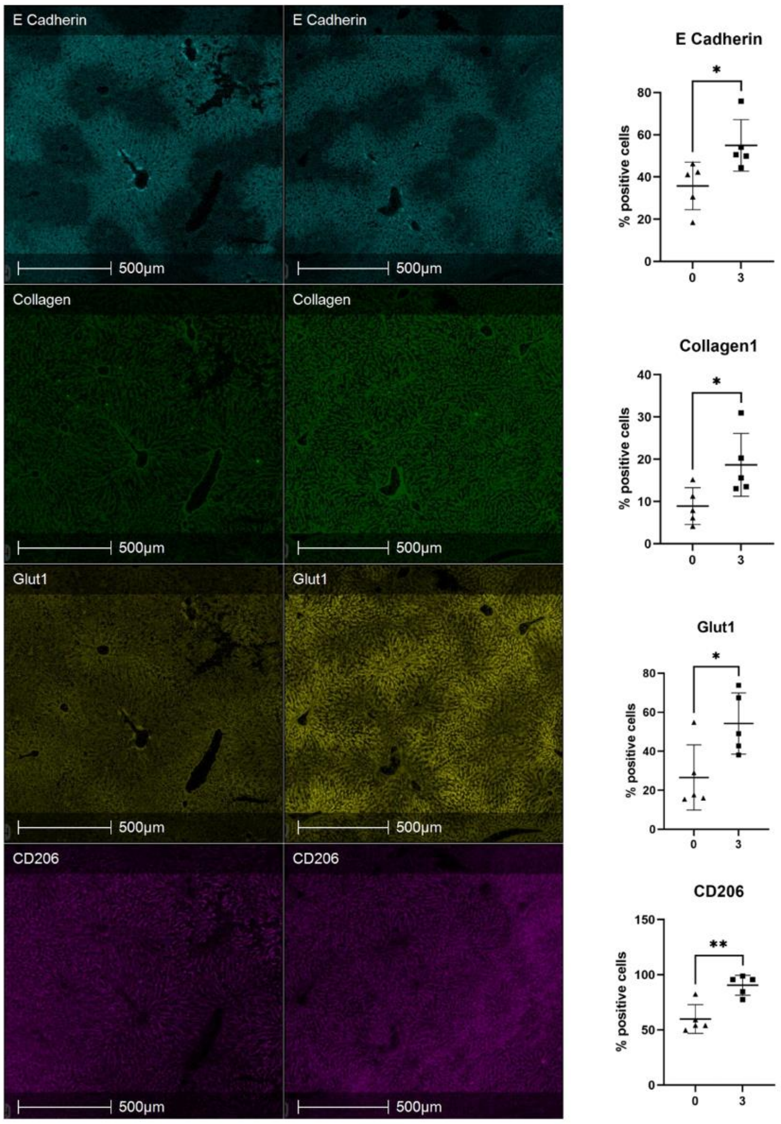


**Figure S20. Representative IMC images of biological markers of cell function in the liver of 3% DSS treated and control untreated mice.** From top to bottom, DNA intercalator identified single cells within the tissue section, markers for cell proliferation and immune cell function CD68, F480, CD45, CD163, E cadherin, collagen1, Glut1 and CD206. Percentage positive cells are represented by bar graphs showing five biological replicates. A t-test was performed to compare the two groups and *$p$<0.05, **$p$<0.01, ***$p$<0.001, ****$p$<0.0001 were considered statistically significant. The percentage of cells expressing markers $CD68^{+}$ (2.50-fold, $p$=0.0114), $CD45^{+}$ (1.96-fold, $p$=0.0465), $F480^{+}$ (2.11-fold, $p$=0.0026), $CD163^{+}$ (2.49-fold, $p$=0.0113), E $cadherin^{+}$ (1.53-fold, $p$=0.00325), $collagen^{+}$ (2.09-fold, $p$=0.0422), $Glut1^{+}$ (2.03-fold, $p$=0.0272) and $CD206^{+}$ (1.50-fold, $p$=0.0033) all showed an increase.

**Supplementary Tables**

**Supplementary Table 1**

| Marker | Tag | Cell type | Clone | Tag2 | Species | Dilution |
|---|---|---|---|---|---|---|
| ATPase | 141 | Membrane | EP1845Y | 141Pr | Rabbit | V; 1:100 |
| Cleaved caspase 3 | 142 | Apoptosis | D3E9 | 142Nd | Rabbit | V; 1:25 |
| Vimentin | 143 | Mesenchymal cells | D21H3 | 143Nd | Rabbit | V; 1:400 |
| B220 | 144 | B cells | RA3-6B2 | 144Nd | Rat | V; 1:25 |
| CD68 | 145 | Macrophages | FA-11 | 145Nd | Rat | V; 1:100 |
| CD31 | 146 | | | 146Nd | | V; 1:50 |
| CD45 | 147 | Myeloid cells | 30-F11 | 147Sm | Rat | V; 1:100 |
| PanCK | 148 | Epithelial cells | C11 | 148Nd | Mouse | V; 1:200 |
| CD19 | 149 | B cells | 6D5 | 149Sm | Rat | V; 1:50 |
| CD103 | 150 | Dendritic cells | AF1990 | 150Nd | Goat | V; 1:25 |
| Ly6G | 151 | | 1A8 | 151Eu | Rat | V; 1:50 |
| PAKT | 152 | | D9E | 152Sm | | V; 1:25 |
| CD11c | 153 | Dendritic cells | D1V9Y | 153Sm | Rabbit | V; 1:50 |
| CD11b | 154 | Macrophage subset | M1/70 | 154Sm | Rat | V; 1:50 |
| F4/80 | 155 | Macrophages | CI:A3-1 | 155Gd | Rat | V; 1:50 |
| CD163 | 156 | M2 Macrophages | TNKUPJ | 156Gd | Rat | V; 1:50 |
| e cadherin | 158 | Epithelial cells | 2.40E+11 | 158Gd | Rabbit | V; 1:100 |
| Collagen 1 | 159 | | Poly | 159 Tb | | V; 1:200 |
| glut1 | 160 | Hypoxia | EPR3915 | 160Gd | Rabbit | V; 1:100 |
| CD69 | 161 | | Poly | 161Dy | | V; 1:400 |
| Ki67 | 162 | Proliferation | B56 | 168Er | Mouse | V; 1:100 |
| a-SMA | 163 | Fibroblast and pericyte phenotype | Polyclonal | 163Dy | Rabbit | V; 1:400 |
| lyve1 | 164 | lymphatic vessels | poly | 164Dy | Rabbit | V; 1:300 |
| FOXP3 | 165 | Treg | FJK-16s | 165Ho | Rat | V; 1:25 |
| epcam | 166 | Epithelial cells | G8.8 | 166Er | Rat | V; 1:900 |
| NKp46 | 167 | NK cells | Polyclonal | 147Sm | Goat | V; 1:25 |
| CD8 | 168 | T cells (CD8+) | 53-6.7 | 146Nd | Rat | V; 1:50 |
| CD206 | 169 | M2 Macrophages | CD68C2 | 169Tm | Rat | V; 1:50 |
| arg1 | 170 | M2 Macrophages | Poly | 170Er | Sheep | V; 1:100 |

| | | | | | | |
|---|---|---|---|---|---|---|
| CD4 | 172 | T cells (CD4+) | RM4-5 | 172Yb | Rat | V; 1:50 |
| MHCII | 174 | Immune cells | M5/114.15.2 | 174Yb | Rat | V; 1:200 |
| Granzyme B | 176 | Activated T cells | Polyclonal | 176Yb | Goat | V; 1:300 |
| Collagen IV | 209 | ECM | Polyclonal | 209Bi | Rabbit | V; 1:100 |

**Supplementary Table 1.** List of immune cell marker antibodies, clone, tags and dilutions used for all IMC staining.

**Supplementary Table 2**

| Phenotype | Markers |
|---|---|
| Helper T cells | CD3 + CD4 |
| Cytotoxic T cells | CD3 + CD8 |
| Regulatory T cells (Tregs) | CD3 + CD4 + FOXP3 |
| Activated Cytotoxic T cells | CD3 + CD8 + Granzyme B |
| Natural killer cell | NKp46 |
| Activated natural killer cell | NKp46 + Granzyme B |
| B cells | B220 + CD19 |
| Neutrophils | Ly6G + CD11b |
| Dendritic cells | CD11c + M1 MHCII |
| Antigen-presenting Dendritic cells | CD11c + MHCII + CD103 |
| Macrophages | F4/80 + CD68 + CD11b (some overlap with dendritic cells) |
| M2c Macrophages | F4/80 + CD163 |
| M2a/c Macrophages | F4/80 + Arg1 |
| M2a Macrophages | F4/80 + CD206 |
| M1 and M2b Macrophages | F4/80 |
| M1 Macrophages | F4/80 + MHCII |
| Blood vessel | CD31 |
| Pericyte (found around large blood vessels) | aSMA |
| Mesenchymal cell | Vimentin |
| Epithelial cell | E Cadherin + PanCK + EpCam |
| Proliferation | Ki67 |
| PI3K signalling | pAKT |
| Hypoxia | GLUT1 |
| Apoptosis | CC3 |

**Supplementary Table 2.** Immune cell phenotypes defined by specific markers in IMC analysis.

### Supplementary Table 3

| m/z | Heatmap ID | compound_name | adduct | ppm | Mean 0% | Mean 3% | p-value |
|---|---|---|---|---|---|---|---|
| 131.035 | Glutaric acid(P) | Glutaric acid | M-H | 10 | 18.51141 | 39.78651 | 0.001903 |
| 132.0771 | Creatine | Creatine | M+H | 6 | 26.6331 | 61.16262 | 0.005447 |
| 162.1126 | L-carnitine | L-carnitine | M+H | 6 | 77.24398 | 121.6827 | 0.009899 |
| 167.1078 | m/z 167.1078 | | | | 22.19504 | 45.44605 | 0.001747 |
| 193.1234 | m/z 193.1234 | | | | 15.4677 | 27.59501 | 0.001903 |
| 195.1027 | m/z 195.1027 | | | | 29.71642 | 67.00769 | 0.001903 |
| 211.0975 | m/z 211.0975 | | | | 18.15224 | 34.83791 | 0.001903 |
| 221.1183 | m/z 221.1183 | | | | 12.98418 | 29.31121 | 0.001903 |
| 235.1339 | m/z 235.1339 | | | | 52.56577 | 115.0212 | 0.001845 |
| 251.2016 | m/z 251.2016 | | | | 43.0718 | 92.0656 | 0.001314 |
| 253.2174 | m/z 253.2174 | | | | 1918.392 | 656.5393 | 0.001903 |
| 254.2207 | m/z 254.2207 | | | | 489.9919 | 167.8958 | 0.001903 |
| 259.1518 | m/z 259.1518 | | | | 67.3124 | 156.7913 | 0.009899 |
| 277.2173 | GLA (P) | gamma-Linolenic acid | M-H | 0 | 858.6101 | 406.1114 | 0.001227 |
| 278.2207 | m/z 278.2207 | | | | 108.4817 | 50.61569 | 0.001227 |
| 285.1249 | m/z 285.1249 | | | | 9.299744 | 24.01997 | 0.001903 |
| 301.1658 | Hydroxy steroid(P) | 11b-Hydroxyandrost-4-ene-3,17-dione | M-H | 5 | 3.26949 | 15.59533 | 0.001903 |
| 303.233 | Arachidonic acid | Arachidonic acid | M-H | 4 | 2488.715 | 4830.248 | 0.001903 |
| 304.2364 | m/z 304.2364 | | | | 799.7057 | 1556.888 | 0.001903 |
| 305.2485 | m/z 305.2485 | | | | 56.22055 | 110.3871 | 0.001903 |
| 317.1759 | Ubiquinone-2(P) | Ubiquinone-2 | M-H | 5 | 16.0724 | 49.1914 | 0.001227 |
| 317.2126 | Leukotriene A4 | Leukotriene A4 | M-H | 7 | 48.1168 | 121.7857 | 0.002981 |
| 319.2281 | 5,6-EET(P) | 5,6-Epoxy-8,11,14-eicosatrienoic acid | M-H | 2 | 141.8269 | 336.1047 | 0.003894 |
| 327.233 | DHA | Docosahexaenoic acid | M-H | 5 | 814.25 | 2169.011 | 0.001974 |
| 328.2364 | m/z 328.2364 | | | | 128.3666 | 342.4678 | 0.001943 |
| 343.2201 | EDPA | Epoxydocosapentaenoic acid | M-H | 4 | 19.77427 | 67.14966 | 0.003894 |
| 349.1975 | 12-Oxo-LTB4(P) | 12-Oxo-20-hydroxy-leukotriene B4 | M-H | 8 | 22.06216 | 63.45958 | 0.002909 |
| 351.2173 | Lipoxin A4 | Lipoxin A4 | M-H | 1 | 55.84294 | 116.4331 | 0.003457 |
| 357.2406 | m/z 357.2406 | | | | 30.05336 | 61.45339 | 0.009899 |
| 363.2105 | m/z 327.23471 | | | | 17.24567 | 57.95692 | 0.001747 |
| 367.2442 | PGG2(P) | Prostaglandin G2 | M-H | 1 | 15.26352 | 45.37761 | 0.001314 |
| 369.277 | m/z 369.277 | | | | 11.44372 | 41.41741 | 0.009899 |
| 391.2122 | 14-Hydro-neuro(P) | 14-Hydroperoxy-H4-neuroprostane | M-H | 5 | 6.858846 | 30.54859 | 0.001903 |
| 395.3138 | m/z 395.3138 | | | | 24.91365 | 64.43831 | 0.009899 |
| 395.3502 | m/z 395.3502 | | | | 13.67504 | 45.53829 | 0.005447 |
| 425.3394 | 20α-OHC(P) | 20alpha-hydroxy cholesterol | M+Na | 2 | 38.54771 | 69.84752 | 0.009899 |
| 437.3397 | m/z 437.3397 | | | | 10.34558 | 32.61636 | 0.009899 |
| 441.3336 | 20a,22b-DOHC(P) | 20a,22b-Dihydroxycholesterol | M+Na | 2 | 19.46659 | 41.79346 | 0.009933 |
| 463.3036 | m/z 463.3036 | | | | 43.04457 | 5.886998 | 0.009899 |
| 465.1743 | PGH2(P) | Prostaglandin H2 2-glyceryl Ester | M+K | 2 | 12.78091 | 38.20404 | 0.004235 |
| 465.3045 | CS(P) | Cholesterol sulfate | M-H | 1 | 33.2425 | 4.233776 | 0.000998 |
| 465.3192 | m/z 465.3192 | | | | 96.64857 | 218.7687 | 0.004235 |
| 466.3078 | m/z 466.3078 | | | | 27.57423 | 63.99506 | 0.000998 |
| 508.3044 | m/z 508.3044 | | | | 10.09489 | 29.71496 | 0.003614 |

| | | | | | | | |
|---|---|---|---|---|---|---|---|
| 509.4576 | FAHFA(16:0)(P) | FAHFA(16:0/6-O-16:0) | M-H | 2 | 53.5792 | 13.78694 | 0.001903 |
| 510.4609 | m/z 510.4609 | | | | 26.14738 | 5.056658 | 0.001903 |
| 511.4732 | m/z 511.4732 | | | | 175.7986 | 88.69382 | 0.001903 |
| 512.4764 | m/z 512.4764 | | | | 40.79809 | 18.6329 | 0.001903 |
| 546.354 | LycoPC(20:3) | LysoPC(20:3(8Z,11Z,14Z)/0:0) | M+H | 3 | 55.5838 | 102.0115 | 0.004235 |
| 557.4561 | m/z 557.4561 | | | | 49.9586 | 19.31112 | 0.001227 |
| 567.5356 | m/z 567.5356 | | | | 22.93004 | 64.37812 | 0.001277 |
| 568.5387 | m/z 568.5387 | | | | 3.96036 | 14.91338 | 0.001903 |
| 583.4942 | m/z 583.4942 | | | | 45.28996 | 8.891721 | 0.001903 |
| 584.4978 | m/z 584.4978 | | | | 10.26839 | 1.554362 | 0.001943 |
| 587.5028 | m/z 587.5028 | | | | 20.31425 | 35.32787 | 0.014693 |
| 594.3413 | PC(20:1)(P) | PC(18:1(12Z)-2OH(9,10)/2:0) | M-H | 2 | 110.1503 | 29.7129 | 0.001903 |
| 605.4545 | DG(32:1)(P) | DG(14:0/18:1(9Z)/0:0) | M+K | 2 | 78.64177 | 25.3103 | 0.010204 |
| 607.4734 | m/z 607.4734 | | | | 8.390444 | 31.04291 | 0.002871 |
| 607.4943 | DG(34:1) (P) | DG(16:0/18:1(12Z)-O(9S,10R)/0:0) | M-H | 1 | 43.57882 | 13.68875 | 0.002337 |
| 629.455 | DG(16:0) (P) | DG(14:0/20:3(8Z,11Z,14Z)/0:0) | M+K | 2 | 209.8574 | 38.52766 | 0.009933 |
| 631.4899 | DG(36:4) (P) | DG(20:4(5Z,7E,11Z,14Z)-OH(9)/0:0/i-16:0) | M-H | 9 | 36.97692 | 7.677196 | 0.003829 |
| 701.5625 | PA(36:2) (P) | PA(20:2(11Z,14Z)/16:0) | M+H | 2 | 82.98722 | 9.306941 | 0.009933 |
| 702.5005 | PE-NMe(32:2) (P) | PE-NMe(18:2(9Z,12Z)/14:0) | M+H | 14 | 36.02354 | 3.804072 | 0.009933 |
| 703.5279 | PA(36:1) (P) | PA(22:1(13Z)/14:0) | M+H | 0 | 68.36887 | 9.47622 | 0.009899 |
| 704.5315 | PC(30:1) (P) | PC(16:1(9Z)/14:0) | M+H | 14 | 20.69057 | 2.720816 | 0.009899 |
| 841.6671 | PA (46:2) (P) | PA(24:1(15Z)/22:1(13Z)) | M+H | 1 | 54.33888 | 13.32923 | 0.009899 |
| 843.6834 | PA (46:1) (P) | PA(24:1(15Z)/22:0) | M+H | 1 | 73.93822 | 20.10841 | 0.009899 |
| 847.7393 | m/z 847.7393 | | | | 23.56876 | 3.401429 | 0.003831 |
| 854.73 | m/z 854.73 | | | | 128.6605 | 21.60667 | 0.009899 |
| 855.7419 | TG(32:2) P) | TG(15:0/20:1(11Z)/15:0) | M+Na | 0 | 143.3241 | 28.17171 | 0.009899 |
| 856.7457 | m/z 856.7457 | | | | 73.97373 | 14.58817 | 0.009899 |
| 865.5019 | PG(38:4)(P) | PG(18:0/20:4(6Z,8E,10E,14Z)-2OH(5S,12R)) | M+Cl | 2 | 7.877709 | 35.65681 | 0.002871 |
| 867.6741 | PA(46:0)(P) | PA(22:0/24:0) | M+Na | 0 | 111.6849 | 24.31171 | 0.009899 |
| 868.6808 | PE-NMe(44:3)(P) | PE-NMe(24:1(15Z)/20:2(11Z,14Z)) | M+H | 7 | 65.14419 | 14.15516 | 0.009899 |
| 869.6996 | PA(44:2)(P) | PA(24:1(15Z)/24:1(15Z)) | M+H | 2 | 226.1649 | 56.43565 | 0.009899 |
| 870.7031 | PC(46:2)(P) | PC(24:1(15Z)/18:1(11Z)) | M+H | 7 | 126.6299 | 32.11966 | 0.009899 |
| 871.7151 | PA(48:1)(P) | PA(24:1(15Z)/24:0) | M+H | 2 | 136.7504 | 39.93581 | 0.009899 |
| 881.7571 | TG(54:5)(P) | TG(14:0/18:0/22:5(4Z,7Z,10Z,13Z,16Z)) | M+H | 2 | 198.736 | 41.89184 | 0.004733 |
| 882.7615 | m/z 882.7615 | | | | 105.5943 | 22.51055 | 0.004733 |
| 893.7012 | TG(52:4)(P) | TG(16:1(9Z)/14:1(9Z)/22:2(13Z,16Z)) | M+K | 1 | 124.2978 | 25.89853 | 0.004838 |
| 894.7032 | PC(44:4)(P) | PC(24:1(15Z)/20:3(5Z,8Z,11Z)) | M+H | 9 | 71.4225 | 14.86303 | 0.004975 |
| 895.7146 | TG(52:3)(P) | TG(16:0/22:2(13Z,16Z)/14:1(9Z)) | M+K | 0 | 197.9684 | 52.55442 | 0.004983 |
| 896.7183 | PC(44:3)(P) | PC(24:1(15Z)/20:2(11Z,14Z)) | M+H | 9 | 105.9555 | 28.17985 | 0.005399 |
| 341.2457 | MA(P) | Methyl Arachidonate | M+Na | 1 | 9.578257 | 32.72954 | 0.005401 |
| 367.3187 | m/z 367.3187 | | | | 20.14158 | 54.79385 | 0.009899 |

**Supplementary Table 3.** The *m/z*, abbreviated names shown in heatmap (Fig. 1A), and full compound name for molecules found to be significantly changed in the ileum of DSS-treated mice. The table includes the mean relative abundance value for each molecule within the groups. *P*-values highlighted in grey are for molecules down regulated in the 3% DSS-treated group compared to the control group and *p*-values not highlighted show molecules upregulated in the treated group compared to the control.

**Supplementary Table 4**

| mz | VIP list |
|---|---|
| 162.1130066 | 1.306210041 |
| 466.3070068 | 1.305969954 |
| 465.3049927 | 1.3046 |
| 546.3540039 | 1.259809971 |
| 587.5029907 | 1.235599995 |
| 317.177002 | 1.191900015 |
| 367.2120056 | 1.190739989 |
| 567.5339966 | 1.17723 |
| 256.0589905 | 1.172199965 |
| 391.2109985 | 1.167940021 |
| 363.2099915 | 1.16779995 |
| 277.2170105 | 1.151690006 |
| 278.2219849 | 1.151550055 |
| 343.2290039 | 1.139610052 |
| 568.5390015 | 1.134230018 |
| 557.4559937 | 1.130779982 |
| 865.5020142 | 1.130169988 |
| 251.128006 | 1.126889944 |
| 465.2260132 | 1.11748004 |
| 317.2099915 | 1.115929961 |
| 193.1230011 | 1.114580035 |
| 344.2319946 | 1.111469984 |
| 320.2309875 | 1.110579967 |
| 319.2290039 | 1.110360026 |
| 301.1789856 | 1.110200047 |
| 235.1360016 | 1.105029941 |
| 349.1990051 | 1.104580045 |
| 385.2720032 | 1.104089975 |
| 221.1190033 | 1.099099994 |
| 167.1080017 | 1.099009991 |
| 351.2179871 | 1.095260024 |
| 594.34198 | 1.090180039 |
| 395.3510132 | 1.086809993 |
| 195.1000061 | 1.067849994 |
| 211.095993 | 1.06475997 |
| 372.276001 | 1.062000036 |
| 328.2369995 | 1.060909986 |
| 461.2309875 | 1.059649944 |
| 327.2319946 | 1.057909966 |
| 511.4729919 | 1.05637002 |
| 253.2169952 | 1.053130031 |
| 254.2169952 | 1.052770019 |
| 583.4940186 | 1.052250028 |
| 305.2409973 | 1.051090002 |
| 132.076004 | 1.050529957 |
| 512.4780273 | 1.049790025 |
| 285.1289978 | 1.049319983 |

**Supplementary Table 4.** PLS-DA generated VIP>1 list of molecules in the ileum that contribute towards group separation in the DSS colitis model.

**Supplementary Table 5**

| *m/z* | Heatmap ID | compound_name | adduct | ppm | mean 0% | mean 3% | p-value |
|---|---|---|---|---|---|---|---|
| 137.071 | 1-MNA(P) | 1-Methylnicotinamide | M+H | 12 | 33.89 | 88.04 | 0.001162 |
| 140.0682 | L-Valine(P) | L-Valine | M+Na | 7 | 165.8 | 405.4 | 0.011395 |
| 159.0664 | Pimelic acid(P) | Pimelic acid | M-H | 4 | 19.29 | 34.46 | 0.03815 |
| 160.133 | 5-AVAB(P) | 5-amino valeric acid betaine | M+H | 6 | 741 | 406.4 | 0.02202 |
| 167.018 | Uric acid(P) | Uric acid | M-H | 12 | 28.57 | 93.51 | 0.021785 |
| 193.1234 | m/z 193.1227 | | | | 9.994 | 23.22 | 0.023974 |
| 221.1183 | m/z 221.1183 | | | | 10.11 | 27.89 | 0.03815 |
| 235.1339 | m/z 235.1339 | | | | 43.17 | 87.87 | 0.040343 |
| 261.1493 | m/z 261.1493 | | | | 6.989 | 27.22 | 0.023974 |
| 263.1652 | m/z 263.1652 | | | | 7.707 | 25.11 | 0.023974 |
| 273.1861 | ODT acid | (10Z,14E,16E)-10,14,16-Octadecatrien-12-ynoic acid | M-H | 6 | 17.56 | 35.71 | 0.040343 |
| 277.2172 | GLA | gamma-Linolenic acid | M-H | 3 | 415.5 | 202.1 | 0.045769 |
| 278.2207 | m/z 278.2207 | | | | 78.49 | 37.38 | 0.045769 |
| 300.0397 | N-AGLANS | N-Acetylglucosamine 6-sulfate | M-H | 3 | 0.09121 | 11.7 | 0.006095 |
| 301.1808 | Hydroxy steroid (P) | 11b-Hydroxyandrost-4-ene-3,17-dione | M-H | 6 | 1.192 | 18.85 | 0.037431 |
| 317.1759 | Ubiquinone-2(P) | Ubiquinone-2 | M-H | 2 | 7.63 | 60.24 | 0.02292 |
| 319.228 | 5-HETE(P) | 5-HETE | M-H | 2 | 145 | 296 | 0.040343 |
| 320.2312 | 19(S)-HETE(P) | 19-Hydroxyeicosatetraenoic acid | M-H | 2 | 29.83 | 62.71 | 0.040343 |
| 331.2643 | EA(P) | Ethyl Arachidonate | M-H | 4 | 125.4 | 552.9 | 0.043938 |
| 332.2676 | m/z 332.2676 | | | | 29.2 | 131 | 0.043938 |
| 333.2709 | m/z 333.2709 | | | | 1.288 | 13.13 | 0.043938 |
| 343.228 | EDPA (P) | Epoxydocosapentaenoic acid | M-H | 1 | 8.95 | 62.92 | 0.021785 |
| 344.2312 | m/z 344.2312 | | | | 1.016 | 26.92 | 0.021785 |
| 345.2436 | DHT propionate(P) | Dihydrotestosterone propionate | M-H | 2 | 5.341 | 31.91 | 0.023974 |
| 347.2593 | CTXA2(P) | Carbocyclic thromboxane A2 | M-H | 0 | 4.247 | 31.24 | 0.021785 |
| 359.223 | Resolvin D5(P) | Resolvin D5 | M-H | 2 | 1.069 | 20.29 | 0.021785 |
| 359.2957 | m/z 359.2957 | | | | 5.139 | 41.88 | 0.03815 |
| 375.2177 | Resolvin D1(P) | Resolvin D1 | M-H | 1 | 0.6427 | 12.25 | 0.023974 |
| 391.2126 | 14-Hydro-neuro(P) | 14-Hydroperoxy-H4-neuroprostane | M-H | 5 | 1.981 | 25.04 | 0.023974 |

**Supplementary Table 5.** The *m/z*, abbreviated names shown in heatmap (Fig. 1B), and full compound name for molecules found to be significantly changed in the colon of DSS-treated mice. Table includes the mean relative abundance value for each molecule within the groups. *P*-values highlighted in grey are for molecules down regulated in the treated group compared to control group and *p*-values not highlighted show molecules upregulated in the treated group compared to the control.

**Supplementary Table 6**

| mz | VIP list |
|---|---|
| 140.069 | 1.66642 |
| 137.070007 | 1.60515 |
| 160.132004 | 1.48317 |
| 300.039001 | 1.2944 |
| 167.018997 | 1.22168 |
| 188.985001 | 1.03454 |
| 278.218994 | 1.00477 |
| 277.21701 | 1.00427 |

**Supplementary Table 6.** PLS-DA generated VIP>1 list of molecules in the colon that contribute towards group separation in DSS treated mice compared to controls.

### Supplementary Table 7

| *m/z* | Heatmap ID | Compound name | ppm | Mean 0% | Mean 3% | p-value |
|---|---|---|---|---|---|---|
| 128.03638 | pyroglut acid(P) | Pyroglutamic acid | 8 | 31.21 | 53.63 | 0.00076 |
| 132.03081 | | | | 15.34 | 36.96 | 0.002047 |
| 151.02595 | L-Aspartic acid(P) | L-Aspartic acid | 4 | 253 | 407.7 | 0.001693 |
| 161.04562 | | | | 59.78 | 39.12 | 0.000995 |
| 178.01636 | | | | 14.14 | 39.73 | 0.002318 |
| 179.05575 | | | | 278.6 | 164.5 | 0.000313 |
| 180.05995 | D-Galactose(P) | D-Galactose | 2 | 17.13 | 6.792 | 0.000039 |
| 253.2166 | | | | 4031 | 916.5 | 0.00003 |
| 253.34612 | Palmitelaidic acid(P) | Palmitelaidic acid | 3 | | 0.04633 | 0.001032 |
| 254.22081 | | | | 683.6 | 155.6 | 0.00003 |
| 254.55981 | | | | 11.25 | 4.667 | 0.001724 |
| 265.21749 | | | | 19.59 | 7.815 | 0.000464 |
| 267.19552 | HDD acid(P) | 10Z-Heptadecenoic acid | 4 | 131.5 | 33.13 | 0.000712 |
| 267.2339 | 9Z-HDD acid(P) | 9Z-Heptadecenoic acid | 4 | 143.7 | 67.7 | 0.000058 |
| 268.19972 | | | | 19.99 | 1.228 | 0.000442 |
| 268.2365 | | | | 25.02 | 7.803 | 0.000111 |
| 275.20038 | Stearidonic acid(P) | Stearidonic acid | 5 | 89.57 | 37.59 | 0.000173 |
| 276.20458 | | | | 16.16 | 2.995 | 0.00003 |
| 277.21838 | GLA(P) | gamma-Linolenic acid | 4 | 1026 | 631.6 | 0.00162 |
| 278.21939 | | | | 194.8 | 119.8 | 0.001713 |
| 279.23319 | Linoelaidic acid(P) | Linoelaidic acid | 1 | 8050 | 11345 | 0.000435 |
| 280.23739 | | | | 1547 | 2188 | 0.000435 |
| 280.46925 | petroselinate(P) | petroselinate | 12 | 13.79 | 2.159 | 0.000176 |
| 281.09928 | | | | 20.67 | 12.52 | 0.000195 |
| 281.24799 | | | | 17096 | 10374 | 0.000071 |
| 281.3999 | | | | 19.81 | 10.28 | 0.000195 |
| 282.2522 | | | | 2205 | 1335 | 0.000071 |
| 283.2564 | | | | 304.5 | 173.8 | 0.000132 |
| 284.25901 | | | | 18.27 | 4.106 | 0.000195 |
| 300.04006 | | | | 0.1048 | 15.65 | 0.00003 |
| 309.27938 | | | | 295.1 | 140 | 0.000071 |
| 310.28359 | | | | 62.7 | 28.72 | 0.000084 |
| 315.20867 | Linoleic acid(P) | Linoleic acid | 3 | 31.41 | 83.05 | 0.000208 |
| 316.21287 | 3-hydroxycarnitine(P) | 3-hydroxynonanoyl carnitine | 0 | 3.112 | 14.41 | 0.000097 |

| | | | | | | |
|---|---|---|---|---|---|---|
| 317.1755 | Ubiquinone-2(P) | Ubiquinone-2 | 1 | 33.33 | 91.29 | 0.001089 |
| 318.17971 | | | | 3.414 | 16.07 | 0.001089 |
| 325.2747 | | | | 18.18 | 3.543 | 0.00156 |
| 337.31077 | | | | 33.79 | 8.324 | 0.000852 |
| 339.21205 | AA | Arachidonic acid | 7 | 15.55 | 47.46 | 0.000422 |
| 351.18415 | | | | 12.93 | 37.57 | 0.000843 |
| 363.23141 | MG(16:1)(P) | MG(16:1(9Z)/0:0/0:0) | 2 | 112.6 | 17.71 | 0.000011 |
| 364.23402 | | | | 34.82 | 1.388 | 0.000011 |
| 391.21224 | 14-Hydro-neuro(P) | 14-Hydroperoxy-H4-neuroprostane | 3 | 34.17 | 83.25 | 0.001891 |
| 393.26322 | | | | 222.2 | 164.9 | 0.002594 |
| 403.18854 | PGG2(P) | Prostaglandin G2 | 2 | 1.523 | 15.28 | 0.001926 |
| 417.28578 | Palmitoyl(P) | Palmitoyl glucuronide | 0 | 11.07 | 0.08066 | 0.000588 |
| 433.28111 | | | | 112.2 | 8.95 | 0.000014 |
| 434.28371 | | | | 16.31 | 0.1557 | 0.000045 |
| 435.29591 | | | | 405.8 | 190.4 | 0.000058 |
| 436.30012 | | | | 97.99 | 44.33 | 0.000051 |
| 441.27797 | | | | 0.9505 | 16.74 | 0.000309 |
| 457.28128 | | | | 30.06 | 12.02 | 0.001425 |
| 461.3125 | | | | 528.9 | 189.9 | 0.000019 |
| 462.3151 | | | | 93.72 | 32.94 | 0.000019 |
| 477.22627 | PGG2-2-glyc(P) | Prostaglandin G2 2-glyceryl Ester | 0 | 11.62 | 33.7 | 0.001143 |
| 477.30622 | | | | 28.77 | 2.28 | 0.000073 |
| 480.30924 | | | | 25.87 | 53.13 | 0.002573 |
| 485.31267 | Vitamin K1(P) | Vitamin K1 | 13 | 19.33 | 3.964 | 0.00006 |
| 489.34389 | | | | 20.43 | 0.6295 | 0.000018 |
| 497.34874 | | | | 48.14 | 14.19 | 0.00102 |
| 507.44196 | | | | 43.06 | 9.682 | 0.000309 |
| 508.30225 | | | | 39.53 | 71.85 | 0.002243 |
| 508.44457 | | | | 7.65 | 0.6146 | 0.001089 |
| 509.45837 | FAHFA(32:0)(P) | FAHFA(16:0/10-O-16:0) | 2 | 151.6 | 45.06 | 0.000014 |
| 510.46097 | | | | 79.23 | 18.48 | 0.000014 |
| 511.47317 | | | | 252.6 | 167.8 | 0.000435 |
| 512.47578 | | | | 89.3 | 58.42 | 0.000405 |
| 525.31937 | | | | 12.81 | 2.507 | 0.000468 |
| 526.38274 | | | | 28.34 | 9.224 | 0.002635 |
| 531.44054 | | | | 19.31 | 3.284 | 0.000222 |
| 533.45535 | | | | 147.3 | 69.37 | 0.000048 |

| | | | | | | |
|---|---|---|---|---|---|---|
| 534.45955 | | | | 53.96 | 21.56 | 0.000071 |
| 537.48976 | FAHFA(34:0)(P) | FAHFA(16:0/6-O-18:0) | 2 | 373.3 | 207.1 | 0.00003 |
| 538.49236 | | | | 211.7 | 117.6 | 0.00003 |
| 545.36189 | | | | 17.04 | 1.903 | 0.000286 |
| 552.29698 | | | | 12.21 | 29.3 | 0.001455 |
| 558.33341 | LysoPC(18:0)(P) | LysoPC(0:0/18:0) | 0 | 7.585 | 35.18 | 0.000017 |
| 559.47193 | | | | 181.6 | 320 | 0.000442 |
| 560.47614 | | | | 70.89 | 125 | 0.000488 |
| 563.50474 | | | | 321.4 | 204.2 | 0.000309 |
| 564.50735 | Octadecadiene(P) | N-(2R-Hydroxyhexadecanoyl)-2S-amino-9-methyl-4E,8E-octadecadiene-1,3R-diol | 13 | 85.31 | 54.37 | 0.000309 |
| 572.48182 | Cer(d35:1)(P) | Cer(d18:1/16:0) | 1 | 11.5 | 124.7 | 0.000164 |
| 573.48443 | | | | 1.133 | 29.7 | 0.000248 |
| 574.47904 | Cer(d36:4)(P) | Cer(d16:1/20:3(6,8,11)-OH(5)) | 9 | 0.3994 | 20.14 | 0.000588 |
| 581.47809 | TG(32:0)(P) | TG(16:0/8:0/8:0) | 1 | 36.06 | 6.427 | 0.000731 |
| 582.4823 | | | | 10.71 | 0.6303 | 0.001891 |
| 583.47371 | | | | 32.9 | 106 | 0.000902 |
| 583.4945 | | | | 71.64 | 8.063 | 0.000019 |
| 584.47632 | Cer(d37:6)(P) | Cer(d17:1/20:5(6E,8Z,11Z,14Z,17Z)-OH(5)) | 13 | 7.452 | 28.05 | 0.001612 |
| 584.4971 | | | | 40.29 | 1.491 | 0.000018 |
| 591.53613 | | | | 14.2 | 1.305 | 0.000018 |
| 592.32607 | | | | 30.81 | 65.89 | 0.001891 |
| 594.34087 | | | | 63.66 | 138.6 | 0.002356 |
| 595.34508 | DG(30:6)(P) | DG(22:6(4Z,7Z,11E,13Z,15E,19Z)-2OH(10S,17)/0:0/8:0) | 7 | 18.37 | 42.62 | 0.00202 |
| 601.46145 | | | | 8.282 | 0.09036 | 0.000246 |
| 607.49468 | | | | 148.7 | 33.91 | 0.000071 |
| 608.49728 | Cer(d16:1/PGF1alpha)(P) | Cer(d16:1/PGF1alpha) | 13 | 30.26 | 88.49 | 0.007231 |
| 611.52589 | | | | 277.3 | 115 | 0.000031 |
| 612.52849 | | | | 169.3 | 69.76 | 0.000035 |
| 613.5359 | | | | 16.76 | 3.646 | 0.000062 |
| 625.46483 | DG(34:3)(P) | DG(20:3(8Z,11Z,14Z)/14:0/0:0) | 1 | 9.643 | 1.155 | 0.000953 |
| 626.53619 | | | | 3.029 | 36.16 | 0.000435 |
| 629.49284 | DG(34:1)(P) | DG(14:1(9Z)/20:0/0:0) | 2 | 39.4 | 9.589 | 0.000843 |
| 630.49545 | | | | 20.2 | 2.804 | 0.000132 |
| 631.49166 | DG(36:3)(P) | DG(i-16:0/20:3(8Z,11Z,14Z)-O(5,6)/0:0) | 4 | 33.35 | 11.46 | 0.001511 |
| 632.49586 | | | | 11.99 | 2.376 | 0.001877 |
| 637.54087 | DG(36:0)(P) | TG(18:0/10:0/8:0) | 1 | 315.7 | 147.9 | 0.00004 |
| 638.33081 | | | | 9.962 | 44.13 | 0.001454 |

| | | | | | | |
|---|---|---|---|---|---|---|
| 638.54508 | Cer(t36:1)(P) | Cer(d18:0/PGF1alpha) | 13 | 134.1 | 61.48 | 0.000045 |
| 639.48532 | DG(35:3)(P) | DG(15:0/20:3(5Z,8Z,11Z)/0:0) | 14 | 13.38 | 0.5345 | 0.000248 |
| 641.50013 | DG(35:2n6)(P) | DG(15:0/0:0/20:2n6) | 13 | 50.44 | 20.78 | 0.001425 |
| 642.50433 | Cer(d20:1/PGJ2)(P) | Cer(d20:1/PGJ2) | 9 | 17.77 | 3.468 | 0.001939 |
| 648.35047 | | | | 15.79 | 44.4 | 0.00202 |
| 655.50783 | DG(36:2)(P) | DG(14:0/22:2(13Z,16Z)/0:0) | 1 | 36.26 | 6.024 | 0.00006 |
| 656.51203 | Cer(t38:3)(P) | Cer(t18:0/20:3(6,8,11)-OH(5)) | 14 | 17.28 | 1.879 | 0.000022 |
| 657.4139 | DG(PGE2)(P) | DG(13:0/PGE2/0:0) | 0 | 3.86 | 25.56 | 0.00033 |
| 662.36776 | | | | 25.52 | 62.64 | 0.002591 |
| 682.59258 | Cer(d42:2)(P) | Cer(d18:1/24:1(15Z)) | 2 | 28.7 | 73.8 | 0.000878 |
| 683.59359 | | | | 16.94 | 43.56 | 0.000883 |
| 702.39684 | azahexane(P) | 1-(4-Biphenylyl)-4(S)-hydroxy-5(S)-2,5-bis((N-(methoxycarbonyl-)-L-tert-leucinyl)amino)-6-phenyl-2-azahexane | 14 | 8.762 | 22.31 | 0.000435 |
| 773.53389 | | | | 6.33 | 50.69 | 0.000111 |
| 790.50304 | | | | 13.91 | 59.56 | 0.000712 |
| 791.50564 | | | | 2.714 | 16.58 | 0.001345 |
| 792.52904 | | | | 12.39 | 26.28 | 0.002318 |
| 794.53745 | PC(PGJ2/P-16:0)(P) | PC(PGJ2/P-16:0) | 0 | 21.33 | 56.5 | 0.000309 |
| 795.53845 | SM(d36:2)(P) | SM(d18:1/18:1)-2OH(9,10)) | 5 | 12.25 | 35.06 | 0.001239 |
| 810.52957 | PE(P-40:6)(P) | PE(22:5(4Z,7Z,10Z,13Z,16Z)/P-18:1(11Z)) | 11 | 19.26 | 90.77 | 0.000071 |
| 811.53218 | PA(i-39:1)(P) | PA(18:1(12Z)-2OH(9,10)/i-21:0) | 7 | 4.074 | 28.44 | 0.000069 |
| 826.48811 | PC(36:7)(P) | PC(14:1(9Z)/22:6(5Z,7Z,10Z,13Z,16Z,19Z)-OH(4)) | 10 | 11.35 | 36.41 | 0.000309 |
| 834.52655 | PS(40:6)(P) | PS(20:4(8Z,11Z,14Z,17Z)/20:2(11Z,14Z)) | 3 | 6.631 | 21.69 | 0.00104 |
| 842.52341 | | | | 4.119 | 11.37 | 0.000132 |
| 847.73952 | | | | 12.09 | 0.9954 | 0.001349 |
| 858.51714 | PC(DiMe/22:5)(P) | PC(22:5(4Z,7Z,10Z,13Z,19Z)-O(16,17)/DiMe(9,3)) | 14 | 10.28 | 25.35 | 0.000712 |
| 873.75291 | PC(24:4)(P) | | | 18.07 | 2.219 | 0.000931 |
| 885.55072 | PI(38:4)(P) | PI(18:1(9Z)/20:3(5Z,8Z,11Z)) | 1 | 106.5 | 273.7 | 0.000246 |
| 886.55172 | PC(DiMe/22:6)(P) | PC(DiMe(9,5)/22:5(4Z,7Z,10Z,13Z,19Z)-O(16,17)) | 10 | 54.38 | 143.5 | 0.000208 |
| 162.11389 | L-carnitine | L-Carnitine | 9 | 259 | 968.9 | 0.003349 |
| 163.11851 | | | | 1.159 | 17.64 | 0.003564 |
| 184.09451 | L-Carnitine(P) | L-Carnitine | 1 | 37.8 | 213 | 0.003964 |
| 200.06846 | | | | 67.52 | 275.3 | 0.003564 |
| 218.14107 | Propionylcarnitine(P) | Propionylcarnitine | 11 | 64.06 | 300.6 | 0.003388 |
| 221.18648 | | | | 158.4 | 429.4 | 0.004688 |
| 351.25031 | 5a-Tetrahydrocorticosterone(P) | 5a-Tetrahydrocorticosterone | 8 | 328.6 | 17.36 | 0.002451 |

| | | | | | | |
|---|---|---|---|---|---|---|
| 367.22426 | MG(16:1)(P) | MG(16:1(9Z)/0:0/0:0) | 1 | 350.6 | 33.89 | 0.002347 |
| 379.28231 | MG(20:4)(P) | MG(20:4(8Z,11Z,14Z,17Z)/0:0/0:0) | 5 | 592.3 | 209.7 | 0.002451 |
| 380.27904 | LTB4 ethanolamide(P) | Leukotriene B4 ethanolamide | 11 | 87.62 | 25.73 | 0.002451 |
| 391.22464 | MG(18:3)(P) | MG(18:3(9Z,12Z,15Z)/0:0/0:0) | 0 | 29.87 | 8.094 | 0.002451 |
| 395.25626 | LysoPA(P-16:0)(P) | LysoPA(P-16:0/0:0) | 2 | 763.8 | 299.3 | 0.002474 |
| 396.25562 | | | | 173.2 | 64.48 | 0.002474 |
| 397.25498 | | | | 36.37 | 6.782 | 0.002451 |
| 411.25125 | DG(18:0)(P) | DG(8:0/10:0/0:0) | 1 | 58.03 | 16.56 | 0.002451 |
| 415.20924 | | | | 145.7 | 7.112 | 0.002474 |
| 439.20961 | | | | 143.8 | 30.38 | 0.002451 |
| 441.22674 | | | | 237.6 | 133.4 | 0.005634 |
| 443.24123 | | | | 392.4 | 135.7 | 0.003044 |
| 444.24322 | | | | 174.4 | 42.44 | 0.002451 |
| 521.32529 | | | | 359.4 | 5.185 | 0.004147 |
| 522.32728 | | | | 48.03 | 0.1579 | 0.006701 |
| 523.33979 | PA(23:0)(P) | PA(10:0/13:0) | 1 | 1172 | 193.5 | 0.002451 |
| 533.38071 | | | | 319.9 | 16.39 | 0.003564 |
| 549.35992 | | | | 351.7 | 37.16 | 0.002755 |
| 550.35928 | | | | 343.8 | 19.65 | 0.002474 |
| 560.31078 | LysoPC(18:1)(P) | PC(18:1(6Z)/0:0) | 1 | 100.4 | 21.11 | 0.002451 |
| 565.35228 | | | | 60.93 | 6.034 | 0.002451 |
| 575.37216 | | | | 35.63 | 1.92 | 0.004916 |
| 589.48152 | DG(34:4)(P) | DG(14:0/0:0/20:4n6) | 2 | 130.9 | 0.6817 | 0.008035 |
| 597.33963 | | | | 253.2 | 23.12 | 0.002451 |
| 603.44361 | DG(32:2)(P) | DG(16:1n7/0:0/16:1n7) | 8 | 92.45 | 6.827 | 0.009217 |
| 613.4819 | DG(36:6)(P) | DG(16:1n7/0:0/20:5n3) | 1 | 167.5 | 6.163 | 0.003964 |
| 617.51352 | DG(34:3)(P) | DG(14:0/20:3(5Z,8Z,11Z)/0:0) | 3 | 229.1 | 24.94 | 0.002345 |
| 618.51551 | Cer(t18:0/20:5)(P) | Cer(t18:0/20:5(6E,8Z,11Z,14Z,17Z)-OH(5)) | 10 | 178.1 | 10.2 | 0.002203 |
| 629.45585 | DG(34:3n11)(P) | DG(14:1n5/0:0/20:2n6) | 3 | 612.6 | 35.94 | 0.002474 |
| 630.45784 | Cer(d16:1/20:3)(P) | Cer(d16:1/20:3(8Z,11Z,14Z)-2OH(5,6)) | 13 | 121.6 | 4.223 | 0.002474 |
| 633.48747 | DG(34:1)(P) | DG(18:0/0:0/16:1n7) | 3 | 747.8 | 107 | 0.002203 |
| 634.48946 | LysoPC(26:1)(P) | LysoPC(26:1(5Z)/0:0) | 14 | 595.1 | 67.62 | 0.002203 |
| 635.49408 | DG(35:4)(P) | DG(20:4(8Z,11Z,14Z,17Z)-2OH(5S,6R)/0:0/i-15:0) | 1 | 27.82 | 3.367 | 0.002203 |
| 643.42845 | DG(i-34:4)(P) | DG(i-14:0/0:0/20:4(5Z,7E,11Z,14Z)-OH(9)) | 8 | 27.94 | 0.5011 | 0.006621 |
| 643.52839 | DG(38:5)(P) | DG(20:2n6/0:0/18:3n3) | 2 | 171.1 | 22.3 | 0.002347 |
| 644.53038 | Cer(t40:5)(P) | Cer(t18:0/22:5(4Z,7Z,10Z,13Z,19Z)-O(16,17)) | 9 | 131.1 | 9.777 | 0.002345 |
| 645.45084 | DG(i-34:3)(P) | DG(18:3(9,11,15)-OH(13)/0:0/i-16:0) | 3 | 493.8 | 46.54 | 0.003564 |

| | | | | | | |
|---|---|---|---|---|---|---|
| 646.45283 | Cer(d16:1)(P) | Cer(d16:1/PGD1) | 13 | 97.01 | 6.799 | 0.003564 |
| 647.46534 | DG(i-34:1)(P) | DG(18:1(9Z)-O(12,13)/0:0/i-16:0) | 1 | 390.7 | 48.57 | 0.003964 |
| 648.46995 | | | | 142.8 | 12.19 | 0.003964 |
| 657.48522 | DG(36:3)(P) | DG(16:0/20:3(11Z,14Z,17Z)/0:0) | 0 | 1709 | 459.1 | 0.002474 |
| 658.40568 | | | | 27.73 | 6.539 | 0.002474 |
| 658.48983 | Cer(t18:0/20:4)(P) | Cer(t18:0/20:4(5Z,7E,11Z,14Z)-OH(9)) | 14 | 726 | 183.7 | 0.002474 |
| 659.50234 | DG(36:2)(P) | DG(18:1n9/0:0/18:1n9) | 2 | 584.9 | 97.86 | 0.002203 |
| 660.50696 | | | | 245.7 | 33.41 | 0.002203 |
| 661.4432 | DG(i-34:3)(P) | DG(20:3(8Z,11Z,14Z)-2OH(5,6)/0:0/i-14:0) | 1 | 112.8 | 4.947 | 0.004147 |
| 661.50895 | DG(36:1)(P) | DG(22:0/0:0/14:1n5) | 12 | 33.63 | 2.176 | 0.002203 |
| 662.44782 | Cer(d40:8)(P) | Cer(d18:2(4E,14Z)/22:5(4Z,7Z,10Z,13Z,19Z)-O(16,17)) | 10 | 37.82 | 0.6588 | 0.006576 |
| 673.48021 | DG(i-36:3)(P) | DG(18:3(9,11,15)-OH(13)/0:0/i-18:0) | 0 | 417 | 48.68 | 0.003279 |
| 674.37963 | | | | 125.3 | 32.56 | 0.003249 |
| 674.48482 | Cer(t38:4)(P) | Cer(t18:0/20:4(8Z,11Z,14Z,17Z)-2OH(5S,6R)) | 14 | 161.9 | 12.41 | 0.003249 |
| 679.45532 | PA(i-32:1)(P) | PA(18:1(12Z)-2OH(9,10)/i-14:0) | 1 | 516.2 | 51.75 | 0.002451 |
| 680.45993 | Cer(d40:7)(P) | Cer(d18:1/22:6(4Z,7Z,11E,13Z,15E,19Z)-2OH(10S,17)) | 8 | 205.1 | 15.8 | 0.002451 |
| 681.46455 | PG(i-29:0)(P) | PG(i-14:0/i-15:0) | 8 | 205.6 | 17.24 | 0.002203 |
| 682.47443 | Cer(t40:6)(P) | Cer(t18:0/22:5(4Z,7Z,10Z,13Z,19Z)-O(16,17)) | 9 | 392 | 16.11 | 0.002203 |
| 683.47379 | | | | 34.64 | 0.4178 | 0.002451 |
| 705.46493 | DG(i-16:0/0:0/PGD1)(P) | DG(5-iso PGF2VI/0:0/i-18:0) | 3 | 447.9 | 68.34 | 0.002451 |
| 706.47218 | PS(30:1)(P) | PS(16:1(9Z)/14:0) | 13 | 283.7 | 36 | 0.002451 |
| 707.48994 | DG(PGF1alpha/0:0/i-16:0)(P) | DG(PGF1alpha/0:0/i-16:0) | 6 | 319.5 | 30.81 | 0.002203 |
| 708.49193 | Cer(d20:1/22:6) | Cer(d20:1/22:6(4Z,7Z,11E,13Z,15E,19Z)-2OH(10S,17)) | 6 | 195.9 | 9.771 | 0.002203 |
| 709.49655 | PG(i-14:0/i-17:0)(P) | PG(i-14:0/i-17:0) | 7 | 23 | 0.7987 | 0.002451 |
| 719.45331 | PG(i-31:3)(P) | PG(18:3(9,11,15)-OH(13)/i-13:0) | 5 | 61.21 | 5.565 | 0.006969 |
| 754.53604 | PE-NMe2(35:4)(P) | PE-NMe2(20:4(8Z,11Z,14Z,17Z)/15:0) | 3 | 186.7 | 17.41 | 0.002451 |
| 755.54066 | SM(d35:1)(P) | SM(d19:1/16:0) | 8 | 72.84 | 2.346 | 0.002474 |
| 759.57491 | DG(PGD1/0:0/i-21:0)(P) | DG(PGD1/0:0/i-21:0) | 0 | 6.355 | 49.33 | 0.004688 |
| 770.51 | PE-NMe2(33:1)(P) | PE-NMe2(18:1(9Z)/15:0) | 0 | 378.8 | 52.84 | 0.003249 |
| 771.51461 | DG(41:5)(P) | DG(20:5(7Z,9Z,11E,13E,17Z)-3OH(5,6,15)/0:0/i-21:0) | 4 | 167.2 | 14.16 | 0.002735 |
| 786.60229 | PC(36:2) | PC(18:1(9Z)-O(12,13)/P-18:0) | 2 | 13.51 | 112.5 | 0.006722 |
| 802.52628 | PS(36:2)(P) | PS(18:1(9Z)-O(12,13)/18:1(9Z)) | 4 | 28.55 | 1.329 | 0.002347 |
| 818.50023 | DG(LTE4/0:0/i-18:0)(P) | DG(LTE4/0:0/18:0) | 1 | 78.96 | 14.09 | 0.002451 |
| 831.55493 | PG(i-37:0)(P) | PG(i-13:0/i-24:0) | 4 | 95.13 | 37.37 | 0.003044 |
| 861.66987 | | | | 63.42 | 138.4 | 0.00735 |

| | | | | | | |
|---|---|---|---|---|---|---|
| 871.71342 | TG(50:1)(P) | TG(14:0/18:1(9Z)/18:0) | 2 | 290.1 | 19.67 | 0.002451 |
| 872.71804 | PC(42:1)(P) | PC(24:1(15Z)/18:0) | 9 | 139.9 | 6.056 | 0.002451 |
| 897.72829 | TG(52:2)(P) | TG(14:0/18:1(9Z)/20:1(11Z)) | 0 | 768.6 | 91.46 | 0.002474 |
| 898.73291 | PE-NMe(46:2)(P) | PE-NMe(24:1(15Z)/22:1(13Z)) | 10 | 416.5 | 41.96 | 0.002451 |
| 899.73753 | TG(52:1)(P) | TG(14:0/24:0/14:1(9Z)) | 11 | 166.5 | 13.2 | 0.002451 |

**Supplementary Table 7.** The *m/z* and full compound name for molecules found to be significantly changed in the liver of DSS-treated mice. Table includes the mean relative abundance value for each molecule within the groups. *P*-values highlighted in grey are for molecules decreased in DSS-treated mice compared to controls and *p* values not highlighted show molecules increased in the treated group compared to controls.

**Supplementary Table 8**

| *m/z* | VIP list |
|---|---|
| 656.5120239 | 1.208789945 |
| 489.3439941 | 1.190179944 |
| 842.5189819 | 1.18987 |
| 364.2340088 | 1.189360023 |
| 558.3330078 | 1.188580036 |
| 363.2309875 | 1.186579943 |
| 655.5079956 | 1.185269952 |
| 591.5360107 | 1.184820056 |
| 509.4580078 | 1.181589961 |
| 510.4609985 | 1.181229949 |
| 433.2810059 | 1.180879951 |
| 300.0400085 | 1.178550005 |
| 462.3150024 | 1.176789999 |
| 461.3129883 | 1.176280022 |
| 584.4970093 | 1.17445004 |
| 485.3129883 | 1.17227006 |
| 276.2049866 | 1.171900034 |
| 611.526001 | 1.171239972 |
| 583.4949951 | 1.171090007 |
| 810.5300293 | 1.170300007 |
| 180.0599976 | 1.170050025 |
| 811.5319824 | 1.169929981 |
| 612.5289917 | 1.16989994 |
| 365.2269897 | 1.169819951 |
| 533.4550171 | 1.165969968 |
| 537.4899902 | 1.165949941 |
| 538.4920044 | 1.165410042 |
| 436.2999878 | 1.165019989 |
| 254.220993 | 1.164790034 |
| 253.2169952 | 1.164330006 |
| 826.4879761 | 1.164319992 |
| 630.4949951 | 1.16407001 |
| 435.29599 | 1.163190007 |
| 637.5410156 | 1.160719991 |
| 309.2789917 | 1.160339952 |
| 638.5449829 | 1.158370018 |
| 773.5339966 | 1.155329943 |
| 434.2839966 | 1.155169964 |
| 310.2839966 | 1.154590011 |
| 267.2340088 | 1.152150035 |
| 534.460022 | 1.151170015 |
| 316.2130127 | 1.150329947 |
| 552.2969971 | 1.149690032 |
| 337.3110046 | 1.149279952 |
| 607.4949951 | 1.148560047 |
| 477.3059998 | 1.147539973 |
| 613.5360107 | 1.147359967 |
| 886.552002 | 1.144719958 |
| 281.2479858 | 1.144649982 |

| | |
|---|---|
| 282.2520142 | 1.144119978 |
| 608.4970093 | 1.143650055 |
| 702.3969727 | 1.141000032 |
| 794.5339966 | 1.14064002 |
| 885.5510254 | 1.140020013 |
| 572.4819946 | 1.134539962 |
| 268.2349854 | 1.13409996 |
| 275.2000122 | 1.133659959 |
| 858.5170288 | 1.130879998 |
| 283.256012 | 1.129999995 |
| 657.4140015 | 1.127689958 |
| 634.4890137 | 1.127339959 |
| 633.4869995 | 1.125460029 |
| 771.5180054 | 1.12451005 |
| 531.440979 | 1.124500036 |
| 315.2090149 | 1.122720003 |
| 280.4689941 | 1.122689962 |
| 708.4920044 | 1.122099996 |
| 179.0559998 | 1.121639967 |
| 774.5360107 | 1.121209979 |
| 707.4899902 | 1.120239973 |
| 281.098999 | 1.118800044 |
| 281.3999939 | 1.11864996 |
| 284.2590027 | 1.118350029 |
| 573.4840088 | 1.118070006 |
| 545.3619995 | 1.115869999 |
| 635.4940186 | 1.113720059 |
| 601.4609985 | 1.113649964 |
| 512.4769897 | 1.113309979 |
| 639.4849854 | 1.112759948 |
| 441.2780151 | 1.112040043 |
| 280.2369995 | 1.110820055 |
| 279.2330017 | 1.110739946 |
| 511.4729919 | 1.109990001 |
| 681.4650269 | 1.109109998 |
| 618.5159912 | 1.106960058 |
| 660.507019 | 1.104900002 |
| 790.5029907 | 1.103520036 |
| 563.5050049 | 1.103170037 |
| 564.507019 | 1.102710009 |
| 525.3189697 | 1.101979971 |
| 507.4419861 | 1.101600051 |
| 659.5020142 | 1.101140022 |
| 661.5089722 | 1.10089004 |
| 682.473999 | 1.099670053 |
| 644.5300293 | 1.09849 |
| 617.5139771 | 1.097959995 |
| 795.5380249 | 1.097309947 |
| 559.4719849 | 1.096490026 |
| 367.223999 | 1.095010042 |
| 560.4760132 | 1.092620015 |

| | |
|---|---|
| 629.492981 | 1.092370033 |
| 643.5280151 | 1.09205997 |
| 626.5360107 | 1.092000008 |
| 339.2120056 | 1.09149003 |
| 802.526001 | 1.091429949 |
| 592.3259888 | 1.090680003 |
| 834.5269775 | 1.088619947 |
| 513.4799805 | 1.087520003 |
| 268.2000122 | 1.08739996 |
| 706.4719849 | 1.087170005 |
| 417.2860107 | 1.087069988 |
| 265.2170105 | 1.08573997 |
| 705.4650269 | 1.084959984 |
| 351.25 | 1.084939957 |
| 831.5549927 | 1.083950043 |
| 597.3400269 | 1.083539963 |
| 574.4790039 | 1.081650019 |
| 524.3439941 | 1.081590056 |
| 899.7379761 | 1.077250004 |
| 818.5 | 1.076810002 |
| 638.3309937 | 1.076629996 |
| 754.5360107 | 1.076390028 |
| 161.0460052 | 1.07565999 |
| 391.2250061 | 1.07562995 |
| 523.3400269 | 1.07506001 |
| 380.2789917 | 1.074820042 |
| 680.460022 | 1.073089957 |
| 872.7180176 | 1.072639942 |
| 550.3590088 | 1.071429968 |
| 565.3519897 | 1.071179986 |
| 679.4550171 | 1.071099997 |
| 439.2099915 | 1.070899963 |
| 415.2090149 | 1.069949985 |
| 682.5930176 | 1.069910049 |
| 709.4970093 | 1.069370031 |
| 267.1960144 | 1.068809986 |
| 581.4780273 | 1.06851995 |
| 560.3109741 | 1.067569971 |
| 630.4580078 | 1.06704998 |
| 397.2550049 | 1.067029953 |
| 595.3449707 | 1.066359997 |
| 648.3499756 | 1.066249967 |
| 549.3599854 | 1.065389991 |
| 128.0359955 | 1.065250039 |
| 791.5059814 | 1.065029979 |
| 629.4559937 | 1.064450026 |
| 444.2430115 | 1.064059973 |
| 873.7529907 | 1.063189983 |
| 411.2510071 | 1.062909961 |
| 871.7130127 | 1.062849998 |
| 379.2820129 | 1.062070012 |

| | |
|---|---|
| 351.1820068 | 1.061429977 |
| 683.5939941 | 1.060950041 |
| 583.473999 | 1.06020999 |

**Supplementary Table 8.** PLS-DA generated VIP>1 list of molecules in the liver that contribute towards group separation in DSS-treated mice compared to controls.

**Supplementary Table 9**

| *m/z* | Heatmap ID | compound_name | adduct | ppm | Mean 0% | Mean 3% | p-value |
|---|---|---|---|---|---|---|---|
| 117.0558 | 5-HPA(P) | 5-Hydroxypentanoic acid | M-H | 10 | 76.95 | 154 | 0.0118 |
| 118.08744 | Betaine(P) | Betaine | M+H | 5 | 124.5 | 247.1 | 0.0036 |
| 124.0152 | m/z 124.0152 | | | | 4220 | 3031 | 0.0499 |
| 134.0473 | m/z 134.0473 | | | | 88.11 | 166.9 | 0.0795 |
| 156.04378 | 3-AMBA | 3-Amino-3-methylbutanoic acid | M+K | 11 | 1235 | 2173 | 0.0002 |
| 157.04563 | m/z 157.0456 | | | | 45.43 | 103 | 7E-05 |
| 158.04035 | 2-DHBA(P) | (+)-threo-2-Amino-3,4-dihydroxybutanoic acid | M+Na | 13 | 39.42 | 75.59 | 0.0001 |
| 160.13189 | 5-AVAB(P) | 5-amino valeric acid betaine | M+H | 8 | 1775 | 1083 | 0.0287 |
| 161.13849 | m/z 161.1384 | | | | 143.7 | 82.82 | 0.0299 |
| 166.018 | Quinolinic acid(P) | Quinolinic acid | M-H | 11 | 656.3 | 431.6 | 0.0503 |
| 174.0409 | NFLGA(P) | N-Formyl-L-glutamic acid | M-H | 1 | 217.9 | 118.5 | 0.0369 |
| 183.1391 | m/z 183.1391 | | | | 126.9 | 200.7 | 0.0051 |
| 187.0976 | m/z 187.0976 | | | | 375.3 | 550.1 | 0.0665 |
| 188.05581 | NALGA(P) | N-Acetyl-L-glutamic acid | M-H | 3 | 0.3565 | 19.28 | 0.0251 |
| 211.134 | m/z 211.134 | | | | 171.9 | 245.4 | 0.0046 |
| 221.18775 | m/z 221.18775 | | | | 294.1 | 783.7 | 0.0001 |
| 222.1896 | m/z 222.1896 | | | | 13.99 | 70.27 | 0.0002 |
| 227.129 | m/z 227.129 | | | | 112.5 | 178 | 0.0129 |
| 229.1962 | m/z 229.1962 | | | | 17.8 | 32.71 | 5E-05 |
| 249.1859 | m/z 249.1859 | | | | 11 | 26.1 | 0.029 |
| 251.2016 | m/z 251.2016 | | | | 36.05 | 56.97 | 0.0091 |
| 254.2207 | m/z 254.2207 | | | | 300.8 | 192.9 | 0.0276 |
| 256.0595 | N-AMA(P) | N-Acetylmannosamine | M+Cl | 0 | 5.014 | 33.23 | 0.011 |
| 263.0408 | m/z 263.0408 | | | | 112.4 | 52.39 | 0.0131 |
| 273.1861 | ODT acid(P) | (10Z,14E,16E)-10,14,16-Octadecatrien-12-ynoic acid | M-H | 3 | 201.2 | 311.4 | 0.0003 |
| 274.1896 | m/z 274.1896 | | | | 30.6 | 55.33 | 7E-05 |
| 277.0597 | m/z 277.0597 | | | | 41.15 | 95.22 | 0.0122 |
| 279.0562 | Biotin(P) | Biotin | M+Cl | 7 | 7.915 | 28.21 | 0.0101 |
| 279.233 | DHL acid(P) | Dihomolinoleic acid | M-H | 0 | 6841 | 8970 | 0.0351 |
| 280.2363 | Petroselinate(P) | petroselinate | M-H | 12 | 876.3 | 1151 | 0.0348 |
| 281.0536 | m/z 281.0536 | | | | 50.53 | 22.8 | 0.0223 |
| 301.1811 | Hydroxy steroid (P) | 11b-Hydroxyandrost-4-ene-3,17-dione | M-H | 1 | 8.653 | 40.3 | 0.0491 |
| 317.1759 | Ubiquinone-2(P) | Ubiquinone-2 | M-H | 1 | 30.77 | 103.8 | 0.0055 |

| | | | | | | | |
|---|---|---|---|---|---|---|---|
| 318.1737 | m/z 318.1737 | | | | 1.079 | 15.98 | 0.0198 |
| 327.2331 | DHA(P) | Docosahexaenoicacid(DHA) | M-H | 1 | 1236 | 3224 | 0.0704 |
| 328.2364 | m/z 328.2364 | | | | 290.7 | 763.3 | 0.0391 |
| 329.2335 | 9,10,13-TriHOME(P) | 9,10,13-TriHOME | M-H | 9 | 20.58 | 59.11 | 0.0744 |
| 332.2267 | m/z 332.2267 | | | | 38.38 | 12.67 | 0.0126 |
| 341.2139 | 14-oxo-DoHE(1-)(P) | 14-oxo-DoHE(1-) | M-H | 5 | 7.892 | 73.42 | 0.0618 |
| 343.228 | EDP acid(P) | Epoxydocosapentaenoic acid | M-H | 1 | 89.45 | 212 | 0.0016 |
| 344.2302 | m/z 344.2302 | | | | 22.08 | 71.88 | 0.0016 |
| 359.223 | Resolvin D5(P) | Resolvin D5 | M-H | 1 | 32.96 | 75.22 | 0.0058 |
| 363.2099 | m/z 363.2099 | | | | 56.1 | 142.1 | 0.0156 |
| 363.2309 | MG(16:1) | MG(16:1(9Z)/0:0/0:0) | M+Cl | 2 | 54.54 | 15.94 | 0.0085 |
| 365.2097 | 9,12,13-TriHOME(P) | 9,12,13-TriHOME | M+Cl | 1 | 11.24 | 42.09 | 0.0089 |
| 373.2019 | m/z 373.2019 | | | | 17.21 | 82.12 | 0.0324 |
| 375.2171 | Resolvin D1(P) | Resolvin D1 | M-H | 1 | 27.97 | 63.26 | 0.0062 |
| 413.2464 | Nor-DCA(P) | Nordeoxycholic acid | M+Cl | 0 | 444.4 | 200.6 | 0.0444 |
| 414.2498 | m/z 414.2498 | | | | 111.5 | 49.1 | 0.0435 |
| 415.2339 | MG(20:3)(P) | MG(20:3(8Z,11Z,14Z)/0:0/0:0) | M+Cl | 0 | 211.4 | 102.2 | 0.0505 |
| 415.2711 | m/z 415.2711 | | | | 42.05 | 15.19 | 0.0341 |
| 416.2374 | m/z 416.2374 | | | | 34.01 | 13.04 | 0.0425 |
| 439.2626 | MG(22:5)(P) | MG(22:5(4Z,7Z,10Z,13Z,16Z)/0:0/0:0) | M+Cl | 1 | 75.25 | 29.07 | 0.0448 |
| 467.2988 | m/z 467.2988 | | | | 70.7 | 27.08 | 0.02 |
| 558.3332 | LysoPC(18:0)(P) | LysoPC(0:0/18:0) | M+Cl | 1 | 25.31 | 52.76 | 0.016 |
| 611.5255 | m/z 611.5255 | | | | 112.5 | 70.13 | 0.0179 |
| 736.6459 | m/z 736.6459 | | | | 189.5 | 110.5 | 0.0423 |
| 737.6491 | m/z 737.6491 | | | | 90.76 | 51.31 | 0.0406 |
| 742.57217 | PC(O-34:3)(P) | PC(O-16:1(9Z)/18:2(9Z,12Z)) | M+H | 3 | 141.7 | 61.65 | 0.0057 |
| 750.5431 | PE(P-38:4)(P) | PE(P-18:1(9Z)/20:3(5Z,8Z,11Z)) | M-H | 4 | 161.7 | 96.69 | 0.0441 |
| 758.54479 | m/z 758.5448 | | | | 269.3 | 100.9 | 0.0075 |
| 768.58703 | PC(P-34:0)(P) | PC(P-18:0/16:0) | M+Na | 1 | 71.58 | 37.78 | 0.0107 |
| 784.55964 | PC(O-34:1)(P) | PC(O-18:1(9Z)/16:0) | M+K | 3 | 210.3 | 113.4 | 0.0067 |
| 788.5455 | PS(36:1)(P) | PS(22:1(13Z)/14:0) | M-H | 1 | 136 | 56.33 | 0.0105 |
| 789.5492 | SM(d40:6)(P) | SM(d18:1/22:5(4Z,7Z,10Z,13Z,19Z)-O(16,17)) | M-H | 7 | 83.55 | 28 | 0.004 |

**Supplementary Table 9.** The *m/z*, abbreviated names shown in the heatmap in Fig. 6, and full compound name for molecules found to be significantly changed in the spleen of DSS-treated mice. The table includes the mean relative abundance value for each molecule within the groups. *P*-values highlighted in grey are for molecules down regulated in 3% DSS-treated mice compared to controls and *p*-values not highlighted show molecules increased in the treatment group compared to the control group.

**Supplementary Table 10**

| *m/z* | VIP list |
|---|---|
| 273.1870117 | 1.278789997 |
| 274.1889954 | 1.276319981 |
| 157.0460052 | 1.264089942 |
| 229.197998 | 1.253790021 |
| 158.0399933 | 1.252120018 |
| 156.0440063 | 1.244019985 |
| 221.1880035 | 1.221740007 |
| 222.1900024 | 1.205440044 |
| 558.3330078 | 1.138569951 |
| 343.2279968 | 1.112179995 |
| 758.5449829 | 1.111770034 |
| 344.230011 | 1.111410022 |
| 784.5599976 | 1.087800026 |
| 742.5720215 | 1.077800035 |
| 183.1390076 | 1.074849963 |
| 789.5490112 | 1.067000031 |
| 118.086998 | 1.057729959 |
| 359.2219849 | 1.044929981 |
| 611.526001 | 1.042279959 |
| 317.1759949 | 1.041479945 |
| 227.128006 | 1.039909959 |
| 211.1329956 | 1.039870024 |
| 251.201004 | 1.027750015 |
| 277.0610046 | 1.023830056 |
| 788.5449829 | 1.023040056 |
| 375.2170105 | 1.020480037 |
| 279.0559998 | 1.016469955 |
| 215.072998 | 1.013659954 |
| 768.5869751 | 1.008110046 |
| 332.2260132 | 1.004760027 |
| 365.2099915 | 1.003100038 |

**Supplementary Table 10.** PLS-DA generated VIP>1 list of molecules in the spleen that contribute towards group separation in the DSS colitis model.

**Supplementary Table 11**

| *m/z* | Heatmap ID | compound_name | adduct | ppm | Mean 0% | Mean 3% | p-value |
|---|---|---|---|---|---|---|---|
| 257.176 | TDA(P) | Tetradecanedioic acid | M-H | 1 | 99.36 | 10.26 | 0.018893 |
| 277.217 | GLA(P) | gamma-Linolenic acid | M-H | 5 | 699.4 | 344.5 | 0.016252 |
| 278.21 | | | | | 129.2 | 57.14 | 0.015677 |
| 305.249 | DGLA(P) | Dihomo-gamma-linolenic acid | M-H | 4 | 620 | 362.7 | 0.010765 |
| 306.243 | *m/z* 306.2433 | | | | 130.8 | 72.03 | 0.007959 |
| 317.183 | *m/z* 317.183 | | | | 24.04 | 87.71 | 0.042873 |
| 363.248 | *m/z* 363.2475 | | | | 104.7 | 19.77 | 0.003618 |
| 367.27 | *m/z* 367.2697 | | | | 199.1 | 111.3 | 0.01858 |
| 603.477 | DG(32:0)(P) | DG(19:0/13:0/0:0) | M+Cl | 5 | 49.23 | 9.139 | 0.001994 |
| 754.537 | PC(34:4)(P) | PC(20:4(8Z,11Z,14Z,17Z)/14:0) | M+H | 0 | 49.34 | 26.78 | 0.00601 |
| 770.512 | PC(32:1)(P) | PC(18:1(9Z)/14:0) | M+K | 4 | 81.58 | 47.07 | 0.025015 |
| 771.516 | PG(36:4)(P) | PG(20:4(5Z,8Z,11Z,14Z)/16:0) | M+H | 2 | 34.46 | 16.62 | 0.009635 |
| 772.528 | PE(35:0)(P) | PE(20:0/15:0) | M+K | 4 | 1123 | 721 | 0.110219 |
| 773.531 | PG(36:3)(P) | PG(18:3(9Z,12Z,15Z)/18:0) | M+H | 3 | 495.8 | 318.6 | 0.107909 |
| 817.501 | PG(40:8)(P) | PG(20:4(8Z,11Z,14Z,17Z)/20:4(8Z,11Z,14Z,17Z)) | M-H | 1 | 16.03 | 149.8 | 0.081714 |
| 865.502 | PG(38:4)(P) | PG(20:4(6Z,8E,10E,14Z)-2OH(5S,12R)/18:0) | M+Cl | 0 | 42.29 | 615.5 | 0.025988 |

**Supplementary Table 11.** The *m/z*, abbreviated names shown in heatmap in Fig. 7, and full compound name for molecules found to be significantly changed in the kidney of DSS-treated mice. Table includes the mean relative abundance value for each molecule within the groups. *P*-values highlighted in grey are for molecules decreased in 3% DSS treated mice compared to controls and *p*-values not highlighted show molecules increased in the treatment group compared to controls.

**Supplementary Table 12**

| *m/z* | VIP list |
|---|---|
| 754.5380249 | 1.343119979 |
| 771.5150146 | 1.293339968 |
| 770.5130005 | 1.258380055 |
| 772.5280151 | 1.05061996 |
| 773.5310059 | 1.049360037 |

**Supplementary Table 12.** PLS-DA generated VIP>1 list of molecules in the colon that contribute towards group separation in the DSS colitis model.
